# Characterization of root exudates of black oat in the presence of interspecific weed species neighbours and intraspecific neighbours, and their effects on root traits

**DOI:** 10.1101/2024.09.02.610795

**Authors:** Çağla G. Eroğlu, Alexandra A. Bennett, Teresa Steininger-Mairinger, Stephan Hann, Markus Puschenreiter, Judith Wirth, Aurélie Gfeller

## Abstract

Root exudates are composed of primary and secondary organic compounds that function as signalling molecules and play important roles in plant-environment interactions. The quantity and composition of root exudates vary depending on the species, genotype, and environmental conditions, including the presence and identity of neighbouring plants. Although cover crops are commonly used in agricultural practices for their ecosystem services, such as weed suppression, pathogen control, and soil structure improvement, studies on their root exudates are limited. Our study provides the first characterization of the root exudates of black oat interacting with weed neighbours, redroot pigweed and blackgrass, as well as with neighbours of the same species, black oat. We investigated how these interactions influence the black oat root exudation patterns. Furthermore, we investigated the impact of black oat presence on neighbours and how exposure to black oat root exudate treatments affects the root traits of weeds. The upregulated compounds detected in root exudates in response to neighbouring plants primarily belonged to the organic oxygen compounds superclass, with most of which identified as amino acids and carbohydrates. In the presence of redroot pigweed, a general increase in root exudation was observed, with amino acids and sugar sulphates being upregulated. The presence of black oat had varying effects among the neighbouring plants. While significant decreases observed and black oat in redroot pigweed root traits, an increase was observed in blackgrass. Similarly, more pronounced effects were observed in redroot pigweed compared to blackgrass upon root exudate application. This study provides a characterization and insights into the dynamic nature of root exudates and their influence on root traits and plant interactions.

## 1. INTRODUCTION

Root exudates consist of organic compounds that belong to primary metabolism, such as sugars, amino acids, and organic acids, as well as compounds that belong to secondary metabolism, such as phenolics, terpenoids, and alkaloids (Badri & Vivanco, 2009; Maurer et al., 2021; Zhalnina et al., 2018). They act as signalling molecules and play vital roles in plant-plant and plant-microbiome interactions by shaping soil physicochemical properties and microbial communities (Alahmad et al., 2024; Mondal et al., 2023).

Root exudates, which contain naturally occurring compounds called allelochemicals, impact other organisms in their vicinity. These plant-derived allelochemicals can have suppressive effects (e.g., causing nutrient depletion, hindering metabolic processes, and inhibiting germination, growth, and development) or promoting impacts (e.g., making certain nutrients available, promoting beneficial microbial communities, and providing growth-stimulating compounds) on neighbouring plants (Bais et al., 2006; Inderjit & Duke, 2003). The root exudates released by different plants in a shared environment affect one another, resulting in changes in the composition of root exudates each plant releases. The quantity and composition of root exudates vary depending on many factors such as plant species, genotype, and environmental conditions. The family and species of plants are significant sources of variation in root exudation and their fate due to different root system architecture and the presence of microbial communities (Gunjal, 2023; Mondal et al., 2023; Upadhyay et al., 2022). The release of compounds from actively growing root cells regulates the levels of primary metabolites in the root tip over time. Sugars, amino acids, and organic acids which can be found in root exudates in high concentrations are capable of triggering different responses in root system architecture from stimulating lateral root development and elongation to root meristem depletion which facilitates resource uptake (Canarini et al., 2019; Malamy & Ryan, 2001)

The root exudates of agriculturally important crops such as maize, rice, and wheat (Aulakh et al., 2001; Bacilio-Jiménez et al., 2003; Hu et al., 2018; Oburger et al., 2014; Santangeli et al., 2024; Tawaraya et al., 2018), as well as model plants such as *Arabidopsis thaliana* and *Brachypodium distachyon* (Chaparro et al., 2013; Kawasaki et al., 2016; McLaughlin et al., 2023; Mönchgesang et al., 2016; Sharma et al., 2020), have been studied mostly in the frame of microbiome interactions, stress responses, or variations among different varieties or cultivars. Additionally, root exudates of cover crops which provides multiple ecosystem benefits such as buckwheat (Eroğlu et al., 2024; Gfeller et al., 2018; Kalinova et al., 2007; Szwed et al., 2019; Vieites-Álvarez et al., 2023), sorghum (Ayoodunfa, 1978; Dayan et al., 2003; Erickson et al., 2001; Seitz et al., 2024; P. Wang et al., 2021), hairy vetch (Mardani-Korrani et al., 2021; Seitz et al., 2024), brassica species (Gargallo-Garriga et al., 2018; Li et al., 2021; Wadhwa & Narula, 2012), and rye (Hazrati et al., 2020; Pérez & Ormenoñuñez, 1991; Seitz et al., 2024) have been analysed to investigate their influence on soil health and microbiome and plant-plant interactions. However, despite being a widely used cover crop due to its positive impact on agricultural practices through suppressing weeds and improving soil structure (Marquez et al., 2022), studies on the root exudates of black oat are lacking. Black oat is a winter annual belonging to the Poaceae family and originates from the Mediterranean region (Dial, 2014). Even though there are studies on other Poaceae species, the root exudate composition of black oat has remained unknown so far.

Due to the unknown nature of the exudates of interest, a non-targeted analysis (NTA) chemometrics approach can provide information to help prioritize and begin annotating compounds within the rhizosphere (Gertsman & Barshop, 2018). Prioritization helps highlight compounds of interest through statistical approaches, such as significance testing (Kumar et al., 2018) and dimension reduction techniques. Additionally, techniques which relate compounds to one another based upon their fragmentation patterns (Csermely et al., 2013), such as molecular networking, can further assist with prioritization. Chemical relatedness can be inferred during molecular networking as compounds with many shared fragments and resulting strong similarity scores are more likely to be structurally related (M. Wang et al., 2016). Identification of these prioritized compounds can then be performed based upon their m/z, isotope patterns, and fragmentation patterns. Identification confidence levels can vary from level 1 (a confirmed structure based upon a reference standard) to level 3 (a tentative candidate in which the m/z and MS^2^ information corroborate a structure, but there is not enough other information to be sure of the exact structure) and lower (Schymanski et al., 2014).

Here, we investigated the impact of the presence of interspecific weed neighbours, redroot pigweed (*Amaranthus retroflexus* L.) and black grass (*Alopecurus myosuroides* Huds.), and intraspecific neighbours on the root exudation patterns and root traits of black oat (*Avena strigosa*). Furthermore, we assessed how black oat, in turn, impacts weeds root traits. We also examined the application of Black oat root exudates on the root traits of the two weeds. To achieve the goals of this study, previously established split root systems (Eroğlu et al., 2024) were utilized to grow plants and collect root exudates. Furthermore, a dual extraction method was incorporated to obtain exudates from the same system at different time points: after two weeks and after 24 hours (Bennett et al., 2024). Combining these two methods to assess black oat allows for novel insights into root exudation and the impact of black oat presence and root exudates on the root traits of neighbours, both from the same species and different weed species, with the goal of laying the foundation for potential implications in weed management.

## 2. MATERIAL AND METHODS

### 2.1. Plant material and growth conditions

Two Whatman filter papers were placed inside 120 x 15 mm square Petri dishes (Corning® Gosselin™) and moistened by adding half-strength Hoagland’s solution (Hoagland’s No. 2 Basal Salt Mixture, Sigma-Aldrich) with a pH of 5.8. Five black oat (BO) (*Avena strigosa*, variety: Altesse) seeds were placed horizontally in a line on the filter papers. Half-sized blotting papers were placed over the line of BO seeds. The Petri dishes were sealed using parafilm, placed on a stand at a 90° angle, and positioned inside a phytotron (Aralab, Clitec) at 25/20°C (day/night), with a relative humidity of 70%. For the first three days, Petri dishes were kept in the dark; on the fourth day, they were exposed to light with a photosynthetic flux of 200 μmol m^-2^ s^-1^ and a 16/8-hour light/dark photoperiod; on the fifth day, they were transferred to split root systems.

### 2.2. Split-root system preparation

Solid phase extraction (SPE) cartridges, 60 mL (Bond Elut 12131018, Agilent) were used to prepare split root systems. The black contact paper was used to cover the outer surface of the SPE cartridges. Two cartridges were then affixed together using double-sided mounting tape, and the joining parts at the top were melted from the middle to create a small space to place the seedling. All the cartridges were filled with 250-400 µm glass beads (Guyson SA) and were moistened with half-strength Hoagland’s solution (Sigma Aldrich). The roots of each BO seedlings were counted, divided into two equal parts, and carefully taken out of Petri dishes using forceps. Equally split roots were placed in each SPE cartridge of the split root system. Across the two experimental setups there were four different split root conditions and four different single cartridge conditions. The first set of split root conditions included split root black oat without a neighbour (BO-0), with a black oat neighbour (BO-BO), with a redroot pigweed neighbour (BO-P) (n=5) and called as “P experimental set”. The second set of split root conditions included split root black oat without a neighbour (BO-0), with a black oat neighbour (BO-BO), and with a blackgrass neighbour (BO-G) (n=5) and were the “G experimental set” samples. In each split root condition, BO plants had direct root-root contact with the neighbour in one cartridge (cartridge B), while there was no contact in the other (cartridge A) (Figure 1) Single cartridges contained plants from each species: black oat (BO), redroot pigweed (P), and blackgrass (G) growing alone. All plants from both sets were grown in a phytotron under controlled conditions (16/8-hour light/dark cycle at 25/20°C, with a photoperiod and photosynthetic flux of 200 μmol m^-2^ s^-1^, and a relative humidity of 70%). Each cartridge received 5 ml of half-strength Hoagland’s solution every other day. Root exudate extraction was performed on the fourteenth day. The entire experiment was repeated three times, each with a one-week interval between repetitions, using five biological replicates per condition for each repetition (Figure 1).

**Figure 1:**
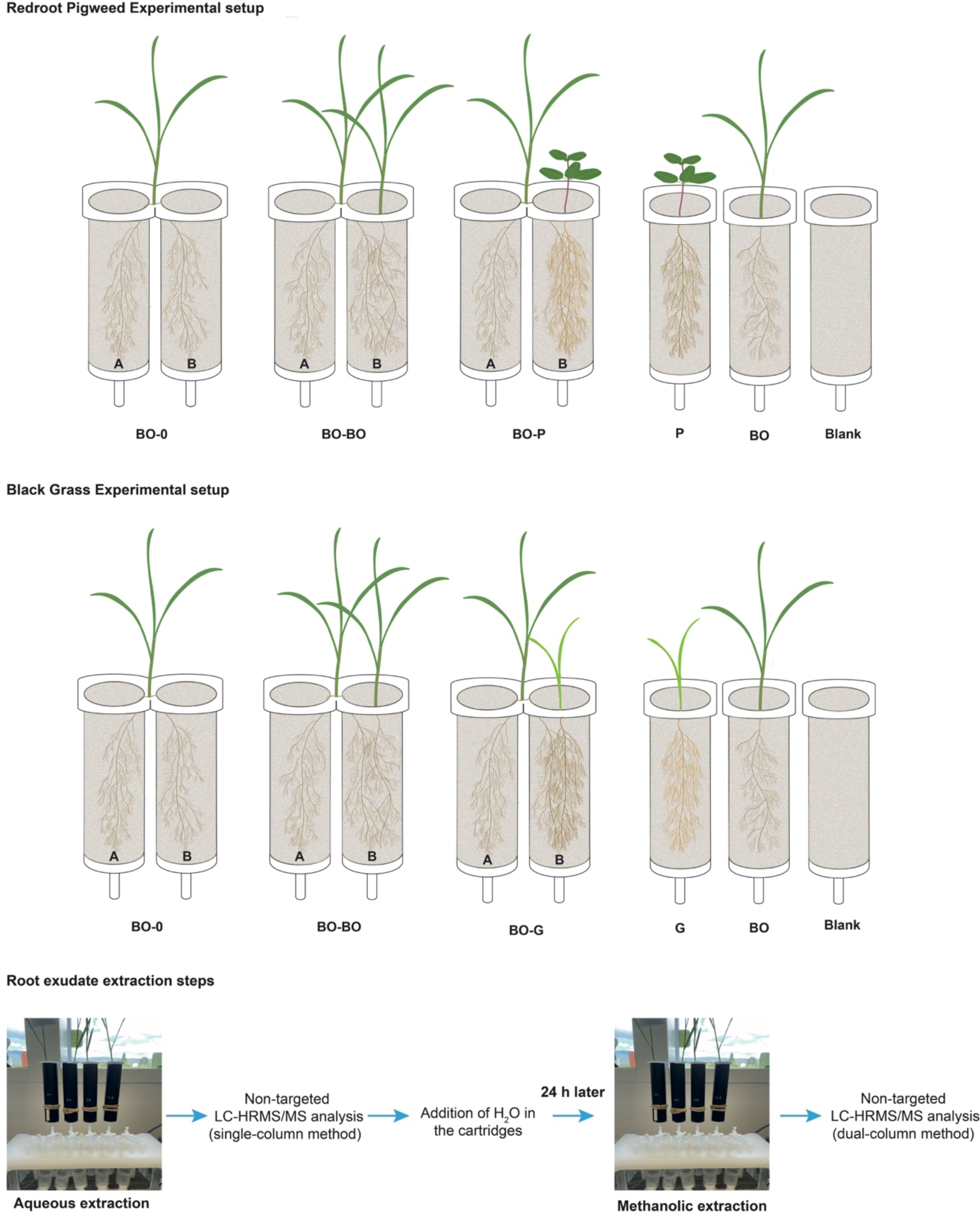
Experimental Setup of Black Oat (BO) Split-Root Systems. The experimental setup comprises various growth conditions. Redroot Pigweed Experimental setup: BO-0: Black oat split-root without any neighbour. Half of the root system is in the first compartment (**BO-0/A**), and the other half is in the second compartment (**BO-0/B**). BO-BO: Black oat split-root with a black oat neighbour. Half of the root system is in the first compartment without root contact with the black oat neighbour (**BO-BO/A**), while the other half is in the second compartment in direct root contact with the black oat neighbour (**BO-BO/B**). BO-P: Black oat split-root with an interspecific redroot pigweed (P) neighbour. Half of the root system is in the first compartment without root contact with the redroot pigweed neighbour (**BO-P/A**), while the other half in the second compartment is in contact with the redroot pigweed neighbour (**BO-P/B**). Blackgrass experimental includes aforementioned BO and BO-BO conditions and also BO-G: Black oat split-root with an interspecific blackgrass (G) neighbour in which half of the root system is in the first compartment without root contact to the blackgrass neighbour (**BO-G/A**), while the other half in the second compartment is in contact with the blackgrass neighbour (**BO-G/B**). BO, and P and G: Non-split root plants in single cartridges: BO: black oat, P: redroot pigweed G: blackgrass and Blank: cartridge filled with glass beads and no plants. Root exudates were initially vacuumed off from each cartridge using nano pure water. After adding water and allowing a 24-hour incubation, the root exudates were extracted again using a methanolic extraction solution.

### 2.3. Manifold setup and root exudate extraction

Root exudates were collected from all cartridges of split and non-split-root plants grown in SPE tubes using an SPE vacuum manifold (Macherey-Nagel) connected to a vacuum pump (V-300, Buchi) controlled by an interface (Buchi I-300 Pro). Conical centrifuge tubes, 50 ml (Corning) were positioned on a rack under the stainless-steel needles of the manifold. The interface was set to 780 mbar, maintaining the glass chamber pressure at 5 mmHg. Each cartridge was placed on the stopcock valves. The valves were opened, and 30 mL of nano pure H_2_O was added to each cartridge over 30 seconds simultaneously. The valves were left open for an additional 30 seconds to vacuum off the root exudates, totalling 1 minute of vacuum per sample. The contents of the centrifuge tubes constituted the aqueous extract. Root exudates from split root cartridges were extracted using two adjacent valves simultaneously by adding the extraction solution to both cartridges. Following the aqueous extraction 15 ml of nano pure water was added to each cartridge and plants were placed back in the Phytotron.

After 24 hours, a second extraction was performed by adding 30 mL of extraction solution consisting of 95% (w/v) methanol (Merck Uvasol), 4.95% (w/v) nano pure water, and 0.05% (w/v) formic acid (VWR, HiPerSolv Chromanorm for LC-MS) with internal standard 3,5-di-tert-butyl-4-hydroxybenzoic acid (Sigma-Aldrich) (final concentration of 0.5 µmol L^-1^). This solution was applied over the span of 30 seconds and vacuumed off using the same manifold system mentioned above for a total of one minute.

Aliquots of 15 ml from each sample obtained through first (water) and second (methanol) extractions were transferred to Pyrex test tubes (SciLabware) and evaporated using a sample vacuum concentrator (Genevac EZ-2 Plus) set at 35 °C. The samples were not dried to completion. For the first hour, the Genevac was set to “HPLC mode.” For the remaining time, until samples were ∼200 µL in volume, the Genevac was set to “aqueous mode.”

### 2.4. Root sample preparation, digitization, and analysis

Following the root exudate extraction, root systems of each plant growing in split-root and non-split-root conditions were digitalised by scanning them using WinRHIZO^TM^ Basic 2021 software (Regent Instruments Inc.) and its root positioning and scanning accessories.

The plants were gently removed from the cartridges, making sure that each root system stayed intact. The roots were washed to remove any glass beads stuck on their surface. Each root sample was then submerged in Reagent’s tray filled with water. The tray was positioned on the scanner glass, and the roots were carefully spread out meticulously to prevent them from crossing on top of each other, then they were scanned and analysed. The root analysis performed using WinRHIZO^TM^ provided root traits such as total root length (mm), root surface area (mm^2^), average total root diameter (mm), root volume (mm^3^), and number of root tips and branching points. Dry root and aboveground weight were also recorded after drying the tissues in a drying oven at 50°C for 48 hours and weighing them using an analytical scale.

### 2.5. Black oat root exudate treatment on redroot pigweed and blackgrass

The root exudates used for treatment were extracted using the same method as explained in the root exudate extraction section. However, for the second extraction performed after 24 hours, nano pure water was used instead of the extraction solution. The second extracts were used to prevent varying Hoagland’s salt concentrations across different experimental conditions. This was verified by checking the pH and electrical conductivity of each sample. For each experimental condition, 10 samples were pooled together to obtain a homogeneous mixture for each treatment. Ten redroot pigweed seeds and five blackgrass seeds were sown in separate single SPE cartridges filled with glass beads. Three millilitres of half-strength Hoagland’s solution were added to each tube daily for the first five days. Both redroot pigweed and blackgrass seedlings were thinned down to three plants on the fifth day. Over the next ten days, the redroot pigweed seedlings were given 3 ml of BO-0, BO-P/A, and BO-P/B, and blackgrass seedlings were given BO-0, BO-G/A, and BO-G/B root exudates, and control plants received 3 ml of nano pure water every day. All the plants also received 1 ml of half-strength Hoagland’s solution every other day to ensure they had enough nutrients (n=10). At the end of the treatment, root morphology analysis was performed using WinRHIZO^TM^ on redroot pigweed and blackgrass seedlings as described in the previous section.

Statistical significance (p < 0.05) of root traits were assessed by conducting either a student’s t-test or one-way ANOVA. As a post hoc test, Dunnett’s multiple comparisons were performed by setting BO-0 as the control and comparing each mean against it. Statistical analysis was performed using GraphPad Prism (version 9.2.0 for MacOS, GraphPad Software).

### 2.6. LCMS sample preparation

Pre-concentrated sample extracts (∼200 µL) were transferred to 1.5 mL HPLC amber glass vials (Altmann Analytik) and reconstituted to 10% methanol and 0.1% formic acid and to be a 30-fold concentration of the original volume (500 µL total).

At initial extraction, particulate was not an issue as the glass beads acted as a filter. However, aqueous extract samples formed particulates during pre-concentration due to the high concentration of salts. These samples were ultracentrifuged (Sorvall Discovery M150 SE) to remove particulate for 15 minutes at 125000g and 4 °C. Precipitate and particulate were not an issue with second MeOH extracts, and they were not ultracentrifuged.

Quality control (QC) samples were comprised of a pooled sample from each experiment set (redroot pigweed and blackgrass) or extraction type (first aqueous and second methanolic) for a total of 4 different QC samples and data sets (redroot pigweed aqueous, redroot pigweed methanolic, blackgrass aqueous, and blackgrass methanolic). Samples were separated into two aliquots for negative and positive ionization modes and stored at −80 °C until analysis.

### 2.7. Mock root exudate mixture

A mock root exudate mixture was made up of 58 chemical standards at 10 µmol/L concentration (see Table S1 for details). These were compounds known from the literature to be exuded by the roots of agricultural plants. This mixture was used to assess the repeatability of the signal intensity and retention times of the instrument on different days of LCMS analysis. Additionally, they were used to create an in-house database for high-level identification of these 58 compounds within the samples.

### 2.8. LC-HRMS/MS data acquisition

#### 2.8.1. Setup

The LC-HRMS/MS system was controlled by Agilent MassHunter acquisition software (version 10.1). A cooled autosampler unit, two binary pumps, and a temperature-controlled column compartment (1290 Infinity II) brought the sample to the primary sprayer of a 6560-ion mobility QTOFMS with a Dual AJS ESI interface. A nano pump for reference mass solution (1260 Infinity) led to the secondary sprayer. All equipment was from Agilent Technologies.

Samples were thawed, vortexed, and stored at 4°C in the autosampler until injection. For all methods, the injection volume was 5 μL, the flow rate was 350 μL min^−1^, and the column temperature was 50 °C. All samples were randomized for injection order. A QC sample was run every 14-16 injections.

A single-column method was implemented for the first aqueous extract samples. The stationary phase was comprised of a Discovery HS F5 (150 x 2.1 mm, 3 μm particle size, Sigma-Aldrich) pentafluorophenyl (PFP) column paired with a Discovery HS F5 Supelguard Cartridge. Mobile phase A was ultrapure H_2_O with 0.1% FA; Mobile phase B was MeOH with 0.1% FA. Gradient details can be seen in Table S2A, and the total analysis time was 15.5 minutes per sample.

For the second methanolic extract of 24 regeneration samples, a dual-column method was utilised. Both the PFP column and Hypercarb porous graphitic carbon (PGC) column (2.1 × 150 mm, 5 μm particle size, Thermo Scientific) equipped with a Hypercarb PGC guard column were connected to a valved within the column compartment which controlled the flow from the two separate binary pump systems. For pump 1, mobile phases A and B were the same as previously mentioned. For pump 2, mobile phase A was again ultrapure H_2_O with 0.1% FA, but mobile phase B was ACN with 0.1% FA. This method had two discrete stages: the retention stage (both columns in tandem) and the elution stage (each column with its pump and gradient) (Bennett et al., 2024). This dual column method allows for a more diverse set of compounds to be analysed when compared to a single column method, but it is not viable for the first aqueous extract due to the PGC column not being robust with matrix variation. Gradient details can be seen in Table S2B&C and the analysis was 21 minutes per sample.

While there were two LC methods, the MS method was the same for both. Online mass calibration was performed using reference mass solution (HP-0921, Purine, TFANH4; m/z=121.0509 and 922.0098 (+); m/z= 119.0363 and 966.0007 (−)). Data was acquired using data-dependent acquisition (DDA) with negative and positive ionization modes performed as separate runs. The scan range was 50 to 1700 m/z. A detailed HR-MS/MS acquisition method report can be seen in Table S2D.

### 2.9. LC-HRMS/MS data analysis

#### 2.9.1. Data pre-processing

Data was recalibrated and centroided with Agilent reprocessing software. The first step of pre-processing was performed in MS DIAL (version 4.9.221218). A void volume cutoff was applied (1.6 minutes for the single-column method and 2.0 minutes for the dual). All isotopologues but not their adducts were aggregated together. Using an extract of a compartment with glass beads and no plant as a blank, a 20% blank filter was applied. Data was normalized to the internal standard (for the second extract only) and pooled QC samples using LOWESS regression (for all extracts). Gap filling was utilised to input baseline values for samples containing no detectible amount of an aligned compounds to avoid issues with missing or zero values during statistical analysis. All MS DIAL pre-processing traits are in Table S3. Further, pre-processing was performed in R. Adducts were aggregated with their protonated and deprotonated features to generate a total compound signal (TCS), an S/N cutoff of 10 was applied, and samples were normalized biologically to root weight.

#### 2.9.2. Statistical analysis

Statistical analysis was performed in R on TCS values. Outliers were defined by the 1.5x interquartile range rule, calculated, and removed. Among-group variation was assessed through F-testing and/or Bartlett’s test. Normality was assessed through Shapiro-Wilk’s test. Significant differences between experimental conditions among the two redroot pigweed or blackgrass experimental setups (the A compartment three different split root conditions – BO-0/A, BO-BO/A, and BO-P/A or BO-G/A – and two different single cartridge conditions – BO and P or G) were determined using Welch’s t-test. A false discovery rate (FDR) multiple-test correction was performed for the t-tests. Fold change was calculated pairwise between each experimental condition. Data was centred and auto scaled, and then partial least square discriminate analysis (PLS-DA) was performed on the different experiment sets separately. Data had to be acquired and thus analysed between the first and second extractions of the P and G experimental sets separately due to the necessity of a QC sample to normalize the data sets and limitations of the analytical equipment. For this reason, all analysis and statistics based upon relative quantification comparison refer to these 4 different data sets (firs extraction and second extraction of the P and G experimental sets).

#### 2.9.3. Identification

Identification was performed from levels 1 to 4 according to Schymanski et al., (2014) For level 1 identification, an in-house database was generated using MS FINDER from the 58 mock root exudate standards and this database was applied to the sample sets within MS DIAL. Level 2 identification was divided into 2a (spectral database match) and 2b (in silico database match). Level 2a was assessed using all available databases within GNPS (Global Natural Products Social Molecular Networking) and MS FINDER. If multiple results were returned, the results with the best score were favoured. If each database returned a result for level 2a, MS FINDER results were favoured. Since each database frequently used different synonyms for the same compound, it was not viable to assess if the databases returned the same or different results for each compound. Level 2b was determined by SIRIUS and MS FINDER. For level 2b, a structure score cutoff was applied (>6 for MS FINDER and > 0.6 for SIRIUS) and the structure with the highest score of all formulas was selected. Again, MS FINDER was the favoured database if both returned a result for a single compound. The CANOPUS feature of SIRIUS used fragmentation pattern fingerprints to determine compound classes (level 3) of each metabolite.

For determining superclass, a class threshold of >0.80 and a superclass threshold or >0.95 was applied and then the compound class with the highest score, irrespective of molecular formula associated, was assigned. For subclass, a threshold of >0.90 was used. Molecular formulas (level 4) were generated using SIRIUS and MS FINDER utilizing parent m/z values and adduct/ionization form information.

Additionally, since identification is based upon databases and algorithms trained on well know metabolites, they tend to favour primary metabolites and well studies specialized metabolites. For this reason, known black oat specialized metabolites which were unlikely to be represented within databases such as avenaol (Kim et al., 2014), scopoletin (Fay & Duke, 1977), and scopolin were manually searched for using known fragments from the literature (377.160 ➔ 97, 235, and/or 263 in positive ionization mode for avenaol; 191.035 ➔ 104.03, 148.02, and/or 176.01 in negative ionization and 193.050 ➔ 122.04, 133.03, and/or 178.03 in positive ionization for scopoletin; 353.088 ➔ 176.01 and/or 191.04 in negative ionization and 355.102 ➔ 133.03, 178.026, and/or 193.05 in positive ionization for scopolin) as they were not present and/or matched to within any available database.

#### 2.9.4. Prioritization & visualization

Metabolites which had a significant p-value (<0.05) after FDR correction and had a log_2_ fold change >0.6 or <-0.6 were prioritized. P-values and fold change results were graphed against each other in volcano plots for visualization of specific metabolites of interest when comparing two different experimental groups. PC1 and PC2 of the PCA and PLS-DA were graphed against each other to visualize differences between three or more sample groups. All samples from the second extracts were utilized to generate four feature-based molecular networks through GNPS (positive and negative ionization and P experimental set and G experimental set). This assessed the relationship of each compound to one another through comparison of fragmentation patterns. Related compounds were given a cosign score with a threshold of 0.6 to quantify how similar they were. All compounds were plotted in relation to one another. As node proximity to one another is not a perfect visualization of compound relatedness with this method, the line thickness of the edge between nodes was based on the cosine relationship score. Identification information was integrated; nodes were coloured by compound class (identification level 3). Identification level 1 and 2a compounds were annotated by name. Confidence level 1 and 2a labels for identified compounds were placed above the nodes they represent. Prioritization information was integrated by highlighting compounds of interest in bold black outlines around the representative node. Compounds upregulated by redroot pigweed in BO-P/A when compared to BO-0/A were noted by triangular nodes, and those upregulated by blackgrass in BO-G/A when compared to BO-0/A were noted by square nodes.

Compounds which were prioritized were given extra attention beyond the automated identification and annotation. The different outputs from each of the processes outlined in the identity confirmation section were assessed to see if they agreed with one another, the fragmentation patterns, and experimental data present. While the identification thresholds of the different software were not ignored, if multiple results passed a threshold, a lower ranked annotation was preferred over higher rank annotation if compounds with which it was connected to within the network corroborated with the lower prioritization result.

## 3. Results

### 3.1. Effect of Black Oat on Neighbouring Redroot Pigweed and Blackgrass

Growing weeds with split BO roots and having direct root-root contact led to significantly lower root length, specific root length, root volume, number of root tips, root surface area, and aboveground dry weight in neighbouring redroot pigweed plants (NP) compared to redroot pigweed growing in a single cartridge without a BO neighbour (P) (Figure 2a, Figure2b, Figure2d, Figure 2e, Figure 2f), while average root diameter and root dry weight showed no significant difference (Figure2c) and aboveground weight was significantly higher in NP (Figure 2g).

**Figure 2:**
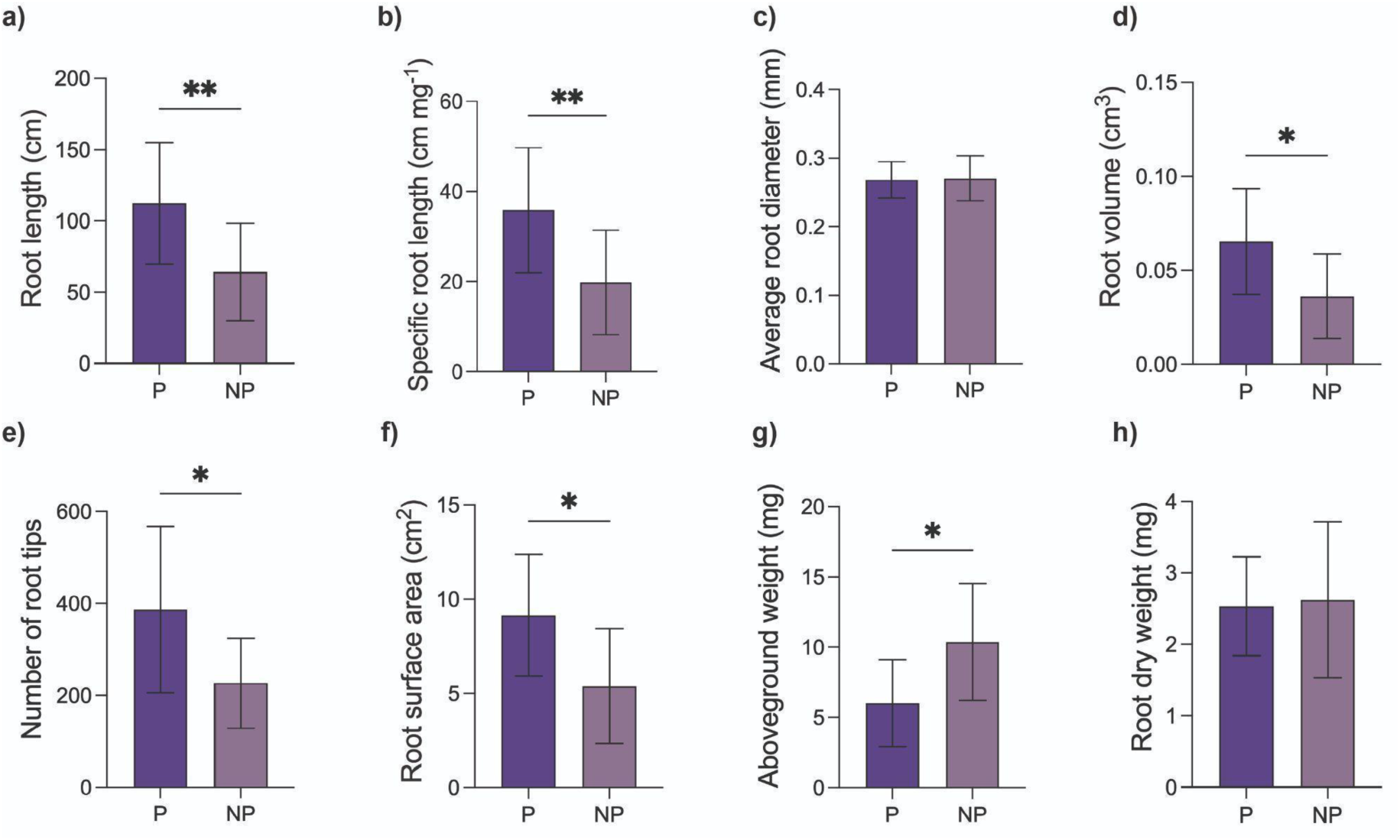
The impact of a black oat neighbour on redroot pigweed root traits. The comparison between redroot pigweed plants growing alone **(P)** and those growing with a split-root black oat **(NP)**. Root Length **(a)**, Specific root length **(b)**, Average Root Diameter **(c)**, Root Volume **(d)**, Number of Root Tips **(e)**, Root Surface Area **(f)**, Aboveground Dry Weight **(g)**, Root Dry Weight **(h)**. The bars represent the mean (n = 15), with error bars indicating the standard deviation. One-way ANOVA with Dunnett’s multiple comparison post hoc test was conducted to determine significant differences in root traits (n = 5, 95% confidence level, p > 0.05 *p ≤ 0.05, **p ≤ 0.01, ***p ≤ 0.001.

The black oats growing alone in a single cartridge were compared with neighbour black oats (NBO) growing with split root BO in a shared compartment (BO-BO/B). While root length, root volume, number of root tips, root surface area, aboveground dry weight, and root dry weight were significantly lower in NBO than BO (Figure 3a, Figure 3d, Figure 3e, Figure 3f, Figure 3g, Figure 3h) specific root length and average root diameter traits were not significantly different among BO and NBO (Figure 3b and Figure 3c).

**Figure 3:**
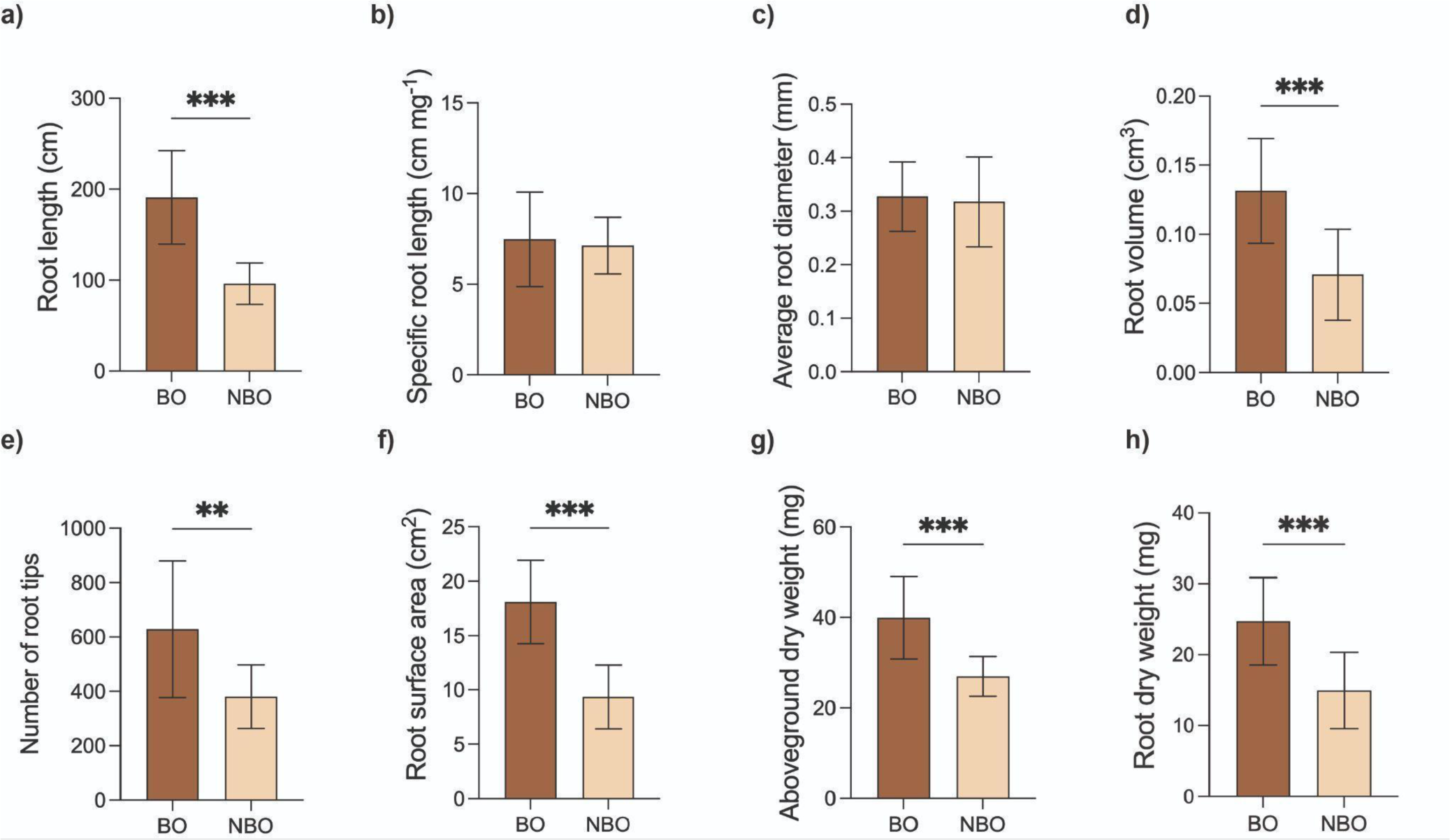
The influence of having an interspecific neighbour black oat on black oat root traits. Black oat Root Length **(a)**, Specific root length **(b)**, Average Root Diameter **(c)**, Root Volume **(d)**, Number of Root Tips **(e)**, Root Surface Area **(f)**, Aboveground Dry Weight **(g)**, Root dry weight **(h)**. Root traits of black oat growing alone (BO) and black oat growing with a black split root (NBO). The bars indicate the mean (n = 15), and the error bars indicate the standard deviation. One-way ANOVA with Dunnett’s multiple comparison post hoc test was performed to determine significant differences in root traits. n = 15, 95% confidence level, p > 0.05 *p ≤ 0.05 **p ≤ 0.01 ***.

Root traits of blackgrass plants growing in a single cartridge (G) were also compared with neighbouring blackgrass (NG) from BO split root systems where they were sharing the same cartridge (BO-G/B) (Figure 4). Root length, specific root length, root volume, number of root tips, root surface area, and root dry weight of NG were significantly increased (Figure 4a, Figure 4b, Figure 4d, Figure 4e, Figure 4f, Figure 4h), while there was no significant change in average root diameter and aboveground dry weight (Figure 4c and Figure 4g).

**Figure 4:**
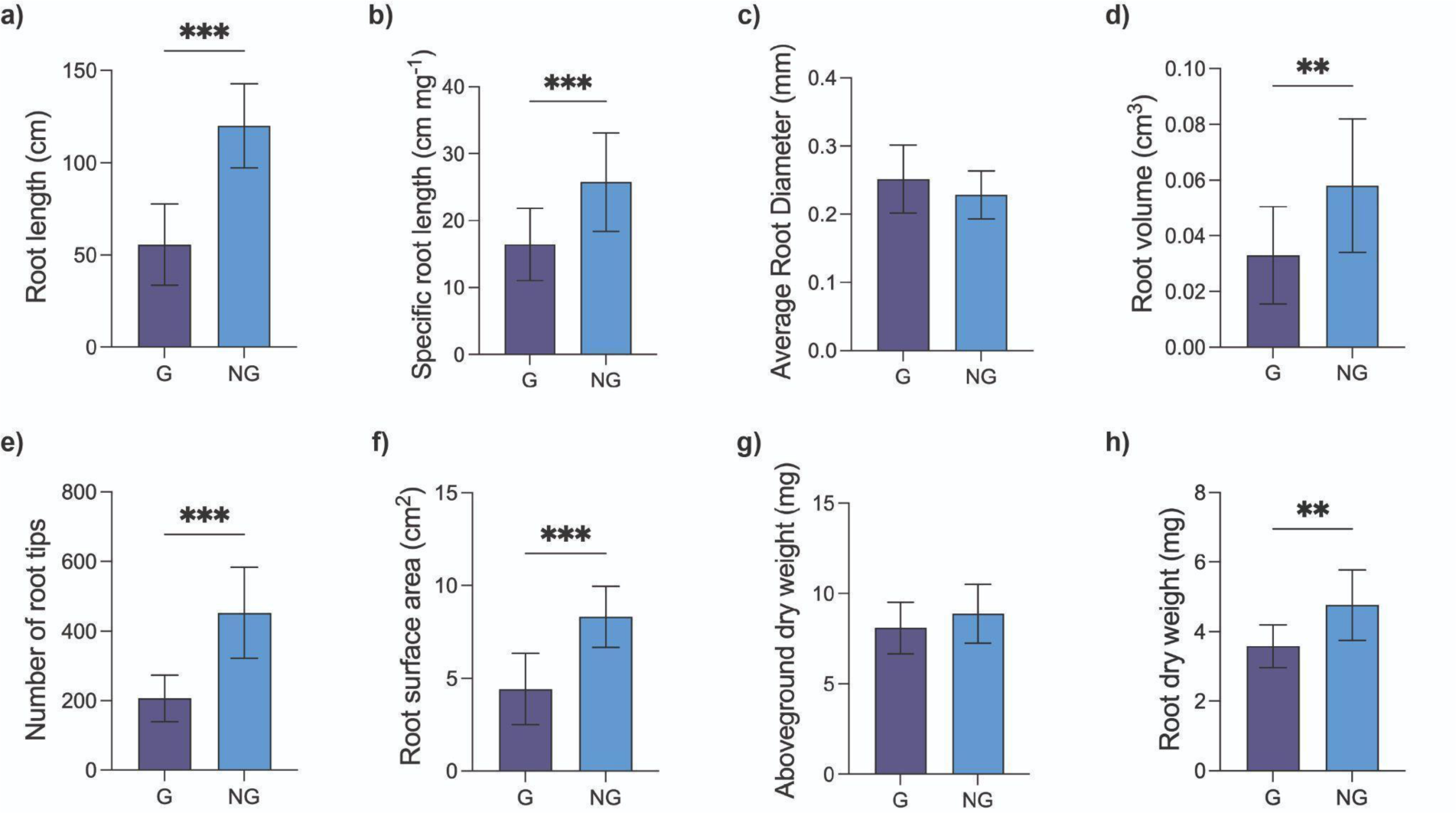
The impact of black oat neighbour on blackgrass root traits. The blackgrass plants growing alone **(G)** and those growing with a split root black oat **(NG)** were compared. Blackgrass Root Length **(a)**, Specific root length **(b)**, Average Root Diameter **(c)**, Root Volume **(d)**, Number of Root Tips **(e)**, Root Surface Area **(f)**, Aboveground Dry Weight **(g)**, Root Dry Weight **(h)**. The bars indicate the mean (n = 15), and the error bars indicate the standard deviation. One-way ANOVA with Dunnett’s multiple comparison post hoc test was performed to determine significant differences in root traits. n = 5, 95% confidence level, p > 0.05 *p ≤ 0.05 **p ≤ 0.01 ***.

### 3.2. Root Exudate Treatment on Redroot Pigweed and Blackgrass

Application of root exudates from black oats grown in split-root systems without a neighbour (BO-0) significantly increased the average root diameter, root volume, root surface area, and root dry weight in redroot pigweed (P) seedlings (Figure 5b, Figure 5c, Figure 5e and Figure 5g). In addition, the number of root tips was significantly reduced in P under the same treatment (Figure 5d). When P seedlings were treated with root exudates from BO growing with neighbouring redroot pigweed, either in direct root contact (BO-P/B) or without root contact (BO-P/A), a reduction in the number of root tips was observed, with significant changes only in the BO-P/B treatment (Figure 5d). Furthermore, root length, root volume and root dry weight were significantly lower in P seedlings treated with BO-P/B compared to plants treated with BO-0 (Figure 5a, Figure 5c, and Figure 5g). No significant changes in aboveground dry weight were observed in P across any treatment (Figure 5f).

**Figure 5:**
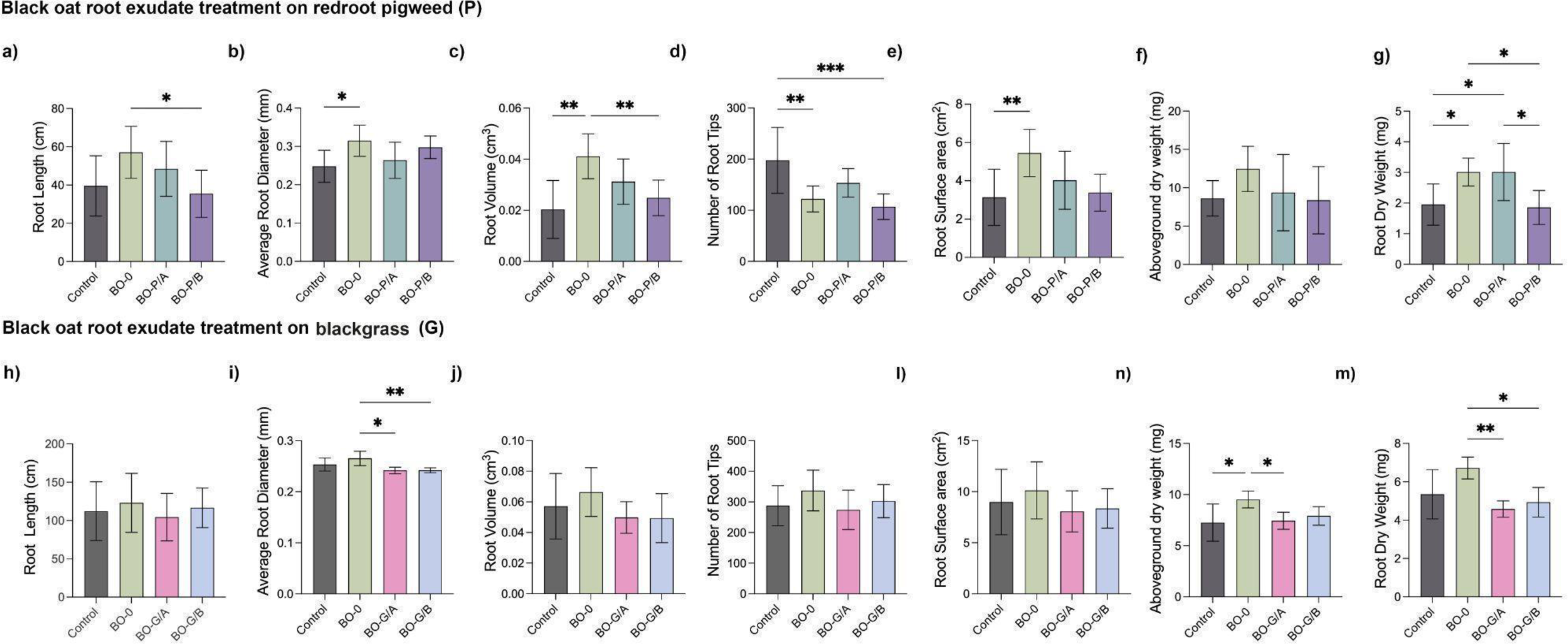
Effect of root exudates collected from black oat split-root systems on redroot pigweed and blackgrass root traits. The first line of graphs shows redroot pigweed traits, and the second line shows blackgrass traits: Root length **(a, h)**, Average Root Diameter **(b, i)**, Root Volume **(c, j)**, Number of Root Tips **(d, k)**, Root Surface Area **(e, l)**, Above ground weight **(f, n)**, Root dry weight **(g, m)**. Redroot pigweed and blackgrass seedlings received root exudates obtained from the A compartments of black oat split-root systems without neighbours **(BO-0/A)**. P seedlings received root exudates obtained from black oat split-root systems with P neighbours: having no direct root contact **(BO-P/A)**, and with direct root contact **(BO-P/B)**, while G seedlings received root exudates obtained from split-root systems with G neighbours: having no direct root contact **(BO-G/A)**, and direct root contact **(BO-G/B)**, and all controls received nano pure water. The bars indicate the mean (n = 5-8, each data point is an average of 3 P and G seedlings). Error bars indicate the standard deviation. One-way ANOVA with Dunnett’s multiple comparison post hoc test was performed to determine significant differences in root traits, at a 95% confidence level, p > 0.05 *p ≤ 0.05 **p ≤ 0.01 ***.

For blackgrass (G) seedlings, treatment with root exudates from BO split-root systems resulted in significant changes only in average root diameter, aboveground dry weight and root dry weight (Figure 5i, Figure 5n, and Figure 5m). Application of root exudates from BO growing with G neighbours (BO-G/A and BO-G/B) resulted in significantly lower average root diameter in G compared to treatment with BO-0 exudates. However, no significant differences were observed when compared to the controls. Additionally, no differences in root length, root volume, number of root tips and root surface area (Figure 5h, Figure 5j, Figure 5k, 10l) were observed across any treatment in G seedlings.

### 3.3. Statistical analysis of LC-HRMS/MS root exudate data

Within all the PCAs (Figure S1), the 11 QC samples analysed for each experimental setup (redroot pigweed aqueous, redroot pigweed methanolic, blackgrass aqueous, and blackgrass methanolic) clustered together. Root exudate composition of the different plant species was analysed in this experiment and both P and G root exudate samples separated from BO root exudate samples. In addition, there was a high degree of overlap for all the samples extracted from BO plants, regardless of the interaction / tested condition.

Similarly, the PLS-DA (Figure 6) also showed a high degree of overlap for the first aqueous extraction for both the P and G experimental sets when comparing BO-0/A, BO-BO/A, BO-P/A in P experimental sets and BO-0/A, BO-BO/A, BO-G/A in G experimental set. For the second methanolic extraction, there was a better separation in the PLS-DA, but there was still some overlap between groups within the centre of the PLS-DA for both P and G experimental sets.

**Figure 6:**
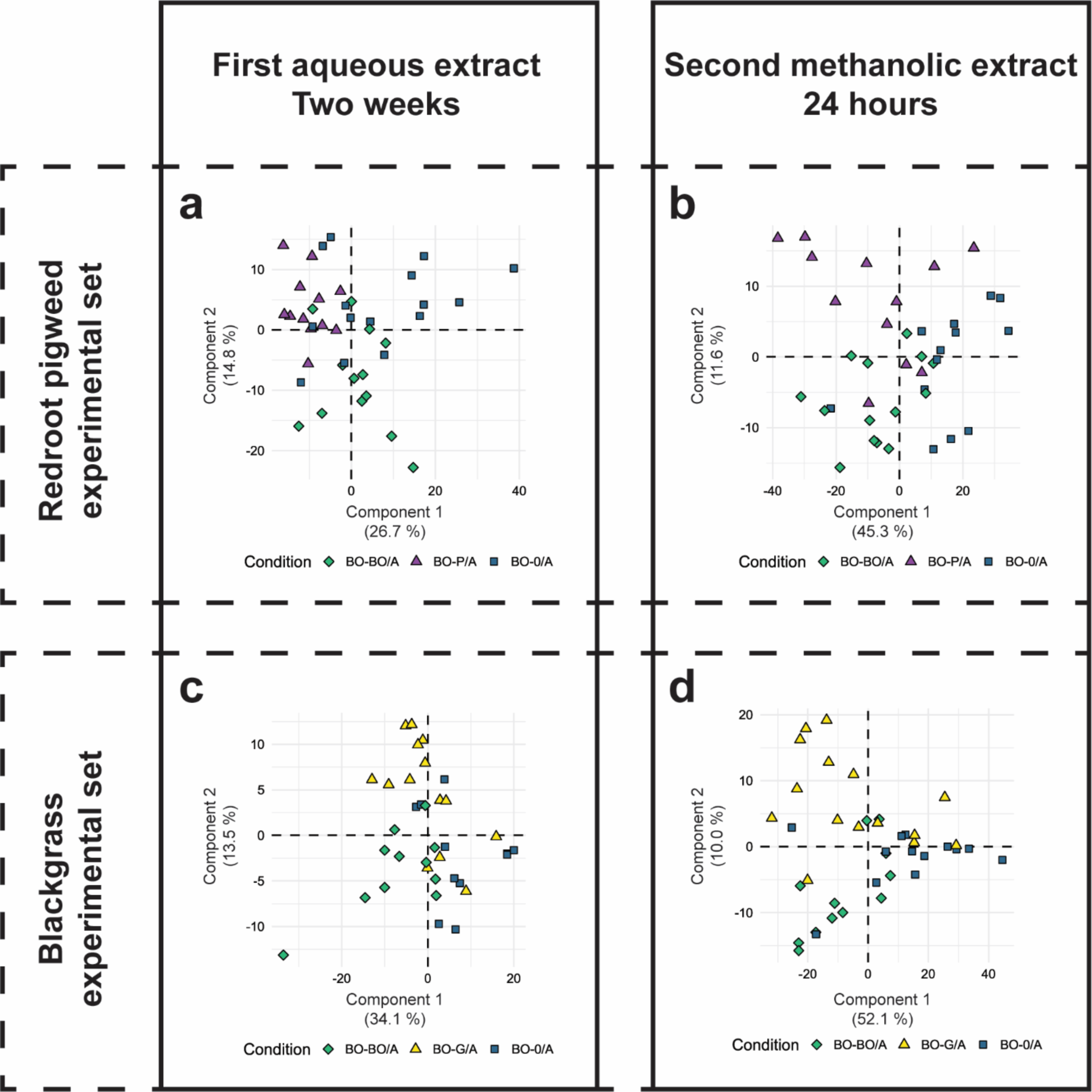
The PLS-DA of P experimental set samples highlight differences between compartment A with no neighbour (BO-0/A), with a same species neighbour (BO-BO/A), and with a P neighbour (BO-P/A) groups for the aqueous extraction of exudates with water after two weeks **(a)** and the second extraction after a 24-hour re-exudation period with methanol **(b)**. A PLS-DA of G experimental set samples (BO-0/A, BO-BO/A, and BO-G/A) makes the same comparisons for the first **(c)** and second **(d)** extracts.

To assess which metabolites were specifically different between groups, univariate statistics were performed. After data preprocessing, there were 705 compounds from the alignment file (containing compounds from all samples and conditions) compared in negative ionization mode for aqueous extract of the P experimental set. However, not all samples contained all 705 of these aligned compounds as gap filling used baseline nonzero values for samples which lacked one or more of the compounds in the alignment file. There were 613 compounds compared in positive ionization mode. For the aqueous extract of the G experimental sets, 317 and 321 compounds were aligned and compared in negative and positive ionization mode, respectively. When grown alone, 337 compounds were considered more expressed in P than BO in the root exudates (Figure S2) and 145 more in G than BO root exudates (Figure S3). With BO-0/A used as a control group, no metabolites were observed to be upregulated in BO-P and only 11 were observed in the G experimental set in the aqueous extraction (Figure 7a and Figure 7c).

**Figure 7:**
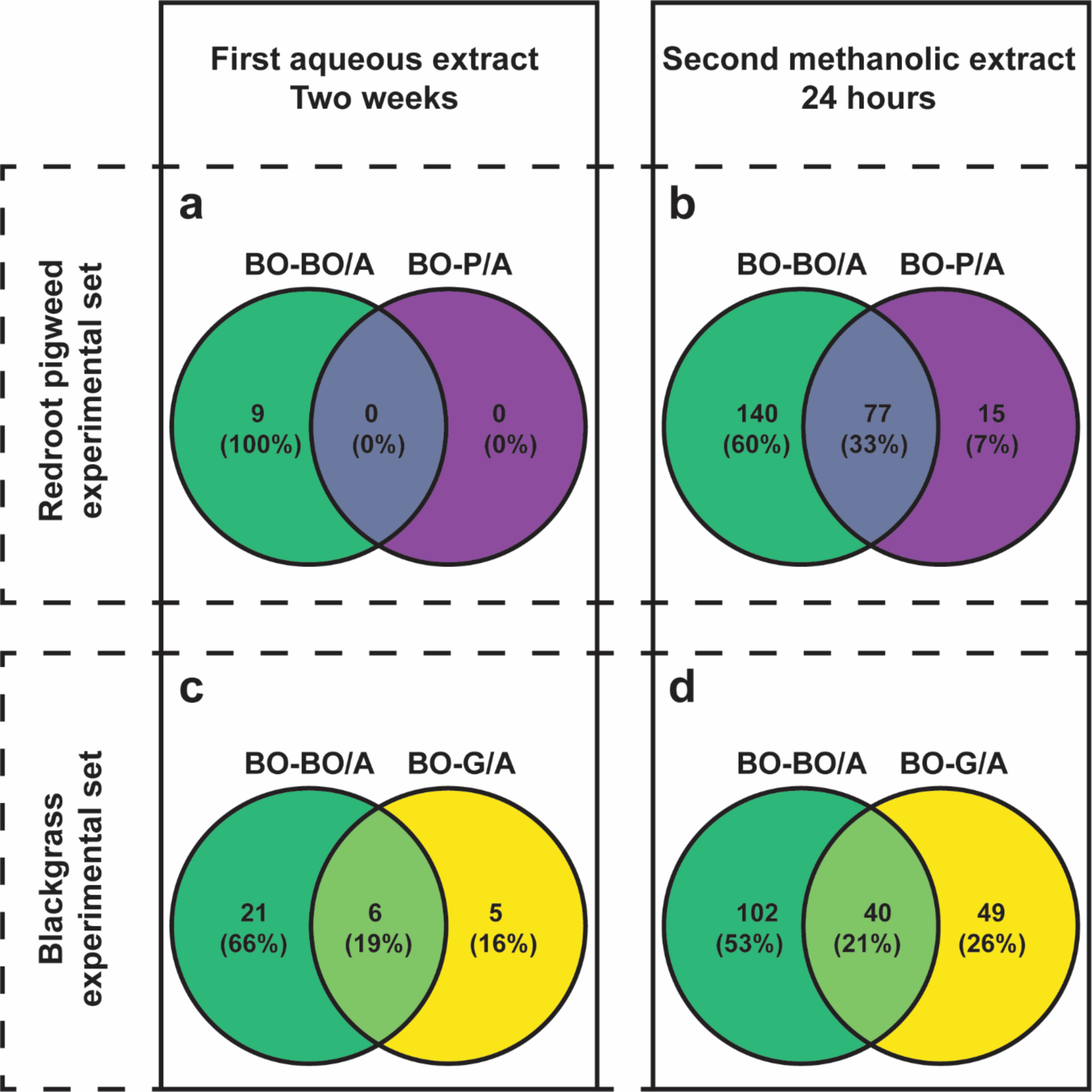
The Venn diagrams from the P experimental set highlight the number of compounds upregulated by the presence of a redroot pigweed (BO-P/A) and/or black oat (BO-BO/A) neighbour when each is compared to no neighbour (BO-0/A) and the number of unique or shared upregulated compounds between the two conditions for the aqueous extraction of exudates with water after two weeks **(a)** and the second extraction after a 24-hour re-exudation period with methanol **(b)**. The same Venn diagrams were generated for G experimental set, but highlighted compounds upregulated by the presence of blackgrass (BO-G/A) comparisons for the first **(c)** and second **(d)** extracts. Upregulation was defined as any compound significantly higher in one group when compared to another as determined by a Welch’s t-test (p<0.05) and a log_2_ fold change greater than 0.6.

For the second extracts, there were 952 and 597 compounds analysed in negative and positive ionization mode for the P experimental set and 674 and 620 compounds in the G experimental set. When grown alone, 150 compounds were considered more expressed in P than BO and 283 more in G than BO. Compared to BO-0/A, 92 compounds (Table S6 and Figure S6) were upregulated in BO/P-A and 89 (Table S7 and Figure S7) were upregulated in BO-G/A (Figure 7b and Figure 7d). These results align with the PLS-DA results indicating larger degree of observable differences within the second extract.

### 3.4. Metabolite identification

Depending on the data set, ∼1-2% of compounds were annotated to confidence level 1 and ∼2-4% and 4-7% were identified to confidence levels 2a and 2b, respectively (Table S8 & S9). 45-65% were annotated to confidence level 3 (Figure S8 & S9). Superclass annotation of different experimental groups from the second extract show that the upregulated compounds (i.e., upregulated by the presence of any neighbour in comparison BO-0/A) are comprised of more organic oxygen compounds (Figure 8). Subclass annotation from CANOPUS shows that amino acids and carbohydrates are present in both the first and second extracts (Figure S10).

**Figure 8:**
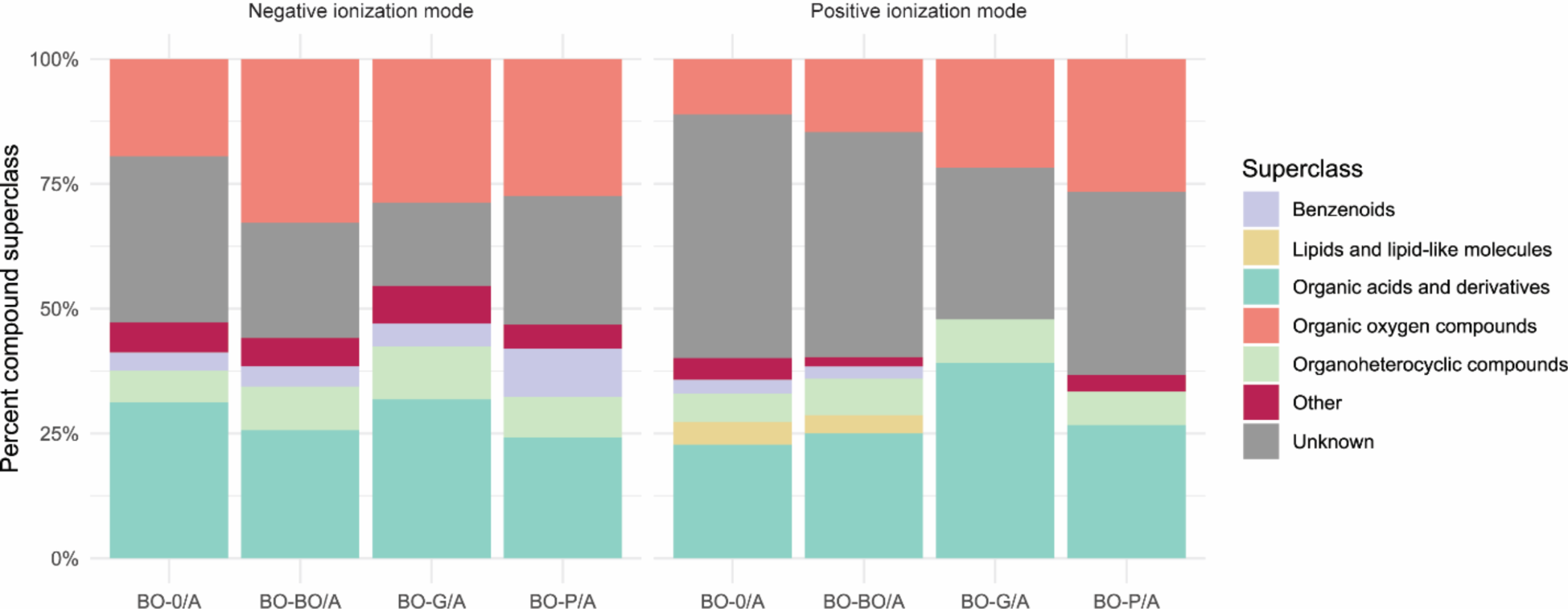
Compounds classes of exudates were annotated using the CANOPUS application of SIRIUS. Since all samples had no missing values and a nonzero value for every compound after alignment, split root black oat with no neighbour (BO-0/A) compounds were considered those which were not expressed significantly higher in redroot pigweed (P) or blackgrass (G) when compared to BO-0/A as denoted by a Welch’s t=test (p<0.05) and a log_2_ fold change greater than 0.6. Compounds that were upregulated by the presence of an intraspecific neighbour (BO-BO/A), an interspecific redroot pigweed neighbour (BO-P/A), and an interspecific blackgrass neighbour (BO-G/A) were those which were significantly higher than BO-0/A using these same thresholds. Compound classes which comprised less than 5% of the total number of compounds for that ionization mode were binned together into the “other” category.

Avenaol was not found within the sample extracts, either due to poor compatibility with the analytical method or due to it not being present within the sample extracts. Scopoletin and scopolin was found within the black oat root exudate extracts, but neither was differentially expressed when comparing any of the A compartments (BO-0/A, BO-BO/A, and BO-G/A).

### 3.5. Molecular network

Molecular networking was used to enhance the characterization of compounds present in second extracts obtained from both P and G experimental sets, providing insight into their structural relationships and the chemical complexity of BO root exudate composition (Figure 9). This process was first validated by confirming that the compounds grouped together according to their respective superclass, given that compounds from the same class share structural similarities and have similar fragmentation patterns. Further validation was achieved by ensuring that the confidence level 1 and 2 compounds which were annotated to be of a similar structure were positioned adjacent to each other in the molecular network. For example, a cluster of organic oxygen compounds in the negative ionization mode from P experimental set highlights that three sugars strongly connected. Further clusters show a series of phophatidylethanolamines and phosphatidylglycerol annotated as part of the lipid and lipid-like superclass and clustering together with other unknown organic acids and derivatives. Annotated amino acids also cluster amongst one another under the organic acids and derivatives superclass. Additionally, the networks highlight that compounds identified to confidence level 1 and 2a match the predicted compound superclass (e.g., specific amino acids determined to be organic acids and derivatives and specific sugars shown to be organic oxygen compounds).

**Figure 9:**
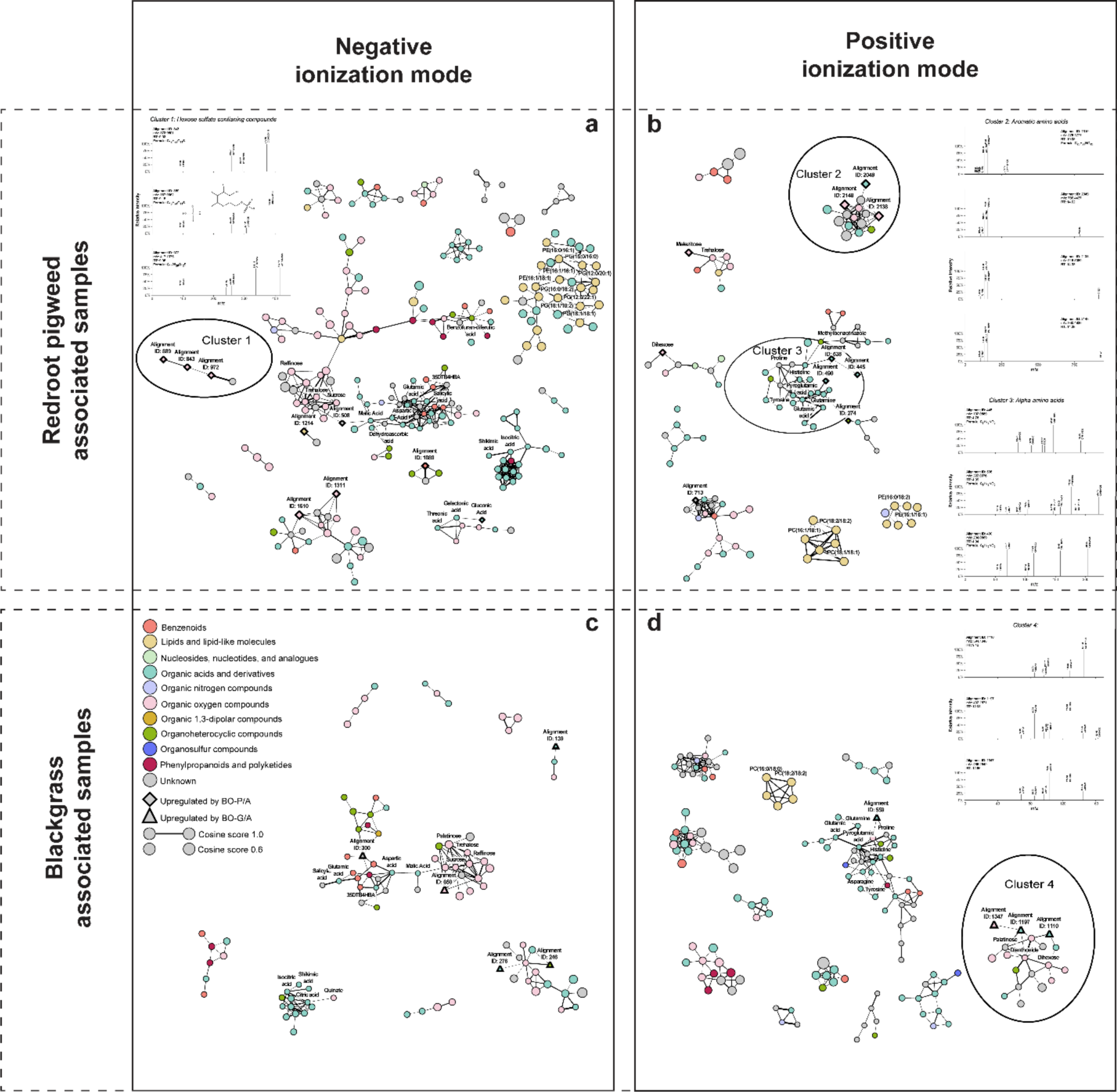
A molecular network generated using GNPS shows how compounds are related based upon their fragmentation patterns for the P experimental set in negative ionization mode (a) and positive ionization mode (b) and the G experimental set in negative ionization mode (c) and positive ionization mode (d). Node colour represents compound class as determined by the CANOPUS application of the SIRIUS open-source software. Node size represents the m/z of the analyte with larger nodes representing a larger m/z. Edge width directly relates to the cosine relationship score indicating the strength of the fragmentation similarity with thicker edges indicating a higher score. Diamond shaped nodes represent compounds that are upregulated by the presence of a redroot pigweed neighbour (BO-P/A) when compared to no neighbour (BO-0/A) as determined by a Welch’s t=test (p<0.05) and a log2 fold change greater than 0.6. Triangle nodes denote compounds upregulated by the presence of a blackgrass (BO-G/A) neighbour. Three clusters containing upregulated compounds are highlighted. Due to a lack of space, phophatidylethanolamines (PE) and phosphatidylglycerol (PG) abbreviations were used within the network.

Four upregulated clusters, three including compounds that were upregulated in BO in the presence of P neighbours, were highlighted. Fragmentation patterns from cluster 1 shows the upregulated compounds share a hexose sulphate functional group. This is supported not only by the presence of the sulphate and sulphate + hexose fragments but also by the SO_3_ neutral loss in all fragmentation patterns. Within cluster 2, there is a compound (Alignment ID 1310) that, though not one of the upregulated compounds, is comprised of a dihexose, a carboxyl group, and an amino group attached to a benzene ring. The benzene, amino, and carboxyl indicating fragments are shared among all upregulated compounds within the cluster indicating that these are all aromatic amino acids, though it is unclear if these compounds also contain a dihexose. Upregulated compounds in cluster 3 have less definitively shared fragments, but they all are within the organic acids and derivatives superclass, cluster near amino acids, share a carboxylic acid functional group loss with the fragmentation pattern, and were annotated as amino acids by CANOPUS. Cluster 4, from compounds upregulated in BO by the presence of G, contains 3 upregulated compounds, but not much structural information can be determined as there are few shared fragments which do not concretely give structural information and there is little other annotation from SIRIUS.

## 4. Discussion

Root exudates are key players in plant signalling and mediate interactions between plants and neighbouring organisms such as microbiota in the rhizosphere, herbivores, and other plants sharing the same environment (Mondal et al., 2023; N. Wang et al., 2021). Therefore, in this study, we aimed to determine the effects of plant neighbours on root exudate composition and root traits.

### 4.1. Root trait alterations in response to the presence of black oat differ among neighbouring black oat, redroot pigweed and blackgrass

Roots demonstrate plasticity, the ability of the root system to adapt under changing soil conditions to reduce the impact of stress, in the presence of neighbouring plants (H. Wang et al., 2006; Yamauchi et al., 1994) in the presence of neighbouring plants. Root morphology and physiology are altered in response to growing in proximity to other plants (Jacob et al., 2017). In our study, we observed a common pattern of significant decreases in several root traits (root length, volume, surface area, and number of root tips) in both NP and NBO plants when interacting with split-root BO plants, as well as differing responses in dry weights (Figure 2 and Figure 3). These changes could be explained by several mechanisms or processes, such as; (i) competition for resources (e.g., light, nutrients, and water) and space (Craine & Dybzinski, 2013), (ii) chemical signalling occurs via root exudates and their possible alterations by the microbial community (Wu et al., 2023) induced by the presence of neighbours and neighbour root exudates, (iii) stress responses such as phytohormonal changes (Lyu & Smith, 2022). Decrease in root length in the presence of neighbours could be associated with the optimisation of resource allocation to other parts of the plant. On the other hand, it has been suggested that root growth can directly respond to the presence of neighbours, independent of resource availability as well, through root exudation or other alterations in the physicochemical conditions (McNickle, 2020). In previous studies, the presence of neighbours significantly decreased the length of first-order lateral roots of wheat plants in the presence of black grass neighbours under high nutrient conditions (Finch et al., 2017). In our experimental setup, plants were sufficiently provided with nutrients to minimize competition for resources. Another study also reported that the root allocation patterns of pea plants were altered by the presence of both pea and oat neighbours. In our previous study, we also observed a similar suppression pattern in P interacting with buckwheat plants (Eroğlu et al., 2024).

The changes in root dry weight and aboveground weight in the presence of BO split-root neighbours varied among species. Both root and aboveground dry weight decreased in NBO interacting with BO split-root plants. On the contrary, the presence of BO neighbours led to increased aboveground weight in NP. Presence of BO also decreased root volume, root surface area, and the number of root tips in both NP and NBO, while these root traits increased in NG. NBO and NP roots might have avoided the roots of split root BO to reduce direct competition leading to lower root proliferation hence decreased root volume, surface area and number of tips in compartment B where they share the same space. Therefore, the increase in NP aboveground dry weight might be explained by a shift in resource allocation towards aboveground growth to absorb more light and have a competitive advantage. (Figure2g). However, as G did not respond in the same way as P and BO, other factors could be at play as well. The root exudate composition of plants might be species-specific, and dependent on the genetic relatedness of plants of interest (Bais 2006, Shemchenko 2014). Studies conducted on various species have shown differences in root exudate composition, including model plants (McLaughlin et al., 2023), five grass and five forb species (Dietz et al., 2020) and 65 plant species from 25 different plant families (Rathore et al., 2023). Rathore et al. (2023) demonstrated that phylogeny was found to be important in explaining differences in root traits. They also showed that the composition of root exudates could be partially predicted by root traits such as root length, root dry weight, and root diameter, although most of the variation is too complex to be explained solely by root traits. These variations impact not only plant-plant interactions but also plant-microbiome interactions. The alterations in microbial communities including the changes in their activity, abundance, and composition have a significant impact on plant growth and root exudate composition (Pétriacq et al., 2017; Sasse et al., 2018). The responses to BO root exudates trigger in neighbouring plants lead to different alterations in the root system architecture of various species, including the weeds P, G, and BO itself. P is a dicotyledon summer weed thriving in high temperatures, tolerating drought, and needing high nitrogen, while G is a monocotyledon autumn weed favouring cooler, moist conditions, and they germinate in different seasons. Even though BO and G are both in the Poaceae family, while P belongs to the distant family Amaranthaceae, the differences in their responses cannot be simply explained by genetic relatedness as the root trait changes induced by BO on its neighbours of the same species were similar to those observed in P rather than G. Presence of BO and P neighbours might have perceived as same type of stress and triggered similar response in BO, while G caused a different type of competitive challenge. Split root BO and neighbour BO are intraspecific neighbours which are likely to have similar resource requirements and root exudates that would trigger specific responses. BO is more likely to grow in the same ecological niche as other BO plants and they would expect to compete stronger with each other than interspecific neighbours and needing stronger adaptive responses. On the other hand, even though BO and G are both from Poaceae family, they don’t share a close evolutionary history as they are from different plant families and genera. Genus Avena is closely related to crops such as wheat, barley, and rye but still differs from those genetically and chemically (Kellogg, 1998; Menon et al., 2016; Valentine et al., 2011). Also, as P thrives in a wide range of habitats (Hamidzadeh Moghadam et al., 2021) it is more likely to share a habitat with BO than G. Previous studies demonstrated that plant roots responded neighbours in a widely varied manner by showing no response to segregating or over-proliferating (Belter & Cahill, 2015). Therefore, aligning with previous findings, we also did not observe a unified BO response to different species of neighbours. Although the responses to P align with our previous study on buckwheat, where buckwheat presence induced similar changes in P root traits as BO did, this was not the case with G. This suggests that there might be species-specific strategies. However, it is important to note that this is only a limited comparison of three different species which are P, G and BO interacting with BO plants. To make a definitive conclusion larger scale studies are required.

### 4.2. Different effects of root exudates collected from split root BO interacting with P and G on root traits of P and G

When applied to P, root exudates collected from BO split root plants with no neighbours (BO-0) lead to a significant increase in average root diameter, root volume, root surface area and root dry weight of P in comparison to controls. As root exudates provide carbon and nitrogen inputs, they can be utilized by both the microbiome and neighbouring plants in the environment (Hu et al., 2018). BO root exudates might have supplied growth-promoting sugars and amino acids to plants and microorganisms, potentially leading to increased root traits. Interestingly, while BO-0 exudates seemed to promote overall root growth, they had an inhibiting effect on root tip formation.

The root traits of P plants treated with exudates from BO directly interacting with P (BO-P/B) and those indirectly interacting (BO-P/A) exhibited some differences. It is highly likely that the local responses generated by BO is stronger than systemic responses in the second compartment where there are no competing roots. While the significant decrease observed in number of root tips following the root exudate application reflects the significant decrease observed in number of root tips P in split root systems when grown with a BO neighbour, parameters such as root length and volume were only significantly decreased when compared to BO-0/A. In addition to direct competition that takes place in the split-root systems, the concentration of root exudate metabolites plays a significant role in neighbouring plants and the microbial community present. Higher concentrations of root exudate metabolites can have a more pronounced inhibitory or stimulatory effect on the growth of neighbouring plants and microorganisms, whereas lower concentrations might result in weaker /negligible impacts. This dose-dependent effect of root exudates on target plant and microbial species has been observed in previous studies (Dayan et al., 2009; Gniazdowska & Bogatek, 2005; Macías et al., 2020; Upadhyay et al., 2022). The effect of BO root exudate treatments on G root traits were different from P. This aligns with the difference we observed in split root systems. G does not respond neither to the presence of BO nor the presence of BO root exudates same way as P. Overall, P shows more substantial changes in root traits under when treated with root exudates obtained from directly interacting plants (BO-P/B) whereas, G is less affected by BO-G/B treatment. Suggesting that P may be more sensitive to the root exudates obtained from interacting plants and possibly the allelochemicals present than G.

### 4.3. Neighbour presence leads to shifts in black oat root exudate metabolome

Root exudate analysis showed that the presence of P and G neighbours shifted the metabolome profile of split-root BO, which had no direct root-root contact with the weed neighbours (BO-P/A and BO-G/A) compared to BO without a neighbour (BO-0/A). Previous studies suggest that plants can modify their root exudates in response to environmental factors, including neighbouring plants. Root exudates trigger different responses in plants depending on the neighbour species/ identity and generate different local and systemic responses (Semchenko et al., 2014). In our previous study, we showed that the composition of buckwheat root exudates differed depending on whether buckwheat or redroot pigweed was its neighbour (Eroğlu et al., 2024).

In this study, we expanded from one extract to two. When comparing differences in metabolite profile, each extraction provided different benefits. The aqueous extract, closer to natural conditions, was of interest for its possibility to observe the undisturbed differential accumulation of exudates. Therefore, reflecting the natural composition across two weeks without creating stress on the plant that an organic solvent may generate. Alternatively, the second extract was important as Hoagland’s solution was mostly eliminated after being washed off by the aqueous extraction, allowing for minimised matrix variation and ion suppression from nutrient salts (King et al., 2000) which poses a potential problem for the analytical method. This is specifically why it would not be beneficial to mix the two extractions to reduce the number of analytical runs. Second, the methanol allows for better extraction of phenolics (Alara et al., 2021) and lipid-like molecules from the exudate solution. While there is a risk of the methanol lysing the cell membranes of the root, this risk is relatively low as the methanol is only applied to the roots for a minute (Pétriacq et al., 2017) and is diluted by the aqueous solution in the cartridges. Last, this extraction allows for a 24-hour circadian snapshot of root exudate release.

While the differences between experimental conditions (compartment A of the split root experiments) in the aqueous extracts were not as pronounced and there were still overlaps between samples within the PCA (Figure 6a, 6c, 7a, 7c), methanolic extraction highlighted these differences slightly better (Figure 6b, 6d, 7b, 7d) as more of the exudate variation among samples could be explained by the different experimental groups for the 24-hour extraction when compared to the two-week extraction. This difference between extracts might be due to the aforementioned presence of the Hoagland’s salt mixture in the aqueous extracts which can cause ion suppression, especially for small polar metabolites eluting within the same range as the salts, resulting in sensitivity reduction. Moreover, it can lead to excessive salt adduct formation which makes identification challenging (Furey et al., 2013; Leitner et al., 2007).

Another reason for the lack of perceived differences in the aqueous extraction could be because the aqueous extraction contains root exudates produced over a prolonged period of two weeks, while the second extraction contains those produced only within 24 hours. Root exudation is a dynamic process that changes over time. Previous studies show the root exudates released within 24 hours after transplanting also have more variation than those collected after two weeks of growing with a neighbour, indicating that root exudate compounds reach saturation over time (McLaughlin et al., 2023). Hence leading to more consistent exudate profiles across samples as BO plants achieve homeostasis in their root exudation patterns. This could result in exudate saturation in the rhizosphere and thus less variation in exudate composition after two weeks. The removal of the exudates during the aqueous extraction could potentially disrupt this homeostasis, allowing for the observation in differences in exudation profiles between experimental groups/neighbours in the second extraction after 24 hours.

Both extracts were utilized to investigate compounds upregulated in response to the presence of weeds, P and G, due to the aforementioned individual information each extraction could provide. In comparison to the aqueous extraction, the methanolic extraction revealed a substantial increase in compounds when BO interacted with a neighbour (Figure 7b and Figure 7d). In both experimental sets, the presence of an intraspecific BO neighbour led to the upregulation of a higher number of compounds in comparison to the metabolites upregulated by the presence of P and G. Interestingly, the same pattern was observed in buckwheat when grown with an intraspecific neighbour or redroot pigweed. There are two possible reasons for this repeated trend; 1) the plants have the same exact resource demands and thus are in direct and heavy competition and these exudates are part of a negative response (Ehlers et al., 2016) or 2) plants of the same species are responding positively to one another. While the logic of the first postulation is there, it is not directly obvious why they would have a positive interaction. However, the literature shows that plants of the same species do interact and respond positively to one another within the rhizosphere (Adler et al., 2018).

In response to the presence of any weedy neighbours, organic oxygen compounds were consistently shown to be upregulated (Figure 8). Organic oxygen compounds are a vast group of compounds which include alcohols, aldehydes, ketones, ether, carboxylic acids and peroxides (Schmiermund, 2023). More interestingly, organic oxygen compounds are also comprised of well-known components of root exudates such as sugars. Additionally, some specific amino acids (e.g., glutamate, aspartate, and serine) and organic acids are classified at both organic oxygen species and as organic acids and derivatives. These compounds would be preferentially annotated as organic acids and derivatives by our workflow (i.e., they would not cause the observed increase in oxygen compounds). Within the network, many of the organic oxygen compounds were further annotated through level 1 and 2 identifications as carbohydrates, indicating an increase in exudation of sugars from BO when grown with any neighbour.

### 4.4. Molecular networking highlights increased presence of sugars and glycosides within black oat exudate solution in response to redroot pigweed neighbours

The network in Figure 9 highlights several interesting findings for the redroot pigweed experimental setup. First, there is an upregulation of sugars (Melezitose and an unknown dihexose) and of some unidentifiable amino acids in cluster 3 when black oat is grown with a redroot pigweed neighbour. Plants might release certain compounds to change the microbial community for their benefit. They could shape microbial communities to recruit beneficial microbes, (Kong & Liu, 2022) and this could be induced by the presence of neighbouring plants (Zhou et al., 2023). The sugars and/or amino acids detected in the root exudates might be part of this response. Root exudates with high sugar content may attract beneficial microbes which in turn could help improve plants in the acquisition of nutrients and the suppression of pathogens. The presence of P neighbours might have triggered BO to release root exudates with higher sugar content in order to recruit beneficial microbes to alleviate the competition stress (Bais et al., 2006; Ulbrich et al., 2022). Also, sugars found in the root exudate solution (Berlemont & Martiny, 2016) could potentially be breakdown products of glycosides, as microbes in the rhizosphere produce glycosidases that break down complex carbohydrates into simpler sugars (Ali et al., 2019).

Second, within cluster 1 there are three compounds which are sulphate containing compounds with a glycoside. There is one well known group of compounds which contains both a sulphate and glycosylation: glucosinolates produced most by the order Brassicales. The compounds upregulated within cluster 1 differ from glucosinolates as they lack a sulphur bridge and nitrogen, but both contain a characteristic sugar and sulphate. Glucosinolates contribute to plant defence against pests and disease and are even used as a material for bio fumigation of soils to control weeds as the breakdown isothiocyanates are enzymatically toxic (Craine & Dybzinski, 2013; Hanschen et al., 2015). While these plants are not related and thus it is highly unlikely that the pathways producing these compounds are related, there is compelling evidence that sugar sulphates may be protective compounds that both lineages have developed independently.

Last, cluster 2 contains three aromatic amino acids. While this seems unrelated to the sugar and sugar sulphates, they are linked within the network to an aromatic amino acid with a glycoside. This glycoside is not observable within the upregulated aromatic amino acids within the cluster. However, glycosides are often only observed through neutral losses within mass spec fragmentation patterns in positive ionization mode (Qu et al., 2004), as is with the case for alignment ID 1310 which is the related sugar containing aromatic amino acid. Since the parent ions are in such low abundance in the MS2 spectrum, it is not surprising that the fragmentation pattern lacks an observable neutral loss for a sugar if it contains one. One supporting piece of evidence that these compounds may contain sugars is the annotation from SIRIUS CANOPUS which labels these compounds both as peptides (Level 5 classification) and gives a possible alternative annotation of glycosylamine for Alignment ID 2049 and 2138.

Current research is conflicting if plants can actively uptake glycosylamines or not with certain studies showing active uptake (Roberts et al., 1971; Camacho-Pereira et al., 2009) and others arguing that this only occurs at extremely high concentrations (P. Roberts & D.L. Jones 2012). However, several studies in higher plants have shown that glycosylamines are potent inhibitors of respiration, hexokinase activity, and interfere with the oxidative pentose phosphate pathway, an important source of biosynthetic precursors and reducing power in plants (Garlick et al., 2002; Hofmann & Roitsch, 2000; P. Roberts & Jones, 2012; Sanz & Ullrich, 1989). As shown here, glycosylamines are also highly toxic to maize roots even at low concentrations (R. M. Roberts et al., 1971).

### 4.5. Concluding statements

The objective of our study was to investigate how the presence of interspecific weed neighbours, redroot pigweed and blackgrass, and the intraspecific neighbour black oat impact root exudate composition and root traits in split-root conditions and upon exposure to root exudate treatments. It is more challenging to identify subtle changes in the exudates of a single plant species in the presence of different neighbours than to detect differences between the root exudate compositions of plants from different species, as they have distinct root metabolomes, as we observed with black oat, redroot pigweed, and blackgrass. The compounds upregulated in the presence of neighbours mostly belonged to organic oxygen superclass, with amino acids and carbohydrates identified. The intraspecific neighbour led to the upregulation of more compounds than the weed species, which might be due to triggering more intense competitive responses and/or inducing the production of allelochemicals that facilitate more effective competition. Black oat plants, having larger root systems than the weeds, might be involved in more direct resource competition compared to weeds with smaller root systems. We observed alterations in root exudate composition in the presence of redroot pigweed, with the upregulation of sugar sulphates which might be protective compounds that help plant cope with stress as well as amino acids and a general increase in root exudation. Root morphology and physiology varied depending on the species interacting with black oat; while root exudates obtained from black oat interacting with redroot pigweed had suppressive effects on root traits, those obtained from black oat interacting with blackgrass had the opposite impact.

The compounds produced in response to interactions with redroot pigweed tentatively characterized, could be utilized to improve the effectiveness cover crop use in weed management. Future studies should focus on different plant species to draw conclusions about general differences between intraspecific and interspecific interactions. Moreover, the underlying molecular mechanisms leading to the production of these compounds, which might be potential allelochemicals, as well as the role of microbial communities and their effects on plant-plant interactions, need to be explored. Further studies should also test more complex matrices, such as using soil or conducting field trials, to test applicability and scalability.

## Acknowledgements

We would like to thank EQ-BOKU VIBT GmbH and the BOKU Core Facility Mass Spectrometry.

## Author Contributions

All authors have read and approved this manuscript. This work has not been accepted or published elsewhere. CE and AB conducted the lab experiments, analyses, data collection and analysis, and prepared the figures and wrote the manuscript.

