## Supplemental material for "Characterization of root exudates of black oat in the presence of interspecific weed species neighbours and intraspecific neighbours, and their effects on root traits"

**Supplementary**

**Table S1:** A series of compounds across five major classes (small organic acids, amino acids, sugars, flavonoids, and fatty acids) known to exudate from the roots of mostly agricultural plants were selected for assessing the chromatographic and data processing workflows. Each compound's name, molecular formula, exact mass in Da, and chemical ontology/class is listed. The species each compound is known to be exudated from, the source of each standard, and the purity of each standard is given. One compound, 3,5-Di-tert-butyl-4-hydroxybenzoic acid, is a synthetic and non-naturally occurring compound used as in internal standard.

| Compound name | Molecular formula | Exact mass (Da) | Ontology | Species | Source of chemical standard |
|---|---|---|---|---|---|
| **Urea** | CH4N2O | 60.0324 | Organic molecular entity | *Arabidopsis thaliana*[1], *Lactuca sativa* L.[2] | Sigma-Aldrich |
| **Glycine** | C2H5NO2 | 75.0320 | Amino acid | *Arabidopsis* thaliana[1], Triticum *aestivum* L.[3], *Brachypodium distachyon*[4] | Sigma-Aldrich |
| **Butyric acid** | C4H8O2 | 88.0524 | Organic acid | *Triticum aestivum* L.[3], | Sigma-Aldrich |
| **β-Alanine** | C3H7NO2 | 89.0477 | Amino acid | *Arabidopsis thaliana*[1], *Triticum aestivum* L.[3], *Brachypodium distachyon*[4], *Zea mays*[5] | Fluka |
| **Oxalic acid** | C2H2O4 | 89.9953 | Organic acid | *Triticum aestivum* L.[3] | Merck Millipore |
| **Valeric acid** | C5H10O2 | 102.0681 | Organic acid | *Triticum aestivum* L.[3] | Fluka |
| **Y -Amino butyric acid, GABA** | C4H9NO2 | 103.0633 | Amino acid | *Arabidopsis thaliana*[1], *Triticum aestivum* L.[3], *Brachypodium distachyon*[4], *Zea mays*[5] | Sigma-Aldrich |
| **Serine** | C3H7NO3 | 105.0426 | Amino acid | *Arabidopsis* thaliana[1], Triticum *aestivum* L.[3], *Brachypodium distachyon*[4] | Sigma-Aldrich |

| | | | | | |
|---|---|---|---|---|---|
| **Uracil** | C4H4N2O2 | 112.0273 | Nucleic Acids | *Arabidopsis thaliana*[1] | Sigma-Aldrich |
| **Proline** | C5H9NO2 | 115.0633 | Amino acid | *Triticum aestivum* L.[3] | Sigma-Aldrich |
| **Fumaric acid** | C4H4O4 | 116.0110 | Organic acid | *Arabidopsis* thaliana[1], Triticum *aestivum* L.[3] | Fluka |
| **Valine** | C5H11NO2 | 117.0790 | Amino acid | *Arabidopsis thaliana*[1], *Triticum aestivum* L.[3], *Brachypodium distachyon*[4] | Merck Millipore |
| **Succinic acid** | C4H6O4 | 118.0266 | Organic acid | *Arabidopsis* thaliana[1], Triticum *aestivum* L.[3] | Fluka |
| **Threonine** | C4H9NO3 | 119.0582 | Amino acid | *Arabidopsis thaliana*[1], *Triticum aestivum* L.[3], *Brachypodium distachyon*[4], *Zea mays*[5] | Sigma-Aldrich |
| **Benzoic acid** | C7H6O2 | 122.0368 | Organic acid, Phenol | *Arabidopsis thaliana*[1], *Hordeum vulgare*[6] | Fluka |
| **Erythritol** | C4H10O4 | 122.0579 | Sugar alcohol | *Arabidopsis thaliana*[1], *Zea mays*[5] | Sigma-Aldrich |
| **Pyroglutamic acid** | C5H7NO3 | 129.0426 | Amino acid | *Arabidopsis thaliana*[1], *Sorghum bicolor* L. Moench[7] | Sigma-Aldrich |
| **Isoleucine** | C6H13NO2 | 131.0946 | Amino acid | *Arabidopsis thaliana*[1], *Triticum aestivum* L.[3], *Brachypodium distachyon*[4], *Zea mays*[5] | Sigma-Aldrich |
| **Leucine** | C6H13NO2 | 131.0946 | Amino acid | *Triticum aestivum* L.[3], *Brachypodium distachyon*[4], *Zea mays*[5] | Sigma-Aldrich |
| **Asparagine** | C4H8N2O3 | 132.0535 | Amino acid | *Arabidopsis thaliana*[1], | Sigma-Aldrich |

| | | | | | |
|---|---|---|---|---|---|
| | | | | *Triticum aestivum* L.[3],<br>*Brachypodium distachyon*[4],<br>*Zea mays*[5] | |
| **Aspartic acid** | C4H7NO4 | 133.0375 | Amino acid | *Triticum aestivum* L.[3],<br>*Brachypodium distachyon*[4] | Merck Millipore |
| **Malic acid** | C4H6O5 | 134.0215 | Organic acid | *Triticum aestivum* L.[3] | Merck Millipore |
| **Threonic acid** | C4H8O5 | 136.0372 | Organic acid | *Arabidopsis thaliana*[1],<br>*Sorghum bicolor* L. Moench[7] | Sigma-Aldrich |
| **Salicylic acid** | C7H6O3 | 138.0317 | Organic acid, Phenol | *Medicago sativa*[8],<br>*Triticum aestivum, Avena fatua,* many others[9] | Fluka |
| **Glutamine** | C5H10N2O3 | 146.0691 | Amino acid | *Triticum aestivum* L.[3],<br>*Brachypodium distachyon*[4],<br>*Zea mays*[5] | Sigma-Aldrich |
| **Lysine** | C6H14N2O2 | 146.1055 | Amino acid | *Triticum aestivum* L.[3],<br>*Brachypodium distachyon*[4] | Sigma-Aldrich |
| **Glutamic acid** | C5H9NO4 | 147.0532 | Amino acid | *Triticum aestivum* L.[3],<br>*Brachypodium distachyon*[4],<br>*Zea mays*[5] | Sigma-Aldrich |
| **Methionine** | C5H11NO2S | 149.0510 | Amino acid | *Triticum aestivum* L.[3],<br>*Brachypodium distachyon*[4] | Sigma-Aldrich |
| **Ribose** | C5H10O5 | 150.0528 | Sugar | *Arabidopsis* thaliana[1],<br>Triticum *aestivum* L.[3] | Sigma-Aldrich |
| **Dopamine** | C8H11NO2 | 153.0790 | Phenol | *Brachypodium distachyon*[10] | Sigma-Aldrich |
| **Histidine** | C6H9N3O2 | 155.0695 | Amino acid | *Brachypodium distachyon*[4],<br>*Oryza sativa*[11] | Fluka |

| | | | | | |
|---|---|---|---|---|---|
| **Courmaric acid** | C9H8O3 | 164.0473 | Organic acid, Phenol | *Medicago sativa*[8], *Ageratum conyzoides*[12], *Eucalyptus spp.*[13], *Triticum vulgare*[14], *Avena fatua*[15] | Fluka |
| **Phenylalanine** | C9H11NO2 | 165.0790 | Amino acid, Phenol | *Triticum aestivum* L.[3], *Brachypodium distachyon*[4] | Sigma-Aldrich |
| **Vanilic acid** | C8H8O4 | 168.0423 | Organic acid, Phenol | *Hordeum* vulgare[6], *Medicago sativa*[8], *Eucalyptus spp.*[13], *Triticum vulgare*[14], *Avena fatua*[15], *Fagopyrum esculentum*[16] | Alfa Aesar |
| **Shikimic acid** | C7H10O5 | 174.0528 | Organic acid | *Arabidopsis thaliana*[1], *Oryza sativa*[11] | Sigma-Aldrich |
| **Arginine** | C6H14N4O2 | 174.1117 | Amino acid | *Brachypodium distachyon*[4], *Oryza sativa*[11] | SAFC Pharma |
| **Aldohexose (glucose)** | C6H12O6 | 180.0634 | Sugar | *Arabidopsis thaliana*[1], *Triticum aestivum* L.[3], *Zea mays*[5] | Sigma-Aldrich |
| **Tyrosine** | C9H11NO3 | 181.0739 | Amino acid, Phenol | *Triticum aestivum* L.[3], *Brachypodium distachyon*[4], *Zea mays*[5] | Fluka |
| **Isocitric acid** | C6H8O7 | 192.0270 | Organic acid | *Oryza sativa*[11] | Sigma-Aldrich |
| **Citric acid** | C6H8O7 | 192.0270 | Organic acid | *Triticum aestivum* L.[3], *Zea mays*[5] | Sigma-Aldrich |
| **Ferulic acid** | C10H10O4 | 194.0579 | Organic acid, Phenol | *Hordeum vulgare*[6], *Medicago sativa*[8], *Ageratum conyzoides*[12], *Triticum vulgare*[14], *Avena fatua*[15], *Fagopyrum tataricum*[17] | Fluka |

| | | | | | |
|---|---|---|---|---|---|
| **Syringic acid** | C9H10O5 | 198.0528 | Organic acid, Phenol | *Medicago sativa*[8], *Eucalyptus spp.*[13], *Triticum vulgare*[14], *Avena fatua*[15] | Sigma-Aldrich |
| **Lauric acid** | C12H24O2 | 200.1776 | Fatty acid | *Arabidopsis thaliana*[1], *Lactuca sativa* L.[2] | Sigma-Aldrich |
| **Tryptophan** | C11H12N2O2 | 204.0899 | Amino acid | *Oryza sativa*[11], *Fagopyrum esculentum*[18] | Sigma-Aldrich |
| **Jasmonic acid** | C12H18O3 | 210.1256 | Organic acid | *Triticum aestivum, Avena fatua,* many others[9] | Sigma-Aldrich |
| **N-acetyl-D-mannosamine** | C8H15NO6 | 221.0899 | Sugar | *Arabidopsis thaliana*[1] | Sigma-Aldrich |
| **Cystathionine** | C7H14N2O4S | 222.0674 | Amino acid | *Triticum aestivum* L.[3] | Sigma-Aldrich |
| **3,5-Di-tert-butyl-4-hydroxybenzoic acid** | C15H22O3 | 250.1569 | Organic acid, Phenol | None - synthetic internal standard | Sigma-Aldrich |
| **Palmitic acid** | C16H32O2 | 256.2402 | Fatty acid | *Arabidopsis thaliana*[1], *Fagopyrum esculentum*[19] | Sigma-Aldrich |
| **Stearic acid** | C18H36O2 | 284.2715 | Fatty acid | *Arabidopsis thaliana*[1], *Fagopyrum esculentum*[19] | Sigma-Aldrich |
| **Catechin** | C15H14O6 | 290.0790 | Flavonoid, Phenol | *Fagopyrum esculentum*[20] | Fluka |
| **Epicatechin** | C15H14O6 | 290.0790 | Flavonoid, Phenol | *Fagopyrum tataricum*[17] | Sigma-Aldrich |
| **Quercetin** | C15H10O7 | 302.0427 | Flavonoid, Phenol | *Fagopyrum esculentum*[20] | Sigma-Aldrich |

| | | | | | |
|---|---|---|---|---|---|
| **Arachidic acid** | C20H40O2 | 312.3028 | Fatty acid | *Fagopyrum esculentum*[19] | Sigma-Aldrich |
| **Myricetin** | C15H10O8 | 318.0376 | Flavonoid, Phenol | *Fagopyrum esculentum*[20] | Sigma-Aldrich |
| **Dihexose (sucrose/maltose)** | C12H22O11 | 342.1162 | Sugar | *Arabidopsis thaliana*[1], *Zea mays*[5] *Triticum aestivum* L.[3] | Sigma-Aldrich |
| **α tocopherol** | C29H50O2 | 430.3811 | Lipid, Phenol | *Arabidopsis thaliana*[1] | Sigma-Aldrich |
| **Rutin** | C27H30O16 | 610.1534 | Flavonoid, Phenol, Glycoside | *Fagopyrum tataricum*[17] | Sigma-Aldrich |

7

**Table S2:** Chromatographic gradients & parameters for LC-HRMS/MS data acquisition

**A:** Single column gradient to PFP column

| Time (min) | % A | % B |
|---|---|---|
| 0.50 | 100 | 0 |
| 4.50 | 95 | 5 |
| 5.00 | 60 | 40 |
| 8.00 | 60 | 40 |
| 10.50 | 0 | 100 |
| 12.50 | 0 | 100 |
| 13.50 | 100 | 0 |
| 15.50 | 100 | 0 |

**B:** Dual column pump 1 gradient to PFP column

| Time (min) | % A | % B |
|---|---|---|
| 3.00 | 100 | 0 |
| 7.00 | 95 | 5 |
| 7.50 | 60 | 40 |
| 10.50 | 60 | 40 |
| 13.00 | 0 | 100 |
| 15.00 | 0 | 100 |
| 16.00 | 100 | 0 |
| 20.00 | 100 | 0 |

**C:** Dual column pump 2 gradient PGC column

| Time (min) | % A | % B |
|---|---|---|
| 2.50 | 100 | 0 |
| 6.50 | 40 | 60 |
| 9.00 | 40 | 60 |
| 11.00 | 0 | 100 |
| 13.00 | 0 | 100 |
| 14.00 | 100 | 0 |
| 20.00 | 100 | 0 |

**D:** Other chromatographic and mass spectrometric parameters

| | |
|---|---|
| Single column and dual column pump 1 chromatographic conditions | |
| Stationary phase | Sigma-Aldrich Discovery HS F5<br>(150 x 2.1 mm, 3 µm particle size) |
| Mobile phase A | 99.9% v/v H2O, 0.1% v/v formic acid |
| Mobile phase B | 99.9% v/v methanol, 0.1% v/v formic acid |
| Flow rate (ml min$^{-1}$) | 0.350 |
| Column oven T (°C) | 50 |
| Injection volume (µl) | 5 |
| Dual column pump 2 chromatographic conditions | |
| Stationary phase | Thermo Scientific Hypercarb column (2.1 × 150 mm, 5 µm particle size |
| Mobile phase A | 99.9% v/v H2O, 0.1% v/v formic acid |

| | |
|---|---|
| Mobile phase B | 99.9% v/v acetonitrile, 0.1% v/v formic acid |
| Flow rate (mL $min^{-1}$) | 0.350 |
| Column oven T (°C) | 50 |
| Injection volume (µl) | - |
| Source parameters | |
| Gas Temp (°C) | 300 |
| Gas Flow (L $min^{-1}$) | 8 |
| Nebulizer (psig) | 35 |
| Sheath Gas Temp (L $min^{-1}$) | 300 |
| Sheath Gas Flow (L $min^{-1}$) | 11 |
| VCap | 3500 |
| Nozzle Voltage (V) | 500 |
| Fragmentor (V) | 360 |
| Skimmer 1 | 65 |
| Octopole RF Peak | 750 |
| Scan parameters: Acquisition Mode AutoMS2 (DDA) | |
| MS Min Range (m/z) | 50 |
| MS Max Range (m/z) | 1700 |
| MS Scan Rate (spectra $sec^{-1}$) | 5.00 |
| Resolution MS1 and MS2 | ~17.5 K @ m/z 322 |
| MS/MS Min Range (m/z) | 50 |
| MS/MS Max Range (m/z) | 1700 |
| MS/MS Scan Rate (spectra $sec^{-1}$) | 7.00 |
| Isolation Width MS/MS | Narrow (~1.3 amu) |
| Minimum cycle time (Hz) | 1 |
| Maximum cycle time (Hz) | 5 |
| Collision Energy Table | |
| Mass | Z1 |
| 0 | 5 |
| 300 | 10 |
| 1700 | 45 |
| Precursor Selection | |
| Max Precursors Per Cycle | 5 |
| Threshold (Abs) | 2000 |
| Threshold (Rel %) | 0.010 |
| Precursor abundance based scan speed | No |
| Purity Stringency (%) | 100.000 |
| Purity Cutoff (%) | 30.000 |
| Isotope Model | Common |
| Active exclusion enabled | Yes |
| Active exclusion excluded after (spectra) | 3 |
| Active exclusion released after (min) | 0.30 |
| Sort precursors | By abundance only |
| Reference Masses | |
| Negative ionization | **Positive ionization** |
| 119.0363 m/z | 121.0509m/z |
| 966.0007 m/z | 922.0098m/z |

**Table S3:** MS DIAL preprocessing parameters

| MS-DIAL ver. 4.9.221218 | | |
|---|---|---|
| MS1 Data type | Centroid | |
| MS2 Data type | Centroid | |
| Ion mode | Negative & Positive | |
| Target | Metabolomics | |
| Mode | ddMSMS | |
| **Data collection parameters** | | |
| Retention time begin (min) | Single: 1.6 | Dual: 2.5 |
| Retention time end (min) | Single: 15.5 | Dual: 21.0 |
| Mass range begin (m/z) | 50 | |
| Mass range end (m/z) | 1700 | |
| MS2 mass range begin (m/z) | 50 | |
| MS2 mass range end (m/z) | 1700 | |
| **Centroid parameters** | | |
| MS1 tolerance (Da) | 0.01 | |
| MS2 tolerance (Da) | 0.025 | |
| **Isotope recognition** | | |
| Maximum charged number | 2 | |
| **Peak detection parameters** | | |
| Smoothing method | Linear Weighted Moving Average | |
| Smoothing level (points) | 6 | |
| Minimum peak width (points) | 6 | |
| Minimum peak height (counts) | 1000 | |
| **Peak spotting parameters** | | |
| Mass slice width | 0.1 | |
| **Deconvolution parameters** | | |
| Sigma window value | 0.5 | |
| MS2 Dec amplitude cut off (counts) | 0 | |
| Exclude after precursor | TRUE | |
| Keep isotope until (Da) | 0.5 | |
| Keep original precursor isotopes | FALSE | |
| **MSP file and MS/MS identification setting** | | |
| Retention time tolerance (min) | 1 | |
| Accurate mass tolerance MS1 (Da) | 0.02 | |
| Accurate mass tolerance MS2 (Da) | 0.03 | |
| Identification score cut off (%) | 80 | |
| Using retention time for scoring | TRUE | |
| Using retention time for filtering | FALSE | |
| **Adduct ion setting** | | |
| **Negative ionization** | **Positive ionization** | |
| [M-H]- | [M+H]+ | |

| | |
|---|---|
| [M+Cl]- | [M+Na]+ |
| [M+FA-H]- | [M+K]+ |
| [M-H2O-H]- | [M+NH4]+ |
| [M-CO2-H]- | [M-NH2]+ |
| | [M+H-H2O]+ |
| | [M+H-CO2]+ |
| **Alignment parameters setting** | |
| Retention time tolerance (min) | 0.2 |
| MS1 tolerance (Da) | 0.015 |
| Retention time factor | 0.5 |
| MS1 factor | 0.5 |
| Peak count filter (%) | 0 |
| N detected in at least one group (%) | 60 |
| Remove feature based on peak height fold-change | TRUE |
| Sample average (height, counts) / blank average (height, counts) | 5 |
| Keep identified and annotated metabolites | TRUE |
| Keep removable features and assign the tag for checking | TRUE |
| Gap filling by compulsion | TRUE |

**Table S4:** Compounds upregulated by BO-BO/A when compared to BO-0/A for the second extract of the redroot the P experimental set. For identification level 1, the annotation" w/o MS2" indicates there was not confident spectral match and that the results can not be fully trusted unless corroborated by another source.

| Alignment ID | Ionization Mode | Level 1 Identification | Level 2a Identification | Level 2b Identification | Average Rt(min) | Average Mz | BO-0/A | BO-BO/A | p.value | fdr adjusted p.value | Fold Change | Log2 fold change |
|---|---|---|---|---|---|---|---|---|---|---|---|---|
| 2 | Negative | Unknown | NA | NA | 11.89 | 57.03 | 58.06 | 97.43 | 0.00 | 0.01 | 1.68 | 0.75 |
| 4 | Negative | Unknown | NA | NA | 11.89 | 59.01 | 160.23 | 260.56 | 0.00 | 0.04 | 1.63 | 0.70 |
| 18 | Negative | Unknown | NA | NA | 4.89 | 71.01 | 110.35 | 184.50 | 0.01 | 0.05 | 1.67 | 0.74 |
| 69 | Negative | Unknown | NA | NA | 11.89 | 99.04 | 146.18 | 293.90 | 0.00 | 0.00 | 2.01 | 1.01 |
| 129 | Negative | Unknown | NA | NA | 12.47 | 121.00 | 100.04 | 214.07 | 0.00 | 0.00 | 2.14 | 1.10 |
| 136 | Negative | Unknown | NA | NA | 11.89 | 125.02 | 75.38 | 129.65 | 0.00 | 0.00 | 1.72 | 0.78 |
| 197 | Negative | Unknown | NA | NA | 9.13 | 150.06 | 17.08 | 28.35 | 0.01 | 0.03 | 1.66 | 0.73 |
| 221 | Negative | Unknown | NA | NA | 12.38 | 162.02 | 43.11 | 81.18 | 0.00 | 0.00 | 1.88 | 0.91 |
| 237 | Negative | w/o MS2:Vanilic acid | NA | NA | 10.47 | 167.04 | 7.36 | 12.26 | 0.01 | 0.02 | 1.67 | 0.74 |
| 287 | Negative | Unknown | NA | L-Tyrosine | 6.87 | 180.07 | 74.97 | 134.94 | 0.01 | 0.04 | 1.80 | 0.85 |
| 322 | Negative | Unknown | D-(-)-Quinic acid; LC-ESI-QTOF; MS2; CE | Quinic acid | 5.18 | 191.06 | 6753.04 | 12377.94 | 0.01 | 0.04 | 1.83 | 0.87 |
| 335 | Negative | Unknown | NA | NA | 4.70 | 195.05 | 102.23 | 176.19 | 0.00 | 0.01 | 1.72 | 0.79 |
| 336 | Negative | Unknown | 1726059 - 40.0 eV | Galactonic acid | 4.98 | 195.05 | 559.81 | 1292.16 | 0.00 | 0.02 | 2.31 | 1.21 |
| 414 | Negative | w/o MS2:N-acetyl-D-mannosamine | NA | NA | 5.09 | 220.08 | 395.67 | 1025.70 | 0.00 | 0.00 | 2.59 | 1.37 |
| 425 | Negative | Unknown | NA | NA | 2.90 | 224.09 | 45.35 | 115.01 | 0.01 | 0.01 | 2.54 | 1.34 |
| 429 | Negative | Unknown | NA | NA | 4.89 | 227.00 | 53.70 | 89.44 | 0.00 | 0.02 | 1.67 | 0.74 |
| 508 | Negative | Unknown | NA | NA | 4.99 | 253.09 | 72.82 | 182.99 | 0.00 | 0.01 | 2.51 | 1.33 |
| 528 | Negative | Unknown | NA | NA | 4.53 | 260.02 | 75.19 | 136.47 | 0.00 | 0.01 | 1.82 | 0.86 |
| 536 | Negative | Unknown | NA | NA | 10.69 | 263.02 | 65.82 | 103.89 | 0.00 | 0.01 | 1.58 | 0.66 |
| 544 | Negative | Unknown | NA | NA | 9.02 | 267.03 | 200.07 | 339.10 | 0.01 | 0.03 | 1.69 | 0.76 |
| 604 | Negative | Unknown | NA | NA | 2.94 | 287.02 | 30.47 | 94.34 | 0.01 | 0.04 | 3.10 | 1.63 |
| 622 | Negative | Unknown | NA | NA | 9.39 | 293.09 | 270.47 | 466.67 | 0.01 | 0.03 | 1.73 | 0.79 |
| 662 | Negative | Unknown | NA | NA | 12.40 | 307.12 | 205.07 | 330.74 | 0.01 | 0.03 | 1.61 | 0.69 |
| 688 | Negative | Unknown | NA | NA | 3.51 | 315.08 | 13.81 | 50.08 | 0.00 | 0.02 | 3.63 | 1.86 |
| 704 | Negative | Unknown | NA | NA | 12.47 | 321.10 | 514.21 | 1126.01 | 0.00 | 0.00 | 2.19 | 1.13 |
| 731 | Negative | Unknown | NA | NA | 3.53 | 331.05 | 97.27 | 262.34 | 0.01 | 0.05 | 2.70 | 1.43 |
| 741 | Negative | Unknown | NA | NA | 4.68 | 334.13 | 11.45 | 21.02 | 0.00 | 0.00 | 1.84 | 0.88 |
| 749 | Negative | Unknown | NA | NA | 9.35 | 337.11 | 152.03 | 248.11 | 0.00 | 0.01 | 1.63 | 0.71 |
| 772 | Negative | Unknown | NA | NA | 11.38 | 343.15 | 48.44 | 103.60 | 0.00 | 0.00 | 2.14 | 1.10 |
| 776 | Negative | Unknown | NA | NA | 2.89 | 344.96 | 132.69 | 265.58 | 0.02 | 0.05 | 2.00 | 1.00 |
| 787 | Negative | Unknown | NA | NA | 12.68 | 349.19 | 135.45 | 295.56 | 0.01 | 0.03 | 2.18 | 1.13 |
| 798 | Negative | Unknown | NA | NA | 11.38 | 353.18 | 81.34 | 184.05 | 0.00 | 0.01 | 2.26 | 1.18 |
| 802 | Negative | Unknown | NA | NA | 9.85 | 355.05 | 37.50 | 70.00 | 0.00 | 0.01 | 1.87 | 0.90 |
| 826 | Negative | Unknown | NA | NA | 12.55 | 365.05 | 69.51 | 116.90 | 0.01 | 0.02 | 1.68 | 0.75 |
| 829 | Negative | Unknown | NA | NA | 11.47 | 365.14 | 137.07 | 247.62 | 0.00 | 0.01 | 1.81 | 0.85 |
| 839 | Negative | Unknown | NA | NA | 9.08 | 371.10 | 158.97 | 265.01 | 0.01 | 0.02 | 1.67 | 0.74 |
| 843 | Negative | Unknown | NA | NA | 9.03 | 373.08 | 3535.57 | 5654.89 | 0.00 | 0.01 | 1.60 | 0.68 |
| 866 | Negative | Unknown | NA | NA | 10.32 | 379.16 | 258.68 | 426.10 | 0.01 | 0.02 | 1.65 | 0.72 |
| 874 | Negative | Unknown | NA | NA | 5.17 | 383.12 | 65.12 | 177.47 | 0.01 | 0.03 | 2.73 | 1.45 |
| 888 | Negative | Unknown | NA | NA | 11.68 | 387.09 | 51.25 | 93.17 | 0.00 | 0.00 | 1.82 | 0.86 |
| 889 | Negative | Unknown | NA | NA | 9.15 | 387.10 | 588.64 | 991.05 | 0.01 | 0.02 | 1.68 | 0.75 |

| | | | | | | | | | | | | |
|---|---|---|---|---|---|---|---|---|---|---|---|---|
| 891 | Negative | Unknown | alpha,alpha-Trehalose - 40.0 eV | NA | 4.98 | 387.12 | 1949.15 | 3256.92 | 0.01 | 0.04 | 1.67 | 0.74 |
| 893 | Negative | Unknown | NA | NA | 9.39 | 387.13 | 1073.43 | 1945.77 | 0.01 | 0.02 | 1.81 | 0.86 |
| 917 | Negative | Unknown | NA | NA | 11.54 | 393.18 | 78.41 | 157.94 | 0.01 | 0.02 | 2.01 | 1.01 |
| 934 | Negative | Unknown | NA | NA | 9.03 | 403.08 | 41.92 | 95.15 | 0.00 | 0.02 | 2.27 | 1.18 |
| 935 | Negative | Unknown | NA | PubChem:(46222441) | 8.49 | 403.09 | 144.79 | 296.08 | 0.02 | 0.04 | 2.04 | 1.03 |
| 942 | Negative | Unknown | NA | NA | 9.03 | 405.05 | 41.70 | 71.12 | 0.00 | 0.01 | 1.71 | 0.77 |
| 960 | Negative | Unknown | NA | COCONUT:(CNP0159349 CNP0218833);Natural Products:(UNPD202200);PubChem:(38363079 38363084 45360328 44715457 125416071 125416072 125416073 125416074);SuperNatural:(SN00032625 SN00030299 SN00032624 SN00030300 SN00032623 SN00032622);ZINC bio:(ZINC31169353 ZINC31169357 ZINC35454681 ZINC35454685 ZINC35454688 ZINC35454690);Training Set | 11.89 | 413.15 | 4009.66 | 7535.58 | 0.00 | 0.01 | 1.88 | 0.91 |
| 961 | Negative | Unknown | NA | COCONUT:(CNP0159349 CNP0218833);Natural Products:(UNPD202200);PubChem:(38363079 38363084 45360328 44715457 125416071 125416072 125416073 125416074);SuperNatural:(SN00032625 SN00030299 SN00032624 SN00030300 SN00032623 SN00032622);ZINC bio:(ZINC31169353 ZINC31169357 ZINC35454681 ZINC35454685 ZINC35454688 ZINC35454690);Training Set | 11.06 | 413.15 | 60.28 | 112.55 | 0.00 | 0.01 | 1.87 | 0.90 |
| 962 | Negative | Unknown | NA | NA | 10.55 | 413.15 | 35.24 | 61.29 | 0.00 | 0.00 | 1.74 | 0.80 |
| 966 | Negative | Unknown | NA | NA | 5.20 | 415.14 | 306.35 | 671.59 | 0.00 | 0.01 | 2.19 | 1.13 |
| 972 | Negative | Unknown | NA | NA | 9.08 | 417.11 | 159.29 | 266.93 | 0.00 | 0.01 | 1.68 | 0.74 |
| 988 | Negative | Unknown | NA | NA | 9.15 | 421.08 | 84.13 | 160.65 | 0.00 | 0.01 | 1.91 | 0.93 |
| 1013 | Negative | Unknown | NA | NA | 9.20 | 429.14 | 15.87 | 32.41 | 0.01 | 0.01 | 2.04 | 1.03 |
| 1018 | Negative | Unknown | NA | NA | 12.59 | 431.10 | 42.92 | 78.13 | 0.01 | 0.03 | 1.82 | 0.86 |
| 1034 | Negative | Unknown | NA | NA | 12.71 | 435.19 | 937.41 | 1936.53 | 0.00 | 0.01 | 2.07 | 1.05 |
| 1061 | Negative | Unknown | NA | NA | 9.40 | 447.11 | 17.13 | 34.28 | 0.02 | 0.04 | 2.00 | 1.00 |
| 1064 | Negative | Unknown | NA | NA | 9.81 | 447.15 | 458.97 | 782.87 | 0.02 | 0.04 | 1.71 | 0.77 |
| 1065 | Negative | Unknown | NA | NA | 9.15 | 447.15 | 267.19 | 492.42 | 0.01 | 0.02 | 1.84 | 0.88 |
| 1070 | Negative | Unknown | NA | NA | 12.31 | 449.20 | 106.14 | 190.03 | 0.00 | 0.00 | 1.79 | 0.84 |
| 1071 | Negative | Unknown | NA | NA | 10.98 | 449.20 | 50.47 | 85.95 | 0.00 | 0.01 | 1.70 | 0.77 |
| 1079 | Negative | Unknown | NA | NA | 12.94 | 451.22 | 366.56 | 964.78 | 0.00 | 0.02 | 2.63 | 1.40 |
| 1085 | Negative | Unknown | NA | NA | 11.01 | 453.20 | 45.33 | 121.38 | 0.00 | 0.02 | 2.68 | 1.42 |
| 1088 | Negative | Unknown | NA | NA | 9.93 | 455.12 | 120.04 | 186.29 | 0.01 | 0.04 | 1.55 | 0.63 |
| 1089 | Negative | Unknown | NA | NA | 9.47 | 455.12 | 547.17 | 960.46 | 0.01 | 0.03 | 1.76 | 0.81 |
| 1092 | Negative | Unknown | NA | NA | 12.86 | 456.15 | 83.74 | 129.58 | 0.00 | 0.01 | 1.55 | 0.63 |
| 1094 | Negative | Unknown | NA | NA | 12.51 | 457.17 | 268.40 | 454.43 | 0.00 | 0.01 | 1.69 | 0.76 |
| 1105 | Negative | Unknown | NA | NA | 9.50 | 461.13 | 36.49 | 64.02 | 0.01 | 0.04 | 1.75 | 0.81 |
| 1113 | Negative | Unknown | NA | NA | 9.54 | 463.14 | 139.78 | 232.06 | 0.01 | 0.04 | 1.66 | 0.73 |
| 1119 | Negative | Unknown | NA | NA | 13.13 | 465.25 | 86.96 | 164.57 | 0.01 | 0.02 | 1.89 | 0.92 |
| 1123 | Negative | Unknown | NA | NA | 9.11 | 467.05 | 30.30 | 54.66 | 0.00 | 0.01 | 1.80 | 0.85 |
| 1125 | Negative | Unknown | NA | NA | 12.77 | 467.21 | 416.56 | 807.86 | 0.01 | 0.01 | 1.94 | 0.96 |
| 1135 | Negative | Unknown | NA | NA | 10.42 | 472.10 | 35.86 | 59.37 | 0.02 | 0.03 | 1.66 | 0.73 |
| 1137 | Negative | Unknown | NA | NA | 10.34 | 473.17 | 69.22 | 112.64 | 0.01 | 0.04 | 1.63 | 0.70 |
| 1142 | Negative | Unknown | NA | NA | 11.90 | 476.14 | 54.78 | 111.44 | 0.00 | 0.01 | 2.03 | 1.02 |
| 1146 | Negative | Unknown | NA | NA | 12.64 | 477.14 | 43.37 | 70.95 | 0.02 | 0.05 | 1.64 | 0.71 |
| 1151 | Negative | Unknown | NA | NA | 10.32 | 479.09 | 42.79 | 92.43 | 0.02 | 0.04 | 2.16 | 1.11 |
| 1158 | Negative | Unknown | NA | NA | 12.51 | 482.19 | 58.08 | 88.28 | 0.01 | 0.03 | 1.52 | 0.60 |
| 1161 | Negative | Unknown | NA | NA | 10.12 | 483.21 | 69.55 | 155.71 | 0.00 | 0.02 | 2.24 | 1.16 |
| 1191 | Negative | Unknown | NA | NA | 10.04 | 493.16 | 4.59 | 13.26 | 0.01 | 0.03 | 2.89 | 1.53 |
| 1193 | Negative | Unknown | NA | NA | 13.21 | 493.23 | 213.84 | 430.17 | 0.01 | 0.03 | 2.01 | 1.01 |

| | | | | | | | | | | | | |
|---|---|---|---|---|---|---|---|---|---|---|---|---|
| 1206 | Negative | Unknown | NA | NA | 5.32 | 503.16 | 2704.38 | 5838.80 | 0.00 | 0.01 | 2.16 | 1.11 |
| 1207 | Negative | Unknown | NA | NA | 11.97 | 503.17 | 36.05 | 78.30 | 0.00 | 0.01 | 2.17 | 1.12 |
| 1214 | Negative | Unknown | NA | NA | 12.58 | 507.21 | 1545.30 | 3292.72 | 0.00 | 0.00 | 2.13 | 1.09 |
| 1220 | Negative | Unknown | NA | NA | 12.77 | 511.11 | 99.32 | 185.49 | 0.00 | 0.01 | 1.87 | 0.90 |
| 1225 | Negative | Unknown | NA | NA | 11.88 | 513.07 | 661.15 | 1442.13 | 0.01 | 0.03 | 2.18 | 1.13 |
| 1230 | Negative | Unknown | NA | NA | 4.96 | 515.12 | 25.10 | 61.98 | 0.00 | 0.01 | 2.47 | 1.30 |
| 1242 | Negative | Unknown | NA | NA | 13.28 | 519.24 | 52.95 | 103.99 | 0.01 | 0.04 | 1.96 | 0.97 |
| 1249 | Negative | Unknown | NA | NA | 9.23 | 523.17 | 48.24 | 81.29 | 0.01 | 0.04 | 1.69 | 0.75 |
| 1258 | Negative | Unknown | NA | NA | 10.36 | 533.16 | 46.32 | 89.14 | 0.00 | 0.00 | 1.92 | 0.94 |
| 1259 | Negative | Unknown | NA | NA | 9.76 | 533.17 | 261.98 | 505.07 | 0.01 | 0.01 | 1.93 | 0.95 |
| 1260 | Negative | Unknown | NA | NA | 5.16 | 533.17 | 509.46 | 1484.26 | 0.00 | 0.00 | 2.91 | 1.54 |
| 1268 | Negative | Unknown | NA | NA | 13.45 | 537.25 | 131.66 | 224.45 | 0.01 | 0.03 | 1.70 | 0.77 |
| 1310 | Negative | Unknown | NA | NA | 9.81 | 561.22 | 40.30 | 65.14 | 0.01 | 0.01 | 1.62 | 0.69 |
| 1311 | Negative | Unknown | NA | NA | 5.10 | 562.20 | 41.51 | 127.99 | 0.00 | 0.01 | 3.08 | 1.62 |
| 1329 | Negative | Unknown | NA | NA | 11.28 | 575.20 | 58.16 | 100.75 | 0.00 | 0.00 | 1.73 | 0.79 |
| 1339 | Negative | Unknown | NA | NA | 10.63 | 589.18 | 43.81 | 69.09 | 0.01 | 0.02 | 1.58 | 0.66 |
| 1348 | Negative | Unknown | NA | NA | 5.32 | 601.13 | 264.42 | 568.59 | 0.01 | 0.04 | 2.15 | 1.10 |
| 1356 | Negative | Unknown | NA | NA | 5.20 | 607.21 | 29.37 | 107.10 | 0.01 | 0.02 | 3.65 | 1.87 |
| 1370 | Negative | Unknown | NA | NA | 5.32 | 621.19 | 52.46 | 111.66 | 0.01 | 0.02 | 2.13 | 1.09 |
| 1390 | Negative | Unknown | NA | NA | 5.49 | 637.18 | 721.09 | 1538.39 | 0.00 | 0.02 | 2.13 | 1.09 |
| 1396 | Negative | Unknown | NA | NA | 13.58 | 641.19 | 5.57 | 9.07 | 0.01 | 0.03 | 1.63 | 0.71 |
| 1399 | Negative | Unknown | NA | NA | 13.05 | 643.20 | 85.85 | 137.10 | 0.00 | 0.00 | 1.60 | 0.68 |
| 1414 | Negative | Unknown | NA | NA | 13.19 | 661.18 | 147.50 | 227.05 | 0.01 | 0.02 | 1.54 | 0.62 |
| 1415 | Negative | Unknown | NA | NA | 12.63 | 661.18 | 54.26 | 129.41 | 0.00 | 0.00 | 2.39 | 1.25 |
| 1432 | Negative | Unknown | NA | NA | 8.03 | 668.20 | 29.69 | 69.56 | 0.02 | 0.04 | 2.34 | 1.23 |
| 1446 | Negative | Unknown | NA | NA | 13.56 | 675.19 | 188.99 | 320.54 | 0.00 | 0.00 | 1.70 | 0.76 |
| 1447 | Negative | Unknown | NA | NA | 13.27 | 675.19 | 759.05 | 1199.50 | 0.00 | 0.01 | 1.58 | 0.66 |
| 1451 | Negative | Unknown | NA | NA | 5.50 | 679.22 | 21.19 | 88.37 | 0.00 | 0.02 | 4.17 | 2.06 |
| 1458 | Negative | Unknown | NA | NA | 5.09 | 683.23 | 3001.75 | 7380.36 | 0.00 | 0.03 | 2.46 | 1.30 |
| 1461 | Negative | Unknown | NA | NA | 13.26 | 685.18 | 216.15 | 371.79 | 0.00 | 0.00 | 1.72 | 0.78 |
| 1466 | Negative | Unknown | NA | NA | 11.25 | 687.19 | 102.71 | 209.85 | 0.00 | 0.00 | 2.04 | 1.03 |
| 1467 | Negative | Unknown | NA | NA | 13.05 | 687.19 | 1336.43 | 2259.57 | 0.00 | 0.00 | 1.69 | 0.76 |
| 1486 | Negative | Unknown | NA | NA | 12.60 | 697.24 | 108.02 | 222.64 | 0.00 | 0.01 | 2.06 | 1.04 |
| 1492 | Negative | Unknown | NA | NA | 12.87 | 703.19 | 158.86 | 246.80 | 0.01 | 0.02 | 1.55 | 0.64 |
| 1494 | Negative | Unknown | NA | NA | 10.01 | 703.26 | 30.60 | 65.33 | 0.01 | 0.02 | 2.14 | 1.09 |
| 1497 | Negative | Unknown | NA | NA | 10.27 | 707.18 | 147.90 | 244.72 | 0.01 | 0.02 | 1.65 | 0.73 |
| 1507 | Negative | Unknown | NA | NA | 11.88 | 711.30 | 25.52 | 49.32 | 0.00 | 0.00 | 1.93 | 0.95 |
| 1539 | Negative | Unknown | NA | NA | 13.05 | 733.20 | 58.43 | 92.88 | 0.00 | 0.01 | 1.59 | 0.67 |
| 1547 | Negative | Unknown | NA | NA | 13.27 | 738.19 | 87.08 | 136.79 | 0.01 | 0.02 | 1.57 | 0.65 |
| 1610 | Negative | Unknown | NA | NA | 5.09 | 781.20 | 60.68 | 147.66 | 0.01 | 0.05 | 2.43 | 1.28 |
| 1611 | Negative | Unknown | NA | NA | 12.88 | 781.22 | 53.35 | 88.52 | 0.00 | 0.01 | 1.66 | 0.73 |
| 1623 | Negative | Unknown | NA | NA | 10.41 | 795.29 | 19.72 | 43.56 | 0.01 | 0.03 | 2.21 | 1.14 |
| 1633 | Negative | Unknown | NA | NA | 13.05 | 805.22 | 83.74 | 132.19 | 0.00 | 0.00 | 1.58 | 0.66 |
| 1635 | Negative | Unknown | NA | NA | 11.08 | 808.21 | 67.59 | 158.86 | 0.01 | 0.02 | 2.35 | 1.23 |
| 1636 | Negative | Unknown | NA | NA | 12.57 | 808.21 | 44.65 | 79.70 | 0.03 | 0.05 | 1.79 | 0.84 |
| 1664 | Negative | Unknown | NA | NA | 10.03 | 826.30 | 26.33 | 47.14 | 0.02 | 0.05 | 1.79 | 0.84 |
| 1691 | Negative | Unknown | NA | NA | 12.58 | 845.45 | 54.65 | 87.43 | 0.02 | 0.04 | 1.60 | 0.68 |
| 1697 | Negative | Unknown | NA | NA | 12.64 | 857.25 | 83.52 | 155.23 | 0.00 | 0.01 | 1.86 | 0.89 |
| 1730 | Negative | Unknown | NA | NA | 13.33 | 943.29 | 226.06 | 416.94 | 0.01 | 0.02 | 1.84 | 0.88 |
| 1764 | Negative | Unknown | NA | NA | 14.65 | 985.61 | 16.25 | 43.33 | 0.01 | 0.04 | 2.67 | 1.42 |
| 1788 | Negative | Unknown | NA | NA | 14.69 | 1033.54 | 20.62 | 43.43 | 0.01 | 0.03 | 2.11 | 1.07 |
| 1808 | Negative | Unknown | NA | NA | 13.25 | 1105.34 | 58.88 | 94.58 | 0.01 | 0.04 | 1.61 | 0.68 |
| 1809 | Negative | Unknown | NA | NA | 13.09 | 1109.41 | 64.05 | 140.99 | 0.00 | 0.00 | 2.20 | 1.14 |
| 1825 | Negative | Unknown | NA | NA | 13.05 | 1375.39 | 37.25 | 89.75 | 0.00 | 0.00 | 2.41 | 1.27 |
| 180 | Positive | Unknown | NA | NA | 5.03 | 110.07 | 115.79 | 298.11 | 0.00 | 0.02 | 2.57 | 1.36 |

| | | | | | | | | | | | | |
|---|---|---|---|---|---|---|---|---|---|---|---|---|
| 216 | Positive | Unknown | NA | NA | 4.88 | 120.07 | 32.56 | 54.98 | 0.01 | 0.04 | 1.69 | 0.76 |
| 273 | Positive | Unknown | NA | NA | 4.70 | 138.05 | 29.62 | 58.68 | 0.01 | 0.02 | 1.98 | 0.99 |
| 274 | Positive | Unknown | NA | 2-Aminobenzoic acid | 4.97 | 138.05 | 650.74 | 1316.18 | 0.01 | 0.02 | 2.02 | 1.02 |
| 296 | Positive | Unknown | NA | NA | 11.88 | 145.05 | 67.62 | 114.20 | 0.00 | 0.01 | 1.69 | 0.76 |
| 308 | Positive | Unknown | NA | NA | 9.67 | 149.12 | 28.11 | 68.45 | 0.00 | 0.02 | 2.44 | 1.28 |
| 343 | Positive | Unknown | NA | NA | 3.69 | 160.10 | 32.95 | 58.40 | 0.01 | 0.03 | 1.77 | 0.83 |
| 352 | Positive | Unknown | NA | NA | 4.96 | 163.06 | 115.47 | 229.01 | 0.00 | 0.05 | 1.98 | 0.99 |
| 353 | Positive | Unknown | NA | NA | 4.86 | 163.06 | 342.25 | 638.95 | 0.00 | 0.04 | 1.87 | 0.90 |
| 354 | Positive | Unknown | NA | NA | 11.87 | 163.06 | 95.10 | 165.21 | 0.00 | 0.01 | 1.74 | 0.80 |
| 363 | Positive | Unknown | NA | Sabiden | 10.50 | 166.09 | 810.86 | 1308.36 | 0.00 | 0.03 | 1.61 | 0.69 |
| 386 | Positive | Unknown | NA | NA | 4.99 | 176.09 | 120.45 | 266.88 | 0.00 | 0.00 | 2.22 | 1.15 |
| 405 | Positive | Unknown | NA | NA | 5.03 | 181.05 | 40.14 | 84.83 | 0.00 | 0.01 | 2.11 | 1.08 |
| 422 | Positive | Unknown | NA | NA | 3.54 | 185.04 | 26.28 | 51.47 | 0.01 | 0.03 | 1.96 | 0.97 |
| 423 | Positive | Unknown | NA | NA | 5.02 | 185.09 | 42.34 | 70.69 | 0.01 | 0.03 | 1.67 | 0.74 |
| 426 | Positive | Unknown | NA | NA | 5.00 | 186.08 | 195.77 | 366.07 | 0.00 | 0.00 | 1.87 | 0.90 |
| 445 | Positive | Unknown | NA | NA | 4.76 | 192.09 | 77.68 | 170.14 | 0.01 | 0.02 | 2.19 | 1.13 |
| 466 | Positive | Unknown | NA | NA | 4.72 | 199.06 | 42.52 | 87.68 | 0.00 | 0.03 | 2.06 | 1.04 |
| 490 | Positive | Unknown | NA | NA | 4.94 | 204.09 | 653.25 | 1377.17 | 0.00 | 0.01 | 2.11 | 1.08 |
| 509 | Positive | Unknown | NA | NA | 5.02 | 211.11 | 46.61 | 81.81 | 0.01 | 0.04 | 1.76 | 0.81 |
| 518 | Positive | Unknown | NA | NA | 11.31 | 215.14 | 70.94 | 107.73 | 0.01 | 0.05 | 1.52 | 0.60 |
| 535 | Positive | Unknown | NA | NA | 5.02 | 221.11 | 66.25 | 110.46 | 0.01 | 0.03 | 1.67 | 0.74 |
| 538 | Positive | Unknown | NA | N-Acetylgalactosamine | 4.99 | 222.10 | 3804.84 | 8875.24 | 0.00 | 0.00 | 2.33 | 1.22 |
| 587 | Positive | Unknown | NA | NA | 4.84 | 235.09 | 69.10 | 116.87 | 0.01 | 0.03 | 1.69 | 0.76 |
| 597 | Positive | Unknown | NA | NA | 3.45 | 238.56 | 15.55 | 44.66 | 0.00 | 0.03 | 2.87 | 1.52 |
| 613 | Positive | Unknown | NA | NA | 6.03 | 244.08 | 66.88 | 134.31 | 0.02 | 0.03 | 2.01 | 1.01 |
| 639 | Positive | Unknown | NA | NA | 4.88 | 249.11 | 127.11 | 214.50 | 0.00 | 0.02 | 1.69 | 0.75 |
| 698 | Positive | Unknown | NA | NA | 11.48 | 267.10 | 41.02 | 75.46 | 0.00 | 0.00 | 1.84 | 0.88 |
| 707 | Positive | Unknown | NA | NA | 5.02 | 269.11 | 1259.20 | 2216.33 | 0.01 | 0.04 | 1.76 | 0.82 |
| 713 | Positive | Unknown | NA | NA | 10.93 | 270.13 | 572.20 | 1065.14 | 0.00 | 0.00 | 1.86 | 0.90 |
| 718 | Positive | Unknown | NA | NA | 5.14 | 272.07 | 74.24 | 170.69 | 0.00 | 0.01 | 2.30 | 1.20 |
| 720 | Positive | Unknown | NA | NA | 5.33 | 272.07 | 74.96 | 156.89 | 0.01 | 0.04 | 2.09 | 1.07 |
| 737 | Positive | Unknown | NA | NA | 5.03 | 277.14 | 47.59 | 94.01 | 0.01 | 0.03 | 1.98 | 0.98 |
| 759 | Positive | Unknown | NA | NA | 9.23 | 284.08 | 26.68 | 63.02 | 0.00 | 0.02 | 2.36 | 1.24 |
| 779 | Positive | Unknown | NA | NA | 2.92 | 289.03 | 22.89 | 66.64 | 0.02 | 0.05 | 2.91 | 1.54 |
| 799 | Positive | Unknown | NA | NA | 4.79 | 293.14 | 36.93 | 146.78 | 0.00 | 0.01 | 3.97 | 1.99 |
| 807 | Positive | Unknown | NA | NA | 9.40 | 295.10 | 62.11 | 109.32 | 0.01 | 0.03 | 1.76 | 0.82 |
| 817 | Positive | Unknown | NA | NA | 12.29 | 298.10 | 48.87 | 86.34 | 0.00 | 0.04 | 1.77 | 0.82 |
| 835 | Positive | Unknown | NA | NA | 4.74 | 305.13 | 63.59 | 129.55 | 0.00 | 0.02 | 2.04 | 1.03 |
| 844 | Positive | Unknown | NA | NA | 11.87 | 307.10 | 145.62 | 258.07 | 0.00 | 0.01 | 1.77 | 0.83 |
| 864 | Positive | Unknown | NA | NA | 14.74 | 313.27 | 60.83 | 113.71 | 0.01 | 0.02 | 1.87 | 0.90 |
| 886 | Positive | Unknown | NA | NA | 5.02 | 323.15 | 22264.91 | 39969.24 | 0.01 | 0.04 | 1.80 | 0.84 |
| 908 | Positive | Unknown | NA | NA | 5.02 | 332.07 | 80.58 | 175.90 | 0.01 | 0.03 | 2.18 | 1.13 |
| 923 | Positive | Unknown | NA | Asn-agam | 4.61 | 336.14 | 7.66 | 13.42 | 0.03 | 0.05 | 1.75 | 0.81 |
| 947 | Positive | Unknown | NA | NA | 5.01 | 347.10 | 28.05 | 70.64 | 0.01 | 0.02 | 2.52 | 1.33 |
| 973 | Positive | Unknown | NA | NA | 5.23 | 353.09 | 30.92 | 65.03 | 0.01 | 0.04 | 2.10 | 1.07 |
| 974 | Positive | Unknown | NA | NA | 5.50 | 353.09 | 35.21 | 82.76 | 0.00 | 0.01 | 2.35 | 1.23 |
| 990 | Positive | Unknown | Spectral Match to D-(+)-Trehalose from NIST14 | NA | 4.81 | 360.15 | 40.17 | 105.34 | 0.00 | 0.03 | 2.62 | 1.39 |
| 993 | Positive | Unknown | NA | NA | 4.92 | 362.10 | 140.06 | 284.41 | 0.00 | 0.02 | 2.03 | 1.02 |
| 1000 | Positive | Dihexose (Maltose/Sucrose) | NA | NA | 4.92 | 365.11 | 123.01 | 247.26 | 0.00 | 0.02 | 2.01 | 1.01 |
| 1026 | Positive | Unknown | NA | NA | 5.02 | 376.06 | 32.45 | 71.82 | 0.01 | 0.03 | 2.21 | 1.15 |
| 1045 | Positive | Unknown | NA | NA | 4.91 | 381.08 | 37.34 | 81.12 | 0.00 | 0.04 | 2.17 | 1.12 |
| 1059 | Positive | Unknown | NA | NA | 4.92 | 385.09 | 19.29 | 48.46 | 0.00 | 0.02 | 2.51 | 1.33 |

| | | | | | | | | | | | | |
|---|---|---|---|---|---|---|---|---|---|---|---|---|
| 1072 | Positive | Unknown | NA | NA | 10.91 | 390.10 | 36.84 | 70.32 | 0.03 | 0.05 | 1.91 | 0.93 |
| 1089 | Positive | Unknown | NA | NA | 11.87 | 397.15 | 183.44 | 288.61 | 0.01 | 0.03 | 1.57 | 0.65 |
| 1111 | Positive | Unknown | NA | NA | 9.39 | 406.17 | 226.51 | 385.60 | 0.01 | 0.02 | 1.70 | 0.77 |
| 1174 | Positive | Unknown | NA | NA | 11.87 | 432.19 | 222.83 | 362.44 | 0.01 | 0.03 | 1.63 | 0.70 |
| 1187 | Positive | Unknown | NA | NA | 11.87 | 437.14 | 217.74 | 389.54 | 0.00 | 0.01 | 1.79 | 0.84 |
| 1241 | Positive | Unknown | NA | NA | 11.89 | 453.12 | 57.99 | 97.38 | 0.01 | 0.03 | 1.68 | 0.75 |
| 1255 | Positive | Unknown | NA | NA | 12.86 | 458.17 | 78.68 | 139.13 | 0.00 | 0.01 | 1.77 | 0.82 |
| 1280 | Positive | Unknown | NA | NA | 9.80 | 466.19 | 135.08 | 228.86 | 0.01 | 0.05 | 1.69 | 0.76 |
| 1336 | Positive | Unknown | NA | NA | 14.68 | 487.27 | 62.81 | 139.42 | 0.01 | 0.02 | 2.22 | 1.15 |
| 1397 | Positive | Unknown | NA | NA | 12.57 | 509.22 | 185.32 | 286.65 | 0.01 | 0.04 | 1.55 | 0.63 |
| 1409 | Positive | Unknown | NA | NA | 13.06 | 515.15 | 41.45 | 65.24 | 0.00 | 0.00 | 1.57 | 0.65 |
| 1430 | Positive | Unknown | Melezitose | NA | 5.14 | 522.20 | 20.53 | 75.18 | 0.00 | 0.00 | 3.66 | 1.87 |
| 1436 | Positive | Unknown | NA | NA | 5.15 | 524.15 | 5.41 | 59.91 | 0.00 | 0.01 | 11.07 | 3.47 |
| 1445 | Positive | Unknown | NA | NA | 5.17 | 527.16 | 20.64 | 69.46 | 0.00 | 0.00 | 3.36 | 1.75 |
| 1461 | Positive | Unknown | NA | NA | 4.92 | 533.16 | 53.45 | 197.84 | 0.00 | 0.03 | 3.70 | 1.89 |
| 1578 | Positive | Unknown | NA | NA | 4.43 | 577.18 | 26.55 | 53.57 | 0.02 | 0.03 | 2.02 | 1.01 |
| 1776 | Positive | Unknown | NA | NA | 13.19 | 663.19 | 152.74 | 244.75 | 0.00 | 0.01 | 1.60 | 0.68 |
| 1807 | Positive | Unknown | NA | NA | 13.28 | 677.21 | 845.27 | 1365.83 | 0.00 | 0.01 | 1.62 | 0.69 |
| 1808 | Positive | Unknown | NA | NA | 13.57 | 677.21 | 150.68 | 248.09 | 0.00 | 0.01 | 1.65 | 0.72 |
| 1834 | Positive | Unknown | NA | NA | 13.06 | 689.21 | 139.98 | 215.93 | 0.01 | 0.02 | 1.54 | 0.63 |
| 1879 | Positive | Unknown | NA | NA | 4.92 | 704.21 | 10.77 | 68.17 | 0.01 | 0.03 | 6.33 | 2.66 |
| 1934 | Positive | Unknown | NA | NA | 12.88 | 726.26 | 44.01 | 76.00 | 0.01 | 0.04 | 1.73 | 0.79 |
| 1995 | Positive | Unknown | NA | NA | 11.45 | 751.19 | 16.74 | 35.69 | 0.02 | 0.05 | 2.13 | 1.09 |
| 2033 | Positive | Unknown | NA | NA | 14.76 | 772.59 | 31.28 | 69.80 | 0.01 | 0.03 | 2.23 | 1.16 |
| 2045 | Positive | Unknown | Spectral Match to 1,2-Dilinoleoyl-sn-glycero-3-phosphocholine from NIST14 | NA | 15.21 | 782.57 | 130.59 | 320.48 | 0.02 | 0.04 | 2.45 | 1.30 |
| 2138 | Positive | Unknown | NA | NA | 14.38 | 916.51 | 45.40 | 83.98 | 0.01 | 0.02 | 1.85 | 0.89 |
| 2152 | Positive | Unknown | NA | NA | 14.68 | 934.65 | 35.41 | 86.50 | 0.01 | 0.04 | 2.44 | 1.29 |
| 2165 | Positive | Unknown | NA | NA | 13.34 | 962.33 | 117.44 | 191.73 | 0.01 | 0.02 | 1.63 | 0.71 |
| 2166 | Positive | Unknown | NA | NA | 14.65 | 963.60 | 17.34 | 46.50 | 0.01 | 0.04 | 2.68 | 1.42 |

**Table S5:** Compounds upregulated by BO-BO/A when compared to BO-0/A for the second extract of the G experimental set. For identification level 1, the annotation" w/o MS2" indicates there was not confident spectral match and that the results can not be fully trusted unless corroborated by another source.

| Alignment ID | Ionization Mode | Level 1 Identification | Average Rt(min) | Average Mz | BO-0/A | BO-BO/A | p.value | fdr adjusted p.value | Fold Change | Log2 fold change |
|---|---|---|---|---|---|---|---|---|---|---|
| 183 | Negative | w/o MS2:N-acetyl-D-mannosamine | 4.91 | 220.08 | 223.99 | 375.82 | 0.01 | 0.02 | 1.68 | 0.75 |
| 246 | Negative | Unknown | 9.05 | 252.02 | 405.65 | 687.86 | 0.02 | 0.05 | 1.70 | 0.76 |
| 254 | Negative | Unknown | 3.07 | 257.02 | 16.11 | 34.54 | 0.02 | 0.03 | 2.14 | 1.10 |
| 263 | Negative | Unknown | 2.94 | 261.00 | 328.54 | 623.99 | 0.01 | 0.03 | 1.90 | 0.93 |
| 276 | Negative | Unknown | 9.10 | 267.02 | 253.95 | 436.81 | 0.01 | 0.04 | 1.72 | 0.78 |
| 374 | Negative | Unknown | 8.92 | 311.07 | 63.11 | 140.61 | 0.00 | 0.01 | 2.23 | 1.16 |
| 398 | Negative | Unknown | 12.62 | 321.10 | 247.44 | 461.58 | 0.01 | 0.02 | 1.87 | 0.90 |
| 464 | Negative | Unknown | 10.12 | 351.13 | 1,980.91 | 3,178.27 | 0.01 | 0.04 | 1.60 | 0.68 |
| 473 | Negative | Unknown | 9.96 | 355.05 | 33.69 | 67.54 | 0.00 | 0.01 | 2.00 | 1.00 |
| 476 | Negative | Unknown | 5.17 | 355.09 | 74.91 | 126.66 | 0.01 | 0.02 | 1.69 | 0.76 |
| 519 | Negative | Unknown | 9.11 | 373.08 | 2,062.29 | 3,317.03 | 0.02 | 0.04 | 1.61 | 0.69 |
| 542 | Negative | Unknown | 9.95 | 384.99 | 29.36 | 48.48 | 0.01 | 0.02 | 1.65 | 0.72 |
| 545 | Negative | Unknown | 10.04 | 385.11 | 79.96 | 126.43 | 0.01 | 0.02 | 1.58 | 0.66 |
| 546 | Negative | Unknown | 10.49 | 385.11 | 433.40 | 664.15 | 0.01 | 0.04 | 1.53 | 0.62 |
| 552 | Negative | Unknown | 11.91 | 387.09 | 50.09 | 86.12 | 0.02 | 0.04 | 1.72 | 0.78 |
| 565 | Negative | Unknown | 2.93 | 390.96 | 107.06 | 203.19 | 0.01 | 0.03 | 1.90 | 0.92 |
| 592 | Negative | Unknown | 9.12 | 405.05 | 29.84 | 50.57 | 0.01 | 0.03 | 1.69 | 0.76 |
| 644 | Negative | Unknown | 9.88 | 431.12 | 11.79 | 18.82 | 0.02 | 0.03 | 1.60 | 0.68 |
| 645 | Negative | Unknown | 9.48 | 431.12 | 205.27 | 383.41 | 0.00 | 0.02 | 1.87 | 0.90 |
| 650 | Negative | Unknown | 9.47 | 433.14 | 758.22 | 1,204.67 | 0.01 | 0.02 | 1.59 | 0.67 |
| 667 | Negative | Unknown | 10.75 | 442.09 | 199.92 | 321.23 | 0.01 | 0.02 | 1.61 | 0.68 |
| 670 | Negative | Unknown | 10.75 | 444.11 | 82.42 | 132.00 | 0.01 | 0.03 | 1.60 | 0.68 |
| 682 | Negative | Unknown | 10.12 | 451.05 | 186.98 | 334.03 | 0.01 | 0.03 | 1.79 | 0.84 |
| 713 | Negative | Unknown | 9.35 | 461.13 | 10.57 | 16.11 | 0.03 | 0.05 | 1.52 | 0.61 |
| 716 | Negative | Unknown | 10.66 | 461.17 | 278.15 | 438.39 | 0.01 | 0.02 | 1.58 | 0.66 |
| 720 | Negative | Unknown | 9.62 | 463.14 | 149.40 | 269.25 | 0.00 | 0.01 | 1.80 | 0.85 |
| 739 | Negative | Unknown | 10.37 | 472.10 | 33.23 | 66.73 | 0.01 | 0.03 | 2.01 | 1.01 |
| 762 | Negative | Unknown | 10.50 | 485.04 | 44.24 | 75.60 | 0.01 | 0.03 | 1.71 | 0.77 |
| 788 | Negative | Unknown | 12.81 | 498.18 | 55.30 | 93.90 | 0.03 | 0.05 | 1.70 | 0.76 |
| 796 | Negative | Unknown | 5.07 | 503.16 | 1,575.30 | 3,175.05 | 0.02 | 0.04 | 2.02 | 1.01 |
| 797 | Negative | Unknown | 12.14 | 503.18 | 65.50 | 132.89 | 0.01 | 0.02 | 2.03 | 1.02 |
| 808 | Negative | Unknown | 12.69 | 507.21 | 1,056.05 | 1,828.34 | 0.01 | 0.02 | 1.73 | 0.79 |
| 814 | Negative | Unknown | 13.13 | 509.22 | 701.73 | 1,091.87 | 0.01 | 0.05 | 1.56 | 0.64 |
| 824 | Negative | Unknown | 4.76 | 515.13 | 63.18 | 156.72 | 0.00 | 0.01 | 2.48 | 1.31 |
| 856 | Negative | Unknown | 9.86 | 533.17 | 162.46 | 280.03 | 0.02 | 0.04 | 1.72 | 0.79 |
| 858 | Negative | Unknown | 4.97 | 533.17 | 148.61 | 388.62 | 0.01 | 0.05 | 2.62 | 1.39 |
| 862 | Negative | Unknown | 5.32 | 535.15 | 11.46 | 32.00 | 0.01 | 0.03 | 2.79 | 1.48 |
| 868 | Negative | Unknown | 4.84 | 537.17 | 95.41 | 194.50 | 0.03 | 0.04 | 2.04 | 1.03 |
| 895 | Negative | Unknown | 12.98 | 552.19 | 100.77 | 157.06 | 0.02 | 0.04 | 1.56 | 0.64 |
| 916 | Negative | Unknown | 9.26 | 573.15 | 115.71 | 214.08 | 0.00 | 0.00 | 1.85 | 0.89 |
| 918 | Negative | Unknown | 11.43 | 575.20 | 39.83 | 61.61 | 0.02 | 0.04 | 1.55 | 0.63 |
| 923 | Negative | Unknown | 9.64 | 583.20 | 111.18 | 190.77 | 0.00 | 0.02 | 1.72 | 0.78 |
| 927 | Negative | Unknown | 12.98 | 589.15 | 73.57 | 125.52 | 0.01 | 0.02 | 1.71 | 0.77 |
| 957 | Negative | Unknown | 14.71 | 619.42 | 68.44 | 143.10 | 0.01 | 0.04 | 2.09 | 1.06 |
| 980 | Negative | Unknown | 5.33 | 651.16 | 40.59 | 77.32 | 0.01 | 0.04 | 1.90 | 0.93 |

| | | | | | | | | | | |
|---|---|---|---|---|---|---|---|---|---|---|
| 993 | Negative | Unknown | 8.13 | 668.20 | 20.67 | 41.05 | 0.01 | 0.01 | 1.99 | 0.99 |
| 1012 | Negative | Unknown | 15.32 | 685.48 | 33.58 | 80.42 | 0.01 | 0.03 | 2.39 | 1.26 |
| 1016 | Negative | Unknown | 11.39 | 687.19 | 70.65 | 119.58 | 0.01 | 0.02 | 1.69 | 0.76 |
| 1024 | Negative | Unknown | 5.06 | 695.22 | 32.46 | 108.82 | 0.02 | 0.05 | 3.35 | 1.75 |
| 1064 | Negative | Unknown | 12.04 | 725.32 | 22.61 | 36.50 | 0.02 | 0.04 | 1.61 | 0.69 |
| 1099 | Negative | Unknown | 11.52 | 747.16 | 44.40 | 102.19 | 0.00 | 0.02 | 2.30 | 1.20 |
| 1104 | Negative | Unknown | 11.52 | 749.17 | 53.16 | 113.53 | 0.00 | 0.02 | 2.14 | 1.09 |
| 1141 | Negative | Unknown | 10.98 | 779.19 | 13.77 | 35.26 | 0.01 | 0.03 | 2.56 | 1.36 |
| 1150 | Negative | Unknown | 13.33 | 787.26 | 32.18 | 52.17 | 0.01 | 0.03 | 1.62 | 0.70 |
| 1163 | Negative | Unknown | 13.06 | 803.20 | 2.52 | 4.36 | 0.00 | 0.01 | 1.73 | 0.79 |
| 1169 | Negative | Unknown | 13.24 | 805.28 | 36.77 | 65.91 | 0.01 | 0.02 | 1.79 | 0.84 |
| 1190 | Negative | Unknown | 5.06 | 825.30 | 25.81 | 89.21 | 0.01 | 0.04 | 3.46 | 1.79 |
| 1209 | Negative | Unknown | 12.65 | 845.45 | 58.49 | 105.90 | 0.02 | 0.04 | 1.81 | 0.86 |
| 1305 | Negative | Unknown | 13.17 | 1,109.40 | 84.31 | 144.13 | 0.02 | 0.05 | 1.71 | 0.77 |
| 1328 | Negative | Unknown | 13.12 | 1,375.39 | 36.63 | 70.19 | 0.00 | 0.01 | 1.92 | 0.94 |
| 96 | Positive | Unknown | 3.69 | 85.03 | 242.86 | 408.31 | 0.01 | 0.03 | 1.68 | 0.75 |
| 97 | Positive | Unknown | 3.30 | 85.03 | 160.70 | 313.11 | 0.01 | 0.03 | 1.95 | 0.96 |
| 161 | Positive | Unknown | 3.57 | 104.11 | 153.19 | 354.16 | 0.02 | 0.04 | 2.31 | 1.21 |
| 213 | Positive | Unknown | 4.86 | 120.07 | 29.12 | 55.05 | 0.00 | 0.01 | 1.89 | 0.92 |
| 220 | Positive | Unknown | 6.70 | 123.04 | 175.80 | 294.66 | 0.02 | 0.05 | 1.68 | 0.75 |
| 230 | Positive | Unknown | 9.60 | 127.04 | 26.28 | 42.08 | 0.01 | 0.03 | 1.60 | 0.68 |
| 269 | Positive | Unknown | 6.72 | 136.08 | 378.75 | 660.71 | 0.01 | 0.03 | 1.74 | 0.80 |
| 301 | Positive | Unknown | 3.70 | 145.05 | 77.72 | 158.15 | 0.01 | 0.02 | 2.03 | 1.02 |
| 306 | Positive | Unknown | 5.16 | 146.09 | 62.85 | 137.74 | 0.01 | 0.03 | 2.19 | 1.13 |
| 311 | Positive | Glutamine | 3.22 | 147.08 | 508.91 | 1,151.84 | 0.00 | 0.01 | 2.26 | 1.18 |
| 326 | Positive | Unknown | 10.62 | 152.07 | 411.53 | 722.78 | 0.02 | 0.03 | 1.76 | 0.81 |
| 330 | Positive | Unknown | 5.43 | 154.02 | 64.09 | 104.72 | 0.01 | 0.04 | 1.63 | 0.71 |
| 335 | Positive | Unknown | 9.89 | 155.07 | 33.20 | 62.01 | 0.01 | 0.03 | 1.87 | 0.90 |
| 363 | Positive | Unknown | 10.72 | 162.05 | 30.26 | 74.14 | 0.02 | 0.04 | 2.45 | 1.29 |
| 451 | Positive | Unknown | 3.63 | 185.04 | 46.21 | 87.65 | 0.02 | 0.04 | 1.90 | 0.92 |
| 454 | Positive | Unknown | 5.01 | 186.08 | 210.18 | 436.29 | 0.00 | 0.01 | 2.08 | 1.05 |
| 465 | Positive | Unknown | 3.70 | 191.04 | 765.07 | 1,393.50 | 0.02 | 0.05 | 1.82 | 0.87 |
| 469 | Positive | Unknown | 4.74 | 192.09 | 84.00 | 143.15 | 0.01 | 0.01 | 1.70 | 0.77 |
| 492 | Positive | Unknown | 3.63 | 200.04 | 2,863.28 | 5,986.76 | 0.02 | 0.05 | 2.09 | 1.06 |
| 509 | Positive | Unknown | 4.96 | 204.09 | 346.96 | 686.27 | 0.01 | 0.03 | 1.98 | 0.98 |
| 559 | Positive | Unknown | 4.99 | 222.10 | 2,845.58 | 5,754.06 | 0.00 | 0.01 | 2.02 | 1.02 |
| 577 | Positive | Unknown | 6.05 | 226.07 | 19.15 | 44.09 | 0.00 | 0.01 | 2.30 | 1.20 |
| 578 | Positive | Unknown | 3.84 | 226.11 | 553.34 | 1,688.08 | 0.00 | 0.01 | 3.05 | 1.61 |
| 662 | Positive | Unknown | 4.85 | 249.11 | 102.03 | 197.05 | 0.02 | 0.05 | 1.93 | 0.95 |
| 673 | Positive | Unknown | 3.45 | 252.14 | 40.03 | 104.74 | 0.00 | 0.01 | 2.62 | 1.39 |
| 711 | Positive | Unknown | 4.88 | 262.09 | 58.74 | 128.87 | 0.01 | 0.03 | 2.19 | 1.13 |
| 723 | Positive | Unknown | 9.56 | 265.07 | 76.29 | 136.48 | 0.01 | 0.02 | 1.79 | 0.84 |
| 770 | Positive | Unknown | 5.04 | 277.14 | 48.53 | 114.33 | 0.01 | 0.04 | 2.36 | 1.24 |
| 784 | Positive | Unknown | 3.76 | 281.07 | 93.53 | 222.23 | 0.02 | 0.05 | 2.38 | 1.25 |
| 785 | Positive | Unknown | 4.88 | 281.07 | 29.48 | 56.69 | 0.02 | 0.04 | 1.92 | 0.94 |
| 800 | Positive | Unknown | 9.33 | 286.09 | 51.68 | 106.12 | 0.01 | 0.04 | 2.05 | 1.04 |
| 809 | Positive | Unknown | 9.16 | 288.09 | 640.51 | 1,256.53 | 0.01 | 0.05 | 1.96 | 0.97 |
| 813 | Positive | Unknown | 9.13 | 289.09 | 122.98 | 231.93 | 0.02 | 0.03 | 1.89 | 0.92 |
| 814 | Positive | Unknown | 4.38 | 289.09 | 272.59 | 598.87 | 0.01 | 0.01 | 2.20 | 1.14 |
| 836 | Positive | Unknown | 3.68 | 295.01 | 52.35 | 101.92 | 0.02 | 0.04 | 1.95 | 0.96 |
| 860 | Positive | Unknown | 11.44 | 301.56 | 29.29 | 68.44 | 0.01 | 0.04 | 2.34 | 1.22 |
| 874 | Positive | Unknown | 12.08 | 307.10 | 265.37 | 416.25 | 0.02 | 0.05 | 1.57 | 0.65 |
| 875 | Positive | Unknown | 12.96 | 307.10 | 202.74 | 334.27 | 0.01 | 0.04 | 1.65 | 0.72 |
| 888 | Positive | Unknown | 11.44 | 311.58 | 33.42 | 82.81 | 0.00 | 0.03 | 2.48 | 1.31 |
| 897 | Positive | Unknown | 10.16 | 314.12 | 46.67 | 80.02 | 0.02 | 0.04 | 1.71 | 0.78 |

| | | | | | | | | | | |
|---|---|---|---|---|---|---|---|---|---|---|
| 910 | Positive | Unknown | 15.38 | 319.19 | 92.28 | 276.54 | 0.01 | 0.02 | 3.00 | 1.58 |
| 923 | Positive | Unknown | 5.00 | 323.22 | 365.65 | 3,602.45 | 0.00 | 0.02 | 9.85 | 3.30 |
| 1025 | Positive | Unknown | 4.88 | 362.10 | 99.60 | 207.93 | 0.02 | 0.05 | 2.09 | 1.06 |
| 1062 | Positive | Unknown | 11.45 | 375.09 | 126.83 | 314.77 | 0.01 | 0.03 | 2.48 | 1.31 |
| 1065 | Positive | Unknown | 11.45 | 376.10 | 137.93 | 337.05 | 0.01 | 0.04 | 2.44 | 1.29 |
| 1072 | Positive | Unknown | 15.33 | 377.74 | 74.90 | 182.36 | 0.01 | 0.04 | 2.43 | 1.28 |
| 1102 | Positive | Unknown | 10.94 | 390.10 | 37.89 | 110.33 | 0.01 | 0.03 | 2.91 | 1.54 |
| 1120 | Positive | Unknown | 15.34 | 397.76 | 27.24 | 77.44 | 0.01 | 0.02 | 2.84 | 1.51 |
| 1123 | Positive | Unknown | 15.39 | 398.76 | 179.20 | 462.86 | 0.02 | 0.05 | 2.58 | 1.37 |
| 1139 | Positive | Unknown | 10.47 | 404.15 | 142.25 | 224.59 | 0.02 | 0.05 | 1.58 | 0.66 |
| 1148 | Positive | Unknown | 15.35 | 410.76 | 54.15 | 148.65 | 0.01 | 0.02 | 2.75 | 1.46 |
| 1170 | Positive | Unknown | 9.95 | 420.19 | 23.74 | 37.26 | 0.01 | 0.03 | 1.57 | 0.65 |
| 1173 | Positive | Unknown | 9.14 | 422.11 | 231.64 | 441.61 | 0.01 | 0.05 | 1.91 | 0.93 |
| 1186 | Positive | Unknown | 5.43 | 427.07 | 29.91 | 49.02 | 0.02 | 0.05 | 1.64 | 0.71 |
| 1202 | Positive | Unknown | 12.72 | 433.11 | 26.55 | 43.95 | 0.02 | 0.05 | 1.66 | 0.73 |
| 1203 | Positive | Unknown | 9.89 | 433.13 | 78.91 | 142.49 | 0.01 | 0.01 | 1.81 | 0.85 |
| 1205 | Positive | Unknown | 10.69 | 434.20 | 194.00 | 296.91 | 0.01 | 0.04 | 1.53 | 0.61 |
| 1206 | Positive | Unknown | 9.64 | 436.18 | 67.64 | 135.83 | 0.00 | 0.01 | 2.01 | 1.01 |
| 1226 | Positive | Unknown | 10.73 | 444.11 | 409.10 | 691.99 | 0.01 | 0.02 | 1.69 | 0.76 |
| 1235 | Positive | Unknown | 9.14 | 446.12 | 64.64 | 128.29 | 0.01 | 0.03 | 1.98 | 0.99 |
| 1293 | Positive | Unknown | 9.89 | 466.19 | 147.91 | 277.34 | 0.01 | 0.02 | 1.88 | 0.91 |
| 1314 | Positive | Unknown | 10.36 | 474.12 | 54.45 | 129.54 | 0.00 | 0.01 | 2.38 | 1.25 |
| 1322 | Positive | Unknown | 12.17 | 476.21 | 40.25 | 88.75 | 0.00 | 0.01 | 2.21 | 1.14 |
| 1327 | Positive | Unknown | 15.36 | 478.33 | 168.81 | 418.99 | 0.01 | 0.04 | 2.48 | 1.31 |
| 1337 | Positive | Unknown | 12.16 | 481.17 | 32.36 | 68.60 | 0.01 | 0.02 | 2.12 | 1.08 |
| 1350 | Positive | Unknown | 14.73 | 487.27 | 42.88 | 92.08 | 0.00 | 0.02 | 2.15 | 1.10 |
| 1365 | Positive | Unknown | 12.70 | 491.21 | 120.61 | 209.03 | 0.01 | 0.03 | 1.73 | 0.79 |
| 1390 | Positive | Unknown | 15.30 | 502.33 | 105.17 | 242.22 | 0.02 | 0.05 | 2.30 | 1.20 |
| 1455 | Positive | Unknown | 12.69 | 531.20 | 141.44 | 216.82 | 0.03 | 0.04 | 1.53 | 0.62 |
| 1762 | Positive | Unknown | 13.28 | 663.19 | 124.81 | 211.79 | 0.01 | 0.04 | 1.70 | 0.76 |
| 1773 | Positive | Unknown | 8.14 | 670.22 | 44.34 | 109.23 | 0.00 | 0.01 | 2.46 | 1.30 |
| 1787 | Positive | Unknown | 15.19 | 676.49 | 115.63 | 240.15 | 0.02 | 0.04 | 2.08 | 1.05 |
| 1788 | Positive | Unknown | 13.36 | 677.21 | 733.18 | 1,165.72 | 0.01 | 0.04 | 1.59 | 0.67 |
| 1815 | Positive | Unknown | 13.14 | 689.21 | 133.71 | 209.94 | 0.01 | 0.03 | 1.57 | 0.65 |
| 1869 | Positive | Unknown | 12.70 | 716.28 | 80.54 | 160.09 | 0.01 | 0.03 | 1.99 | 0.99 |
| 1894 | Positive | Unknown | 12.95 | 726.26 | 43.26 | 82.55 | 0.00 | 0.02 | 1.91 | 0.93 |
| 1920 | Positive | Unknown | 15.15 | 738.51 | 72.55 | 176.69 | 0.02 | 0.04 | 2.44 | 1.28 |
| 1951 | Positive | Unknown | 15.28 | 756.55 | 102.85 | 260.93 | 0.02 | 0.04 | 2.54 | 1.34 |
| 1976 | Positive | Unknown | 14.82 | 772.59 | 30.04 | 63.64 | 0.02 | 0.05 | 2.12 | 1.08 |
| 2021 | Positive | Unknown | 14.73 | 814.64 | 25.47 | 52.18 | 0.01 | 0.02 | 2.05 | 1.03 |
| 2087 | Positive | Unknown | 14.73 | 934.65 | 34.07 | 72.07 | 0.01 | 0.03 | 2.12 | 1.08 |
| 2097 | Positive | Unknown | 14.77 | 955.58 | 19.99 | 50.19 | 0.02 | 0.05 | 2.51 | 1.33 |

**Table S6:** Compounds upregulated by BO-P/A when compared to BO-0/A for the second extract of the P experimental set. For identification level 1, the annotation" w/o MS2" indicates there was not confident spectral match and that the results can not be fully trusted unless corroborated by another source.

| Alignment ID | Ionization Mode | Level 1 Identification | Level 2a Identification | Level 2b Identification | Average Rt(min) | Average Mz | BO-0/A | BO-P/A | p.value | fdr adjusted p.value | Fold Change | Alignment ID |
|---|---|---|---|---|---|---|---|---|---|---|---|---|
| 2 | Negative | Unknown | NA | NA | 11.89 | 57.03 | 58.06 | 121.54 | 0.01 | 0.02 | 2.09 | 2 |
| 4 | Negative | Unknown | NA | NA | 11.89 | 59.01 | 160.23 | 310.51 | 0.01 | 0.04 | 1.94 | 4 |
| 69 | Negative | Unknown | NA | NA | 11.89 | 99.04 | 146.18 | 352.85 | 0.00 | 0.01 | 2.41 | 69 |
| 70 | Negative | Unknown | NA | NA | 3.74 | 99.04 | 5.33 | 33.93 | 0.03 | 0.05 | 6.36 | 70 |
| 129 | Negative | Unknown | NA | NA | 12.47 | 121.00 | 100.04 | 176.63 | 0.02 | 0.03 | 1.77 | 129 |
| 136 | Negative | Unknown | NA | NA | 11.89 | 125.02 | 75.38 | 154.32 | 0.01 | 0.03 | 2.05 | 136 |
| 229 | Negative | Unknown | NA | NA | 3.98 | 165.04 | 24.63 | 43.47 | 0.03 | 0.05 | 1.76 | 229 |
| 335 | Negative | Unknown | NA | NA | 4.70 | 195.05 | 102.23 | 220.16 | 0.03 | 0.04 | 2.15 | 335 |
| 392 | Negative | Unknown | NA | NA | 13.52 | 215.03 | 107.13 | 193.72 | 0.02 | 0.04 | 1.81 | 392 |
| 414 | Negative | w/o MS2:N-acetyl-D-mannosamine | NA | NA | 5.09 | 220.08 | 395.67 | 1088.52 | 0.00 | 0.01 | 2.75 | 414 |
| 425 | Negative | Unknown | NA | NA | 2.90 | 224.09 | 45.35 | 146.89 | 0.00 | 0.01 | 3.24 | 425 |
| 508 | Negative | Unknown | NA | NA | 4.99 | 253.09 | 72.82 | 211.17 | 0.00 | 0.01 | 2.90 | 508 |
| 528 | Negative | Unknown | NA | NA | 4.53 | 260.02 | 75.19 | 158.25 | 0.03 | 0.05 | 2.10 | 528 |
| 682 | Negative | Unknown | NA | NA | 5.17 | 313.11 | 33.77 | 68.59 | 0.00 | 0.01 | 2.03 | 682 |
| 704 | Negative | Unknown | NA | NA | 12.47 | 321.10 | 514.21 | 985.10 | 0.02 | 0.03 | 1.92 | 704 |
| 741 | Negative | Unknown | NA | NA | 4.68 | 334.13 | 11.45 | 24.72 | 0.03 | 0.04 | 2.16 | 741 |
| 744 | Negative | Unknown | NA | NA | 10.87 | 335.14 | 105.81 | 243.47 | 0.02 | 0.04 | 2.30 | 744 |
| 843 | Negative | Unknown | NA | NA | 9.03 | 373.08 | 3535.57 | 6537.33 | 0.01 | 0.02 | 1.85 | 843 |
| 888 | Negative | Unknown | NA | NA | 11.68 | 387.09 | 51.25 | 109.91 | 0.02 | 0.04 | 2.14 | 888 |
| 889 | Negative | Unknown | NA | NA | 9.15 | 387.10 | 588.64 | 1011.53 | 0.02 | 0.05 | 1.72 | 889 |
| 917 | Negative | Unknown | NA | NA | 11.54 | 393.18 | 78.41 | 151.18 | 0.02 | 0.03 | 1.93 | 917 |
| 935 | Negative | Unknown | NA | PubChem:(46222441) | 8.49 | 403.09 | 144.79 | 307.92 | 0.02 | 0.04 | 2.13 | 935 |
| 960 | Negative | Unknown | NA | COCONUT:(CNP0159349 CNP0218833);Natural Products:(UNPD202200);PubChem:(38363079 38363084 45360328 44715457 125416071 125416072 125416073 125416074);SuperNatural:(SN00032625 SN00030299 SN00032624 SN00030300 SN00032623 SN00032622);ZINC bio:(ZINC31169353 ZINC31169357 ZINC35454681 ZINC35454685 ZINC35454688 ZINC35454690);Training Set | 11.89 | 413.15 | 4009.66 | 9883.57 | 0.01 | 0.01 | 2.46 | 960 |
| 961 | Negative | Unknown | NA | COCONUT:(CNP0159349 CNP0218833);Natural Products:(UNPD202200);PubChem:(38363079 38363084 45360328 44715457 125416071 125416072 125416073 125416074);SuperNatural:(SN00032625 SN00030299 SN00032624 SN00030300 SN00032623 SN00032622);ZINC bio:(ZINC31169353 ZINC31169357 ZINC35454681 ZINC35454685 ZINC35454688 ZINC35454690);Training Set | 11.06 | 413.15 | 60.28 | 143.89 | 0.01 | 0.01 | 2.39 | 961 |
| 962 | Negative | Unknown | NA | NA | 10.55 | 413.15 | 35.24 | 68.79 | 0.03 | 0.05 | 1.95 | 962 |
| 966 | Negative | Unknown | NA | NA | 5.20 | 415.14 | 306.35 | 650.84 | 0.01 | 0.04 | 2.12 | 966 |
| 972 | Negative | Unknown | NA | NA | 9.08 | 417.11 | 159.29 | 269.20 | 0.01 | 0.03 | 1.69 | 972 |
| 988 | Negative | Unknown | NA | NA | 9.15 | 421.08 | 84.13 | 166.39 | 0.01 | 0.02 | 1.98 | 988 |
| 991 | Negative | Unknown | NA | NA | 10.05 | 421.16 | 50.93 | 89.64 | 0.02 | 0.04 | 1.76 | 991 |
| 1034 | Negative | Unknown | NA | NA | 12.71 | 435.19 | 937.41 | 1993.15 | 0.01 | 0.02 | 2.13 | 1034 |
| 1061 | Negative | Unknown | NA | NA | 9.40 | 447.11 | 17.13 | 39.42 | 0.01 | 0.02 | 2.30 | 1061 |
| 1070 | Negative | Unknown | NA | NA | 12.31 | 449.20 | 106.14 | 171.73 | 0.02 | 0.05 | 1.62 | 1070 |
| 1089 | Negative | Unknown | NA | NA | 9.47 | 455.12 | 547.17 | 1022.45 | 0.01 | 0.04 | 1.87 | 1089 |
| 1092 | Negative | Unknown | NA | NA | 12.86 | 456.15 | 83.74 | 139.17 | 0.01 | 0.04 | 1.66 | 1092 |

| | | | | | | | | | | | | |
|---|---|---|---|---|---|---|---|---|---|---|---|---|
| 1123 | Negative | Unknown | NA | NA | 9.11 | 467.05 | 30.30 | 64.10 | 0.01 | 0.03 | 2.12 | 1123 |
| 1125 | Negative | Unknown | NA | NA | 12.77 | 467.21 | 416.56 | 812.94 | 0.01 | 0.01 | 1.95 | 1125 |
| 1149 | Negative | Unknown | NA | NA | 10.87 | 479.05 | 27.30 | 73.18 | 0.02 | 0.05 | 2.68 | 1149 |
| 1214 | Negative | Unknown | NA | NA | 12.58 | 507.21 | 1545.30 | 2691.83 | 0.02 | 0.03 | 1.74 | 1214 |
| 1230 | Negative | Unknown | NA | NA | 4.96 | 515.12 | 25.10 | 71.12 | 0.02 | 0.05 | 2.83 | 1230 |
| 1242 | Negative | Unknown | NA | NA | 13.28 | 519.24 | 52.95 | 103.77 | 0.01 | 0.04 | 1.96 | 1242 |
| 1258 | Negative | Unknown | NA | NA | 10.36 | 533.16 | 46.32 | 77.90 | 0.02 | 0.04 | 1.68 | 1258 |
| 1259 | Negative | Unknown | NA | NA | 9.76 | 533.17 | 261.98 | 497.17 | 0.01 | 0.03 | 1.90 | 1259 |
| 1311 | Negative | Unknown | NA | NA | 5.10 | 562.20 | 41.51 | 131.41 | 0.01 | 0.03 | 3.17 | 1311 |
| 1329 | Negative | Unknown | NA | NA | 11.28 | 575.20 | 58.16 | 138.13 | 0.01 | 0.03 | 2.38 | 1329 |
| 1339 | Negative | Unknown | NA | NA | 10.63 | 589.18 | 43.81 | 93.32 | 0.01 | 0.03 | 2.13 | 1339 |
| 1410 | Negative | Unknown | NA | NA | 14.72 | 653.51 | 15.89 | 31.36 | 0.02 | 0.05 | 1.97 | 1410 |
| 1415 | Negative | Unknown | NA | NA | 12.63 | 661.18 | 54.26 | 115.99 | 0.00 | 0.01 | 2.14 | 1415 |
| 1428 | Negative | Unknown | NA | NA | 5.66 | 665.21 | 506.86 | 1049.35 | 0.01 | 0.02 | 2.07 | 1428 |
| 1467 | Negative | Unknown | NA | NA | 13.05 | 687.19 | 1336.43 | 2084.86 | 0.02 | 0.04 | 1.56 | 1467 |
| 1486 | Negative | Unknown | NA | NA | 12.60 | 697.24 | 108.02 | 300.95 | 0.00 | 0.01 | 2.79 | 1486 |
| 1507 | Negative | Unknown | NA | NA | 11.88 | 711.30 | 25.52 | 66.18 | 0.01 | 0.01 | 2.59 | 1507 |
| 1547 | Negative | Unknown | NA | NA | 13.27 | 738.19 | 87.08 | 135.25 | 0.03 | 0.05 | 1.55 | 1547 |
| 1571 | Negative | Unknown | NA | NA | 11.71 | 749.17 | 51.35 | 119.39 | 0.02 | 0.03 | 2.32 | 1571 |
| 1610 | Negative | Unknown | NA | NA | 5.09 | 781.20 | 60.68 | 194.28 | 0.00 | 0.05 | 3.20 | 1610 |
| 1635 | Negative | Unknown | NA | NA | 11.08 | 808.21 | 67.59 | 158.40 | 0.02 | 0.03 | 2.34 | 1635 |
| 1636 | Negative | Unknown | NA | NA | 12.57 | 808.21 | 44.65 | 98.70 | 0.01 | 0.03 | 2.21 | 1636 |
| 1691 | Negative | Unknown | NA | NA | 12.58 | 845.45 | 54.65 | 97.66 | 0.02 | 0.03 | 1.79 | 1691 |
| 1697 | Negative | Unknown | NA | NA | 12.64 | 857.25 | 83.52 | 133.85 | 0.02 | 0.05 | 1.60 | 1697 |
| 1730 | Negative | Unknown | NA | NA | 13.33 | 943.29 | 226.06 | 450.79 | 0.00 | 0.02 | 1.99 | 1730 |
| 1808 | Negative | Unknown | NA | NA | 13.25 | 1105.34 | 58.88 | 94.50 | 0.01 | 0.04 | 1.61 | 1808 |
| 1809 | Negative | Unknown | NA | NA | 13.09 | 1109.41 | 64.05 | 119.66 | 0.01 | 0.02 | 1.87 | 1809 |
| 1825 | Negative | Unknown | NA | NA | 13.05 | 1375.39 | 37.25 | 73.63 | 0.01 | 0.01 | 1.98 | 1825 |
| 35 | Positive | Unknown | NA | NA | 2.97 | 66.99 | 29.23 | 51.07 | 0.00 | 0.02 | 1.75 | 35 |
| 216 | Positive | Unknown | NA | NA | 4.88 | 120.07 | 32.56 | 72.71 | 0.01 | 0.04 | 2.23 | 216 |
| 274 | Positive | Unknown | NA | 2-Aminobenzoic acid | 4.97 | 138.05 | 650.74 | 1509.55 | 0.01 | 0.02 | 2.32 | 274 |
| 386 | Positive | Unknown | NA | NA | 4.99 | 176.09 | 120.45 | 203.22 | 0.00 | 0.00 | 1.69 | 386 |
| 387 | Positive | Unknown | NA | NA | 3.72 | 176.09 | 102.02 | 199.12 | 0.01 | 0.04 | 1.95 | 387 |
| 422 | Positive | Unknown | NA | NA | 3.54 | 185.04 | 26.28 | 62.02 | 0.01 | 0.03 | 2.36 | 422 |
| 445 | Positive | Unknown | NA | NA | 4.76 | 192.09 | 77.68 | 157.42 | 0.02 | 0.04 | 2.03 | 445 |
| 490 | Positive | Unknown | NA | NA | 4.94 | 204.09 | 653.25 | 1400.84 | 0.00 | 0.01 | 2.14 | 490 |
| 538 | Positive | Unknown | NA | N-Acetylgalactosamine | 4.99 | 222.10 | 3804.84 | 7088.53 | 0.00 | 0.00 | 1.86 | 538 |
| 613 | Positive | Unknown | NA | NA | 6.03 | 244.08 | 66.88 | 148.61 | 0.00 | 0.01 | 2.22 | 613 |
| 625 | Positive | Unknown | NA | NA | 5.07 | 246.09 | 155.20 | 409.90 | 0.00 | 0.02 | 2.64 | 625 |
| 639 | Positive | Unknown | NA | NA | 4.88 | 249.11 | 127.11 | 288.35 | 0.00 | 0.01 | 2.27 | 639 |
| 713 | Positive | Unknown | NA | NA | 10.93 | 270.13 | 572.20 | 993.35 | 0.00 | 0.01 | 1.74 | 713 |
| 718 | Positive | Unknown | NA | NA | 5.14 | 272.07 | 74.24 | 185.26 | 0.01 | 0.02 | 2.50 | 718 |
| 720 | Positive | Unknown | NA | NA | 5.33 | 272.07 | 74.96 | 187.19 | 0.00 | 0.01 | 2.50 | 720 |
| 799 | Positive | Unknown | NA | NA | 4.79 | 293.14 | 36.93 | 89.92 | 0.02 | 0.04 | 2.44 | 799 |
| 807 | Positive | Unknown | NA | NA | 9.40 | 295.10 | 62.11 | 114.03 | 0.02 | 0.04 | 1.84 | 807 |
| 973 | Positive | Unknown | NA | NA | 5.23 | 353.09 | 30.92 | 84.58 | 0.01 | 0.03 | 2.74 | 973 |
| 974 | Positive | Unknown | NA | NA | 5.50 | 353.09 | 35.21 | 102.44 | 0.01 | 0.02 | 2.91 | 974 |
| 993 | Positive | Unknown | NA | NA | 4.92 | 362.10 | 140.06 | 310.92 | 0.01 | 0.04 | 2.22 | 993 |
| 1000 | Positive | Dihexose (Maltose/Sucrose) | NA | NA | 4.92 | 365.11 | 123.01 | 243.34 | 0.01 | 0.04 | 1.98 | 1000 |
| 1045 | Positive | Unknown | NA | NA | 4.91 | 381.08 | 37.34 | 87.60 | 0.00 | 0.04 | 2.35 | 1045 |
| 1430 | Positive | Unknown | Melezitose | NA | 5.14 | 522.20 | 20.53 | 83.62 | 0.00 | 0.00 | 4.07 | 1430 |
| 1436 | Positive | Unknown | NA | NA | 5.15 | 524.15 | 5.41 | 89.06 | 0.01 | 0.02 | 16.45 | 1436 |
| 1445 | Positive | Unknown | NA | NA | 5.17 | 527.16 | 20.64 | 70.20 | 0.00 | 0.00 | 3.40 | 1445 |
| 1461 | Positive | Unknown | NA | NA | 4.92 | 533.16 | 53.45 | 239.80 | 0.01 | 0.03 | 4.49 | 1461 |

| | | | | | | | | | | | | |
|---|---|---|---|---|---|---|---|---|---|---|---|---|
| 1879 | Positive | Unknown | NA | NA | 4.92 | 704.21 | 10.77 | 91.55 | 0.00 | 0.03 | 8.50 | 1879 |
| 2049 | Positive | Unknown | NA | NA | 14.02 | 786.44 | 104.64 | 231.98 | 0.02 | 0.04 | 2.22 | 2049 |
| 2138 | Positive | Unknown | NA | NA | 14.38 | 916.51 | 45.40 | 80.80 | 0.02 | 0.04 | 1.78 | 2138 |
| 2148 | Positive | Unknown | NA | NA | 14.28 | 930.48 | 61.44 | 159.72 | 0.01 | 0.01 | 2.60 | 2148 |

**Table S7:** Compounds upregulated by BO-G/A when compared to BO-0/A for the second extract of the G experimental set**.** For identification level 1, the annotation" w/o MS2" indicates there was not confident spectral match and that the results can not be fully trusted unless corroborated by another source.

| Alignment ID | Ionization Mode | Level 1 Identification | Average Rt(min) | Average Mz | BO-0/A | BO-G/A | p.value | fdr adjusted p.value | Fold Change | Log2 fold change |
|---|---|---|---|---|---|---|---|---|---|---|
| 139 | Negative | Unknown | 4.81 | 195.05 | 411.33 | 653.32 | 0.02 | 0.04 | 1.59 | 0.67 |
| 173 | Negative | Unknown | 13.69 | 215.03 | 43.4 | 76.9 | 0.01 | 0.02 | 1.77 | 0.83 |
| 183 | Negative | w/o MS2:N-acetyl-D-mannosamine | 4.91 | 220.08 | 223.99 | 476.15 | 0.01 | 0.02 | 2.13 | 1.09 |
| 233 | Negative | Unknown | 12.39 | 245.09 | 37.17 | 63.77 | 0.02 | 0.05 | 1.72 | 0.78 |
| 246 | Negative | Unknown | 9.05 | 252.02 | 405.65 | 706.89 | 0.02 | 0.05 | 1.74 | 0.8 |
| 254 | Negative | Unknown | 3.07 | 257.02 | 16.11 | 30.91 | 0.02 | 0.03 | 1.92 | 0.94 |
| 276 | Negative | Unknown | 9.1 | 267.02 | 253.95 | 453.71 | 0.01 | 0.03 | 1.79 | 0.84 |
| 300 | Negative | Unknown | 13.39 | 279.01 | 186.7 | 314.47 | 0.01 | 0.02 | 1.68 | 0.75 |
| 312 | Negative | Unknown | 9.95 | 285.06 | 187.08 | 296.33 | 0.02 | 0.04 | 1.58 | 0.66 |
| 398 | Negative | Unknown | 12.62 | 321.1 | 247.44 | 495.59 | 0.01 | 0.02 | 2 | 1 |
| 431 | Negative | Unknown | 11.02 | 335.13 | 56.15 | 118.29 | 0 | 0.01 | 2.11 | 1.08 |
| 436 | Negative | Unknown | 9.44 | 337.11 | 99.28 | 167.43 | 0.01 | 0.03 | 1.69 | 0.75 |
| 449 | Negative | Unknown | 9.53 | 343.1 | 150.07 | 288.49 | 0 | 0.02 | 1.92 | 0.94 |
| 451 | Negative | Unknown | 9.15 | 344.1 | 92.21 | 148.84 | 0.02 | 0.05 | 1.61 | 0.69 |
| 473 | Negative | Unknown | 9.96 | 355.05 | 33.69 | 62.38 | 0.01 | 0.03 | 1.85 | 0.89 |
| 476 | Negative | Unknown | 5.17 | 355.09 | 74.91 | 138.6 | 0.01 | 0.02 | 1.85 | 0.89 |
| 519 | Negative | Unknown | 9.11 | 373.08 | 2,062.29 | 3,302.24 | 0.02 | 0.05 | 1.6 | 0.68 |
| 520 | Negative | Unknown | 8.63 | 373.08 | 63.79 | 109.82 | 0.02 | 0.05 | 1.72 | 0.78 |
| 552 | Negative | Unknown | 11.91 | 387.09 | 50.09 | 113.78 | 0 | 0 | 2.27 | 1.18 |
| 592 | Negative | Unknown | 9.12 | 405.05 | 29.84 | 50.93 | 0.01 | 0.02 | 1.71 | 0.77 |
| 603 | Negative | Unknown | 11.18 | 413.15 | 49.05 | 99.12 | 0 | 0.01 | 2.02 | 1.01 |
| 604 | Negative | Unknown | 12.08 | 413.15 | 2,788.35 | 5,095.02 | 0.01 | 0.03 | 1.83 | 0.87 |
| 629 | Negative | Unknown | 9.47 | 423.11 | 64.82 | 117.97 | 0.01 | 0.02 | 1.82 | 0.86 |
| 633 | Negative | Unknown | 12.91 | 427.16 | 244.22 | 400.56 | 0.01 | 0.02 | 1.64 | 0.71 |
| 645 | Negative | Unknown | 9.48 | 431.12 | 205.27 | 446.97 | 0 | 0.02 | 2.18 | 1.12 |
| 650 | Negative | Unknown | 9.47 | 433.14 | 758.22 | 1,573.34 | 0 | 0 | 2.08 | 1.05 |
| 659 | Negative | Unknown | 9.06 | 439.05 | 17.44 | 27.19 | 0.02 | 0.04 | 1.56 | 0.64 |
| 675 | Negative | Unknown | 9.94 | 447.15 | 54.82 | 89.92 | 0.02 | 0.04 | 1.64 | 0.71 |
| 680 | Negative | Unknown | 9.46 | 450.13 | 91.61 | 201.53 | 0 | 0.01 | 2.2 | 1.14 |
| 686 | Negative | Unknown | 13.02 | 451.22 | 127.53 | 253.89 | 0.02 | 0.04 | 1.99 | 0.99 |
| 691 | Negative | Unknown | 11.12 | 453.2 | 33.92 | 68.82 | 0.01 | 0.04 | 2.03 | 1.02 |
| 713 | Negative | Unknown | 9.35 | 461.13 | 10.57 | 18.5 | 0.02 | 0.04 | 1.75 | 0.81 |
| 714 | Negative | Unknown | 9.67 | 461.13 | 27.73 | 53.7 | 0.03 | 0.05 | 1.94 | 0.95 |
| 718 | Negative | Unknown | 10.56 | 462.19 | 1.25 | 2.83 | 0.02 | 0.04 | 2.28 | 1.19 |
| 720 | Negative | Unknown | 9.62 | 463.14 | 149.4 | 244.64 | 0.02 | 0.05 | 1.64 | 0.71 |
| 721 | Negative | Unknown | 9.08 | 463.15 | 163.38 | 299.65 | 0.01 | 0.03 | 1.83 | 0.88 |
| 739 | Negative | Unknown | 10.37 | 472.1 | 33.23 | 58.38 | 0.02 | 0.03 | 1.76 | 0.81 |
| 747 | Negative | Unknown | 12.08 | 476.14 | 318.64 | 638.46 | 0.01 | 0.02 | 2 | 1 |
| 788 | Negative | Unknown | 12.81 | 498.18 | 55.3 | 100.33 | 0.02 | 0.04 | 1.81 | 0.86 |
| 799 | Negative | Unknown | 9.05 | 505.05 | 64.95 | 111.65 | 0.02 | 0.03 | 1.72 | 0.78 |
| 800 | Negative | Unknown | 9.47 | 505.07 | 59.66 | 122.22 | 0.01 | 0.02 | 2.05 | 1.03 |
| 801 | Negative | Unknown | 9.71 | 505.16 | 9.27 | 18.39 | 0.02 | 0.03 | 1.98 | 0.99 |
| 802 | Negative | Unknown | 9.47 | 505.16 | 98.74 | 184.89 | 0 | 0.01 | 1.87 | 0.9 |
| 808 | Negative | Unknown | 12.69 | 507.21 | 1,056.05 | 1,767.10 | 0.02 | 0.04 | 1.67 | 0.74 |
| 824 | Negative | Unknown | 4.76 | 515.13 | 63.18 | 132.11 | 0.02 | 0.04 | 2.09 | 1.06 |

| | | | | | | | | | | |
|---|---|---|---|---|---|---|---|---|---|---|
| 836 | Negative | Unknown | 10.13 | 521.12 | 10.96 | 27.67 | 0.02 | 0.05 | 2.52 | 1.34 |
| 846 | Negative | Unknown | 13.71 | 525.19 | 17.57 | 31.8 | 0.02 | 0.05 | 1.81 | 0.86 |
| 856 | Negative | Unknown | 9.86 | 533.17 | 162.46 | 280.06 | 0.02 | 0.05 | 1.72 | 0.79 |
| 857 | Negative | Unknown | 12.15 | 533.17 | 18.47 | 31.93 | 0.02 | 0.04 | 1.73 | 0.79 |
| 860 | Negative | Unknown | 9.11 | 535.06 | 47.22 | 87.01 | 0.01 | 0.04 | 1.84 | 0.88 |
| 862 | Negative | Unknown | 5.32 | 535.15 | 11.46 | 37.74 | 0.01 | 0.03 | 3.29 | 1.72 |
| 868 | Negative | Unknown | 4.84 | 537.17 | 95.41 | 230.42 | 0.01 | 0.02 | 2.42 | 1.27 |
| 905 | Negative | Unknown | 4.89 | 562.2 | 38.27 | 100.17 | 0.02 | 0.04 | 2.62 | 1.39 |
| 916 | Negative | Unknown | 9.26 | 573.15 | 115.71 | 246.56 | 0 | 0 | 2.13 | 1.09 |
| 918 | Negative | Unknown | 11.43 | 575.2 | 39.83 | 73.26 | 0.01 | 0.02 | 1.84 | 0.88 |
| 993 | Negative | Unknown | 8.13 | 668.2 | 20.67 | 47.15 | 0.01 | 0.01 | 2.28 | 1.19 |
| 1016 | Negative | Unknown | 11.39 | 687.19 | 70.65 | 121.72 | 0.01 | 0.02 | 1.72 | 0.78 |
| 1026 | Negative | Unknown | 12.71 | 697.23 | 58.61 | 116.66 | 0.01 | 0.04 | 1.99 | 0.99 |
| 1064 | Negative | Unknown | 12.04 | 725.32 | 22.61 | 42.11 | 0.01 | 0.03 | 1.86 | 0.9 |
| 1068 | Negative | Unknown | 13 | 727.21 | 54.05 | 109.22 | 0 | 0.03 | 2.02 | 1.02 |
| 1072 | Negative | Unknown | 13.03 | 729.2 | 5.33 | 9.27 | 0.02 | 0.04 | 1.74 | 0.8 |
| 1163 | Negative | Unknown | 13.06 | 803.2 | 2.52 | 4.19 | 0.01 | 0.02 | 1.66 | 0.73 |
| 1193 | Negative | Unknown | 12.07 | 827.3 | 60.25 | 238.15 | 0.01 | 0.02 | 3.95 | 1.98 |
| 1211 | Negative | Unknown | 11.33 | 849.09 | 1.22 | 1.94 | 0.02 | 0.04 | 1.59 | 0.67 |
| 1243 | Negative | Unknown | 13.41 | 943.29 | 222.65 | 367.3 | 0.02 | 0.05 | 1.65 | 0.72 |
| 1328 | Negative | Unknown | 13.12 | 1,375.39 | 36.63 | 68.56 | 0.01 | 0.03 | 1.87 | 0.9 |
| 161 | Positive | Unknown | 3.57 | 104.11 | 153.19 | 337.9 | 0.01 | 0.02 | 2.21 | 1.14 |
| 233 | Positive | Unknown | 12.09 | 127.04 | 80.4 | 143.23 | 0.01 | 0.03 | 1.78 | 0.83 |
| 335 | Positive | Unknown | 9.89 | 155.07 | 33.2 | 59.39 | 0.01 | 0.03 | 1.79 | 0.84 |
| 369 | Positive | Unknown | 12.09 | 163.06 | 252.5 | 433.76 | 0.02 | 0.04 | 1.72 | 0.78 |
| 383 | Positive | Unknown | 9.72 | 167.07 | 9.12 | 15.19 | 0.01 | 0.03 | 1.67 | 0.74 |
| 389 | Positive | Unknown | 9.11 | 170.08 | 24.9 | 46.13 | 0.02 | 0.04 | 1.85 | 0.89 |
| 469 | Positive | Unknown | 4.74 | 192.09 | 84 | 182.51 | 0 | 0.01 | 2.17 | 1.12 |
| 509 | Positive | Unknown | 4.96 | 204.09 | 346.96 | 713.02 | 0.02 | 0.04 | 2.06 | 1.04 |
| 559 | Positive | Unknown | 4.99 | 222.1 | 2,845.58 | 6,688.03 | 0.01 | 0.02 | 2.35 | 1.23 |
| 577 | Positive | Unknown | 6.05 | 226.07 | 19.15 | 51.08 | 0 | 0 | 2.67 | 1.42 |
| 644 | Positive | Unknown | 6.05 | 244.08 | 62.03 | 136.9 | 0 | 0.01 | 2.21 | 1.14 |
| 673 | Positive | Unknown | 3.45 | 252.14 | 40.03 | 77.36 | 0.02 | 0.04 | 1.93 | 0.95 |
| 840 | Positive | Unknown | 9.47 | 295.1 | 77.02 | 155.01 | 0.01 | 0.03 | 2.01 | 1.01 |
| 874 | Positive | Unknown | 12.08 | 307.1 | 265.37 | 466.97 | 0.01 | 0.03 | 1.76 | 0.82 |
| 1110 | Positive | Unknown | 9.42 | 394.13 | 166.65 | 361.76 | 0.01 | 0.04 | 2.17 | 1.12 |
| 1119 | Positive | Unknown | 12.08 | 397.15 | 256.67 | 453.72 | 0.01 | 0.04 | 1.77 | 0.82 |
| 1141 | Positive | Unknown | 9.46 | 406.17 | 246.51 | 440.51 | 0.01 | 0.03 | 1.79 | 0.84 |
| 1197 | Positive | Unknown | 12.08 | 432.19 | 303.7 | 537.6 | 0.01 | 0.03 | 1.77 | 0.82 |
| 1206 | Positive | Unknown | 9.64 | 436.18 | 67.64 | 113.19 | 0.02 | 0.04 | 1.67 | 0.74 |
| 1347 | Positive | Unknown | 12.86 | 486.25 | 62.03 | 133.72 | 0.01 | 0.02 | 2.16 | 1.11 |
| 1365 | Positive | Unknown | 12.7 | 491.21 | 120.61 | 212.06 | 0.01 | 0.03 | 1.76 | 0.81 |
| 1773 | Positive | Unknown | 8.14 | 670.22 | 44.34 | 102.92 | 0.01 | 0.02 | 2.32 | 1.21 |
| 1869 | Positive | Unknown | 12.7 | 716.28 | 80.54 | 156.11 | 0.01 | 0.03 | 1.94 | 0.95 |

**Table S8:** Number of compounds identified to each confidence level for redroot pigweed second extract samples

| Ionization Mode | Annotation | Number Of Compounds |
|---|---|---|
| Negative | Total Compounds | 952 |
| Negative | Total Fragmented | 846 |
| Negative | Confidence Level 3 | 566 |
| Negative | Confidence Level 2 | 56 |
| Negative | Confidence Level 2b | 37 |
| Negative | Confidence Level 2a | 28 |
| Negative | Confidence Level 1 | 15 |
| Positive | Total Compounds | 597 |
| Positive | Total Fragmented | 514 |
| Positive | Confidence Level 3 | 292 |
| Positive | Confidence Level 2 | 45 |
| Positive | Confidence Level 2b | 34 |
| Positive | Confidence Level 2a | 22 |
| Positive | Confidence Level 1 | 8 |

**Table S9:** Number of compounds identified to each confidence level for black grass second extract samples

| Ionization Mode | Annotation | Number Of Compounds |
|---|---|---|
| Negative | Total Compounds | 674 |
| Negative | Total Fragmented | 581 |
| Negative | Confidence Level 4 | 237 |
| Negative | Confidence Level 3 | 438 |
| Negative | Confidence Level 2 | 30 |
| Negative | Confidence Level 2b | 25 |
| Negative | Confidence Level 2a | 13 |
| Negative | Confidence Level 1 | 14 |
| Positive | Total Compounds | 620 |
| Positive | Total Fragmented | 521 |
| Positive | Confidence Level 4 | 271 |
| Positive | Confidence Level 3 | 297 |
| Positive | Confidence Level 2 | 45 |
| Positive | Confidence Level 2b | 37 |
| Positive | Confidence Level 2a | 17 |
| Positive | Confidence Level 1 | 7 |

**Table S10:** Compounds identified within the redroot pigweed experimental set second extract samples.

| Ionization Mode | Alignment ID | Average Rt(min) | Average Mz | Level 1 Identification | Level 2a Identification | Level 2b Identification |
|---|---|---|---|---|---|---|
| Negative | 0 | 7.501 | 57.034 | Unknown | NA | Propylene glycol |
| Negative | 17 | 5.499 | 71.01338 | Unknown | NA | L-Lactic acid |
| Negative | 124 | 3.011 | 117.019 | Succinic acid | SUCCINIC ACID | NA |
| Negative | 141 | 3.164 | 128.0351 | Pyroglutamic acid | NA | Pyroglutamic acid |
| Negative | 146 | 7.474 | 129.0189 | Unknown | Mesaconic acid; LC-ESI-QTOF; MS2; CE | NA |
| Negative | 156 | 2.913 | 131.046 | Asparagine | NA | L-Asparagine |
| Negative | 157 | 3.138 | 132.0299 | Aspartic Acid | Aspartate; LC-ESI-ITFT; MS2; CE 85.0 eV; [M-H]- | L-Aspartic acid |
| Negative | 160 | 5.48 | 133.0141 | Malic acid PGC | D-(+)-Malic acid | L-Malic acid |
| Negative | 167 | 4.275 | 135.0291 | Threonic Acid | NA | NA |
| Negative | 173 | 13.505 | 137.0237 | Salicylic acid | P-HYDROXYBENZOIC ACID | NA |
| Negative | 186 | 3.161 | 145.0616 | Glutamine | Glutamine; LC-ESI-ITFT; MS2; CE 75.0 eV; [M-H]- | L-Glutamine |
| Negative | 189 | 3.56 | 146.0455 | Glutamic Acid | L-Glutamic acid; LC-ESI-QTOF; MS2; CE | L-Glutamic acid |
| Negative | 194 | 4.886 | 149.0449 | Unknown | NA | D-Xylose |
| Negative | 200 | 13.502 | 153.0187 | Unknown | 3,4-DIHYDROXYBENZOIC ACID | NA |
| Negative | 258 | 4.89 | 173.0088 | Unknown | DEHYDROASCORBIC ACID - 40.0 eV | NA |
| Negative | 260 | 5.592 | 173.0446 | Shikimic acid PGC | NA | Shikimic acid |
| Negative | 283 | 5.312 | 179.0552 | Unknown | NA | myo-Inositol |
| Negative | 287 | 6.866 | 180.0659 | Unknown | NA | L-Tyrosine |
| Negative | 291 | 3.415 | 181.0707 | Unknown | NA | Galactitol |
| Negative | 315 | 8.979 | 191.019 | Unknown | NA | Citric acid |
| Negative | 317 | 7.472 | 191.0193 | Unknown | NA | Citric acid |
| Negative | 318 | 6.392 | 191.0194 | Isocitric acid PGC | ISOCITRIC ACID | Citric acid |
| Negative | 322 | 5.178 | 191.056 | Unknown | D-(-)-Quinic acid; LC-ESI-QTOF; MS2; CE | Quinic acid |
| Negative | 336 | 4.98 | 195.0526 | Unknown | 1726059 - 40.0 eV | Galactonic acid |
| Negative | 370 | 10.741 | 206.0819 | Unknown | NA | Afalanina |

| | | | | | | |
|---|---|---|---|---|---|---|
| Negative | 407 | 11.677 | 219.0766 | Unknown | 5-HYDROXY-TRYPTOPHAN | NA |
| Negative | 764 | 5.097 | 341.1107 | Dihexose (Maltose/Sucrose) | SUCROSE | Sucrose |
| Negative | 837 | 9.828 | 371.0973 | Unknown | NA | COCONUT:(CNP0115847 CNP0131848);PubChem:(45783079 51693489 51693491 101793099 133556366);SuperNatural:(SN00033299 SN00033300);ZINC bio:(ZINC35457686 ZINC35457690);additional;Training Set |
| Negative | 838 | 10.416 | 371.0978 | Unknown | (2S,3S,4S,5R,6R)-6-(3-benzoyloxy-2-hydroxypropoxy)-3,4,5-trihydroxyoxane-2-carboxylic acid | COCONUT:(CNP0115847 CNP0131848);PubChem:(45783079 51693489 51693491 101793099 133556366);SuperNatural:(SN00033299 SN00033300);ZINC bio:(ZINC35457686 ZINC35457690);additional;Training Set |
| Negative | 881 | 13.386 | 385.0921 | Unknown | 8-5'-Benzofuran-diferulic acid | NA |
| Negative | 891 | 4.976 | 387.1164 | Unknown | alpha,alpha-Trehalose - 40.0 eV | NA |
| Negative | 915 | 13.199 | 393.1745 | Unknown | NA | PubChem:(134883503) |
| Negative | 928 | 9.758 | 399.0923 | Unknown | NA | COCONUT:(CNP0170379);PubChem:(51136464);SuperNatural:(SN00039051);ZINC bio:(ZINC49180912);additional;Training Set |
| Negative | 935 | 8.486 | 403.0902 | Unknown | NA | PubChem:(46222441) |
| Negative | 960 | 11.888 | 413.145 | Unknown | NA | COCONUT:(CNP0159349 CNP0218833);Natural Products:(UNPD202200);PubChem:(38363079 38363084 45360328 44715457 125416071 125416072 125416073 125416074);SuperNatural:(SN00032625 SN00030299 SN00032624 SN00030300 SN00032623 SN00032622);ZINC bio:(ZINC31169353 ZINC31169357 ZINC35454681 ZINC35454685 ZINC35454688 ZINC35454690);Training Set |
| Negative | 961 | 11.055 | 413.1454 | Unknown | NA | COCONUT:(CNP0159349 CNP0218833);Natural Products:(UNPD202200);PubChem:(38363079 38363084 45360328 44715457 125416071 125416072 125416073 125416074);SuperNatural:(SN00032625 |

| | | | | | | |
|---|---|---|---|---|---|---|
| | | | | | | SN00030299 SN00032624 SN00030300 SN00032623 SN00032622);ZINC bio:(ZINC31169353 ZINC31169357 ZINC35454681 ZINC35454685 ZINC35454688 ZINC35454690);Training Set |
| Negative | 1033 | 9.105 | 435.0597 | Unknown | NA | PubChem:(102531776) |
| Negative | 1048 | 13.76 | 439.2544 | Unknown | NA | PubChem:(52919663 75954071) |
| Negative | 1103 | 9.189 | 460.1461 | Unknown | NA | PubChem:(53359647) |
| Negative | 1263 | 12.06 | 535.1809 | Unknown | NA | PubChem:(89189281) |
| Negative | 1271 | 5.322 | 539.139 | Unknown | RAFFINOSE CollisionEnergy:102040 | NA |
| Negative | 1305 | 12.951 | 557.1855 | Unknown | NA | 3-[[4,6-bis[bis(2-carboxyethyl)amino]-1,3,5-triazin-2-yl]-(2-carboxyethyl)amino]propanoic acid |
| Negative | 1364 | 10.413 | 613.2341 | Unknown | NA | PubChem:(130251683) |
| Negative | 1449 | 13.876 | 677.3527 | Unknown | NA | COCONUT:(CNP0090331) |
| Negative | 1464 | 15.17 | 686.4767 | Unknown | PE(16:1_16:1) - (2-aminoethoxy)[2,3-di[hexadec-9-enoyloxy]propoxy]phosphinic acid | NA |
| Negative | 1471 | 15.264 | 688.4916 | Unknown | PE(16:0/16:1); [M-H]- C37H71N1O8P1 | NA |
| Negative | 1499 | 14.523 | 707.4866 | Unknown | PG(15:0/16:0); [M-H]- C37H72O10P1 | NA |
| Negative | 1508 | 15.071 | 711.4915 | Unknown | NA | benzyl N-[(2S)-6-amino-1-[3-[4-[3-[[(2S)-6-amino-2-(phenylmethoxycarbonylamino)hexanoyl]amino]propylamino]butylamino]propylamino]hexan-2-yl]carbamate |
| Negative | 1514 | 15.299 | 714.5074 | Unknown | PE(16:1/18:1); [M-H]- C39H73N1O8P1 | NA |
| Negative | 1523 | 14.496 | 719.4865 | Unknown | Massbank:LQB00238 PG 32:1 | NA |
| Negative | 1560 | 14.5 | 745.5029 | Unknown | PG(16:0/18:2); [M-H]- C40H74O10P1 | NA |
| Negative | 1565 | 14.531 | 747.5177 | Unknown | Massbank:LQB00241 PG 34:1 | NA |
| Negative | 1606 | 14.503 | 771.517 | Unknown | PG(18:1/18:2); [M-H]- C42H76O10P1 | NA |
| Negative | 1607 | 14.536 | 773.5337 | Unknown | Massbank:LQB00261 PG 36:2 | NA |
| Negative | 1678 | 14.2 | 833.5185 | Unknown | Massbank:LQB00679 PI 34:2 | NA |
| Negative | 1781 | 13.508 | 1019.505 | Unknown | NA | PubChem:(10581887) |
| Positive | 15 | 3.022 | 58.0649 | Unknown | NA | 2-Methylaziridine |
| Positive | 102 | 6.954 | 86.09585 | Unknown | Piperidine; LC-ESI-QTOF; MS2; CE | Piperidine |

| | | | | | | |
|---|---|---|---|---|---|---|
| Positive | 162 | 3.5 | 104.1073 | Unknown | Choline; LC-ESI-QTOF; MS2; CE | NA |
| Positive | 201 | 3.171 | 116.0708 | Proline PGC | Proline; LC-ESI-ITFT; MS2; CE 85.0 eV; [M+H]+ | NA |
| Positive | 213 | 3.155 | 118.0865 | Unknown | Betaine; LC-ESI-QTOF; MS2; CE | NA |
| Positive | 241 | 3.133 | 130.0499 | Pyroglutamic acid | L-5-Oxoproline | Pyroglutamic acid |
| Positive | 254 | 2.909 | 133.0612 | Asparagine | NA | agedoite |
| Positive | 260 | 3.142 | 134.0454 | Unknown | Aspartic acid; LC-ESI-QTOF; MS2; CE | L-Aspartic acid |
| Positive | 263 | 13.345 | 134.0606 | Unknown | Massbank:TUE00032 5-Methylbenzotriazole | Acetaminophen |
| Positive | 267 | 6.513 | 136.0755 | Unknown | NA | 2-Phenylacetamide |
| Positive | 274 | 4.966 | 138.0549 | Unknown | NA | 2-Aminobenzoic acid |
| Positive | 275 | 10.436 | 138.0917 | Unknown | NA | Tyramine |
| Positive | 302 | 3.13 | 147.0765 | Glutamine | L-Glutamine; LC-ESI-QTOF; MS2; CE | L-Glutamine |
| Positive | 303 | 5.083 | 147.0771 | Unknown | NA | L-Glutamine |
| Positive | 305 | 3.51 | 148.0609 | Glutamic acid | L-Glutamic acid; LC-ESI-QTOF; MS2; CE | L-Glutamic acid |
| Positive | 316 | 10.053 | 152.0707 | Unknown | NA | Acetaminophen |
| Positive | 329 | 5.027 | 156.0766 | Histidine | L-Histidine; LC-ESI-QTOF; MS2; CE | L-Histidine |
| Positive | 347 | 10.911 | 161.06 | Unknown | NA | 6-Methylcoumarin |
| Positive | 360 | 6.556 | 165.0542 | Unknown | NA | 4-Hydroxycinnamic acid |
| Positive | 363 | 10.503 | 166.0867 | Unknown | NA | Sabiden |
| Positive | 364 | 9.565 | 166.0867 | Unknown | NA | L-Phenylalanine |
| Positive | 382 | 4.658 | 175.1079 | Arginine | NA | NA |
| Positive | 384 | 9.14 | 176.07 | Unknown | NA | Indoleacetic acid |
| Positive | 410 | 6.514 | 182.0812 | Unknown | Tyrosine; LC-ESI-ITFT; MS2; CE 45.0 eV; [M+H]+ | L-Tyrosine |
| Positive | 418 | 2.862 | 184.0731 | Unknown | NA | phosphocholine |
| Positive | 495 | 13.153 | 205.0971 | Unknown | L-Tryptophan; LC-ESI-QTOF; MS2; CE | NA |
| Positive | 530 | 8.93 | 220.1189 | Unknown | pantothenic acid CollisionEnergy:205060 | Pantothenic acid |
| Positive | 534 | 11.436 | 221.092 | Unknown | NA | 5-Hydroxy-L-tryptophan |
| Positive | 538 | 4.987 | 222.0976 | Unknown | NA | N-Acetylgalactosamine |
| Positive | 664 | 3.472 | 258.1111 | Unknown | ReSpect:PM018116 sn-Glycero-3-phosphocholine | Gliatilin (TN) |
| Positive | 923 | 4.608 | 336.1408 | Unknown | NA | Asn-agam |

| | | | | | | |
|---|---|---|---|---|---|---|
| Positive | 990 | 4.806 | 360.1506 | Unknown | Spectral Match to D-(+)-Trehalose from NIST14 | NA |
| Positive | 1000 | 4.916 | 365.1062 | Dihexose (Maltose/Sucrose) | NA | NA |
| Positive | 1053 | 9.272 | 384.1148 | Unknown | NA | Succinoadenosine |
| Positive | 1149 | 9.065 | 422.1119 | Unknown | NA | PubChem:(101229455) |
| Positive | 1246 | 12.708 | 454.229 | Unknown | NA | PubChem:(117619884) |
| Positive | 1294 | 14.725 | 471.3955 | Unknown | NA | N-Arachidoyl-5-hydroxytryptamine |
| Positive | 1430 | 5.142 | 522.2022 | Unknown | Melezitose | NA |
| Positive | 1831 | 15.087 | 688.493 | Unknown | Spectral Match to 1,2-Dipalmitoleoyl-sn-glycero-3-phosphoethanolamine from NIST14 | NA |
| Positive | 1913 | 15.191 | 716.523 | Unknown | Spectral Match to 2-Linoleoyl-1-palmitoyl-sn-glycero-3-phosphoethanolamine from NIST14 | NA |
| Positive | 1997 | 14.562 | 752.5299 | Unknown | NA | Dien-microgonotropen-c |
| Positive | 2010 | 15.27 | 758.5715 | Unknown | PC(16:1/18:1); [M+H]+ C42H81N1O8P1 | NA |
| Positive | 2045 | 15.208 | 782.5713 | Unknown | Spectral Match to 1,2-Dilinoleoyl-sn-glycero-3-phosphocholine from NIST14 | NA |
| Positive | 2050 | 15.363 | 786.6017 | Unknown | PC(18:1/18:1); [M+H]+ C44H85N1O8P1 | PC(18:1(9Z)/18:1(9Z)) |
| Positive | 2061 | 14.556 | 797.5313 | Unknown | Spectral Match to 1,2-Dioleoyl-sn-glycero-3-phospho-rac-1-glycerol from NIST14 | NA |
| Positive | 2099 | 13.156 | 841.423 | Unknown | NA | COCONUT:(CNP0326190);Natural Products:(UNPD89153);PubChem:(24066892 102464927);SuperNatural:(SN00293878) |
| Positive | 2102 | 14.725 | 842.6728 | Unknown | NA | COCONUT:(CNP0277004);Natural Products:(UNPD147896);PubChem:(23427479);SuperNatural:(SN00337084) |

**Table S11:** Compounds identified within the black grass experimental set following second extraction. For identification level 1, the annotation" w/o MS2" indicates there was not confident spectral match and that the results can not be fully trusted unless corroborated by another source. If identification level 2 conflicts with these level 1 annotations, confidence level 2 should be used.

| Ionization Mode | Alignment ID | Average Rt(min) | Average Mz | Level 1 Identification | Level 2a Identification | Level 2b Identification |
|---|---|---|---|---|---|---|
| Negative | 20 | 3.771 | 105.0196 | Unknown | D-(+)-Glyceric acid; LC-ESI-QTOF; MS2; CE | NA |
| Negative | 28 | 3.364 | 115.0032 | w/o MS2:Fumaric acid | NA | Fumaric acid |
| Negative | 35 | 3.086 | 117.0188 | Succinic acid | Succinic acid; LC-ESI-QTOF; MS2; CE | Succinic acid |
| Negative | 45 | 2.949 | 131.0458 | Asparagine | NA | NA |
| Negative | 47 | 3.193 | 132.0311 | Aspartic Acid | Aspartate; LC-ESI-ITFT; MS2; CE 85.0 eV; [M-H]- | NA |
| Negative | 50 | 5.307 | 133.0138 | Malic acid PGC | D-(+)-Malic acid; LC-ESI-QTOF; MS2; CE | L-Malic acid |
| Negative | 54 | 4.319 | 135.029 | Threonic Acid | NA | NA |
| Negative | 55 | 13.62 | 137.0249 | Salicylic acid | NA | 4-Hydroxybenzoic acid |
| Negative | 59 | 3.235 | 145.0622 | Glutamine | Glutamine; LC-ESI-ITFT; MS2; CE 80.0 eV; [M-H]- | L-Glutamine |
| Negative | 62 | 3.591 | 146.0466 | Glutamic Acid | L-Glutamic acid | NA |
| Negative | 88 | 5.352 | 173.0462 | Shikimic acid PGC | (-)-Shikimic acid | Shikimic acid |
| Negative | 103 | 5.103 | 179.0562 | Unknown | NA | D-Glucose |
| Negative | 125 | 3.149 | 191.019 | Citric acid | CITRIC ACID | Citric acid |
| Negative | 128 | 6.112 | 191.0208 | Isocitric acid PGC | ISOCITRIC ACID | Citric acid |
| Negative | 131 | 4.985 | 191.0559 | Unknown | D-(-)-Quinic acid; LC-ESI-QTOF; MS2; CE | Quinic acid |
| Negative | 143 | 10.801 | 197.0445 | Syringic acid | NA | NA |
| Negative | 183 | 4.91 | 220.0819 | w/o MS2:N-acetyl-D-mannosamine | NA | N-Acetylgalactosamine |
| Negative | 446 | 4.864 | 341.1099 | Dihexose (Maltose/Sucrose) | Sucrose | Sucrose |
| Negative | 498 | 5.337 | 365.0437 | Unknown | NA | [1,2,3-trihydroxy-1-[(3R,4S,5R,6R)-3,4,5-trihydroxy-6-(hydroxymethyl)oxan-2-yl]propoxy] hydrogen sulfate |

| | | | | | | |
|---|---|---|---|---|---|---|
| Negative | 513 | 10.546 | 371.0975 | Unknown | NA | COCONUT:(CNP0115847 CNP0131848);PubChem:(45783079 51693489 51693491 101793099 133556366);SuperNatural:(SN00033299 SN00033300);ZINC bio:(ZINC35457686 ZINC35457690);additional;Training Set |
| Negative | 594 | 8.631 | 409.0444 | Unknown | NA | 2''-Deoxy-5''-adenylyl imidodiphosphate |
| Negative | 644 | 9.877 | 431.1203 | Unknown | NA | Licoagroside B |
| Negative | 771 | 12.986 | 489.1979 | Unknown | NA | PubChem:(135430392) |
| Negative | 788 | 12.812 | 498.1826 | Unknown | NA | PubChem:(117768116) |
| Negative | 802 | 9.467 | 505.1567 | Unknown | NA | KEGG Mine |
| Negative | 834 | 10.148 | 520.1682 | Unknown | NA | PubChem:(10075389) |
| Negative | 969 | 5.261 | 637.184 | Unknown | NA | PubChem:(54564545) |
| Negative | 1008 | 4.86 | 683.2268 | Unknown | PALATINOSE | NA |
| Negative | 1024 | 5.056 | 695.2246 | Unknown | NA | PubChem:(91847146) |
| Negative | 1026 | 12.706 | 697.2341 | Unknown | NA | PubChem:(101607589) |
| Negative | 1102 | 14.656 | 747.5154 | Unknown | [2,3-dihydroxypropoxy][3-(hexadecanoyloxy)-2-[octadec-9-enoyloxy]propoxy]phosphinic acid | NA |
| Negative | 1148 | 14.185 | 785.4107 | Unknown | NA | PubChem:(132018988) |
| Negative | 1151 | 14.721 | 787.547 | Unknown | NA | PubChem:(88641774) |
| Positive | 21 | 10.938 | 60.04413 | Unknown | NA | Acetamide |
| Positive | 26 | 2.959 | 60.08095 | Unknown | NA | Trimethylamine |
| Positive | 101 | 7.199 | 86.0963 | Unknown | Piperidine; LC-ESI-QTOF; MS2; CE | Piperidine |
| Positive | 131 | 4.87 | 97.02834 | Unknown | NA | 2-Furancarboxaldehyde |
| Positive | 161 | 3.569 | 104.1072 | Unknown | Choline; LC-ESI-QTOF; MS2; CE | NA |
| Positive | 199 | 3.247 | 116.0702 | Proline PGC | Proline; LC-ESI-ITFT; MS2; CE 85.0 eV; [M+H]+ | L-Proline |
| Positive | 231 | 9.123 | 127.0391 | Unknown | NA | 1,2,3-Trihydroxybenzene |
| Positive | 239 | 4.629 | 130.0499 | Unknown | NA | Pidolate |
| Positive | 241 | 3.221 | 130.0506 | Pyroglutamic acid | L-5-Oxoproline; LC-ESI-QTOF; MS2; CE | Pyroglutamic acid |
| Positive | 257 | 2.976 | 133.0608 | Asparagine | NA | agedoite |
| Positive | 262 | 3.231 | 134.0448 | Unknown | NA | aspartate |

| | | | | | | |
|---|---|---|---|---|---|---|
| Positive | 264 | 13.451 | 134.0602 | Unknown | NA | Acetaminophen |
| Positive | 276 | 4.926 | 138.0548 | Unknown | NA | 2-Aminobenzoic acid |
| Positive | 293 | 2.955 | 143.0815 | Unknown | NA | 3-imidazol-1-ylpropane-1,2-diol |
| Positive | 296 | 5.397 | 144.0658 | Unknown | NA | Adipo-2,6-lactam |
| Positive | 309 | 6.724 | 147.0436 | Unknown | NA | Coumarin |
| Positive | 311 | 3.218 | 147.077 | Glutamine | L-Glutamine; LC-ESI-QTOF; MS2; CE | L-Glutamine |
| Positive | 313 | 3.587 | 148.061 | Glutamic acid | L-Glutamic acid; LC-ESI-QTOF; MS2; CE | L-Glutamic acid |
| Positive | 325 | 13.451 | 152.0709 | Unknown | NA | zlchem 1306 |
| Positive | 340 | 5.166 | 156.0769 | Histidine | Histidine; LC-ESI-QTOF; MS2; CE | histidin |
| Positive | 376 | 6.714 | 165.0541 | Unknown | NA | 4-Hydroxycinnamic acid |
| Positive | 379 | 10.654 | 166.086 | Unknown | NA | Sabiden |
| Positive | 380 | 9.643 | 166.0866 | Unknown | NA | L-Phenylalanine |
| Positive | 389 | 9.108 | 170.0814 | Unknown | NA | Adermine |
| Positive | 408 | 4.634 | 175.1081 | w/o MS2:Arginine | NA | N-Acetylornithine |
| Positive | 440 | 6.711 | 182.0809 | Unknown | TYROSINE | tyrosine |
| Positive | 551 | 9.024 | 220.1178 | Unknown | Massbank:LU087003 Pantothenate\|Pantothenic acid\|3-[[(2R)-2,4-dihydroxy-3,3-dimethylbutanoyl]amino]propanoic acid | vitamin B5 |
| Positive | 559 | 4.99 | 222.0974 | Unknown | NA | N-Acetylgalactosamine |
| Positive | 645 | 12.593 | 244.1657 | Unknown | NA | prolyllysine |
| Positive | 692 | 3.566 | 258.1108 | Unknown | sn-Glycero-3-phosphocholine; LC-ESI-QTOF; MS2; CE | NA |
| Positive | 716 | 10.063 | 263.1602 | Unknown | NA | Poly(ethylene glycol) ethyl ether methacrylate |
| Positive | 734 | 8.552 | 268.1044 | Unknown | NA | Adenosine |
| Positive | 745 | 10.972 | 270.1333 | Unknown | NA | COCONUT:(CNP0352172);Natural Products:(UNPD6299);PubChem:(57478618);SuperNatural:(SN00281451) |
| Positive | 757 | 13.999 | 272.2216 | Unknown | NA | acylglycine c:13 |
| Positive | 813 | 9.13 | 289.0923 | Unknown | 2-methyl-3-[(2S,3R,4S,5S,6R)-3,4,5-trihydroxy-6-(hydroxymethyl)oxan-2-yl]oxypyran-4-one | maltol beta-D-glucopyranoside |
| Positive | 894 | 8.727 | 313.0855 | Unknown | NA | Acetaminophen-2-mercapturate |

| | | | | | | |
|---|---|---|---|---|---|---|
| Positive | 1003 | 10.649 | 352.2231 | Unknown | NA | PubChem:(129847081);COCONUT:(CNP0112721) |
| Positive | 1022 | 4.838 | 360.1508 | Unknown | Spectral Match to Palatinose from NIST14 | NA |
| Positive | 1032 | 4.873 | 365.106 | Dihexose (Maltose/Sucrose) | NA | NA |
| Positive | 1187 | 14.97 | 427.3861 | Unknown | NA | Lanosterin |
| Positive | 1202 | 12.715 | 433.1129 | Unknown | Spectral Match to Isovitexin from NIST14 | NA |
| Positive | 1321 | 10.161 | 476.1763 | Unknown | NA | (2R)-2-{[6-O-(beta-D-glucopyranosyl)-beta-D-glucopyranosyl]oxy}-2-phenylethanamide |
| Positive | 1813 | 15.161 | 688.4927 | Unknown | Spectral Match to 1,2-Dipalmitoleoyl-sn-glycero-3-phosphoethanolamine from NIST14 | NA |
| Positive | 1820 | 15.238 | 690.5082 | Unknown | PE-DAG (16:0/16:1) | NA |
| Positive | 1956 | 15.359 | 758.571 | Unknown | Spectral Match to 1-Hexadecanoyl-2-octadecadienoyl-sn-glycero-3-phosphocholine from NIST14 | NA |
| Positive | 1986 | 15.297 | 782.5698 | Unknown | Spectral Match to 1,2-Dilinoleoyl-sn-glycero-3-phosphocholine from NIST14 | NA |

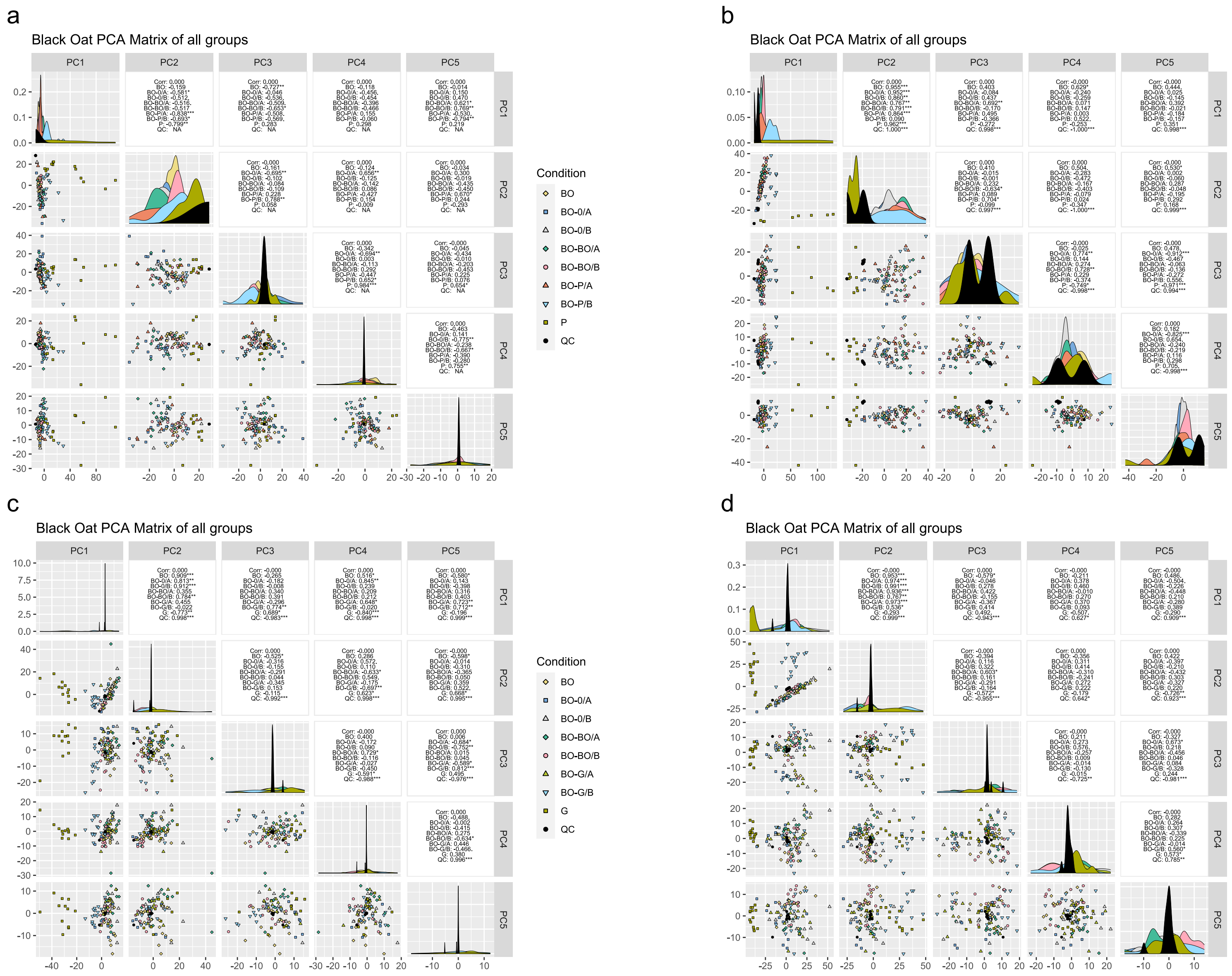
a
Black Oat PCA Matrix of all groups
PC1
PC2
PC3
PC4
PC5
Condition
BO
BO-0/A
BO-0/B
BO-BO/A
BO-BO/B
BO-P/A
BO-P/B
P
QC
b
Black Oat PCA Matrix of all groups
Condition
BO
BO-0/A
BO-0/B
BO-BO/A
BO-BO/B
BO-P/B
BO-P/A
P
QC
c
Black Oat PCA Matrix of all groups
Condition
BO
BO-0/A
BO-0/B
BO-BO/A
BO-BO/B
BO-G/A
BO-G/B
G
QC
d
Black Oat PCA Matrix of all groups
Condition
BO
BO-0/A
BO-BO/B
BO-0/B
BO-BO/A
BO-G/A
BO-G/B
G
QC

**Figure S1:** PCA matrix for the first 5 components of redroot pigweed experiment aqueous extracts (a) and methanolic extracts (b) and blackgrass experiment aqueous extracts (c) and methanolic extracts (d) comparing the 3 A compartments for each experiment.

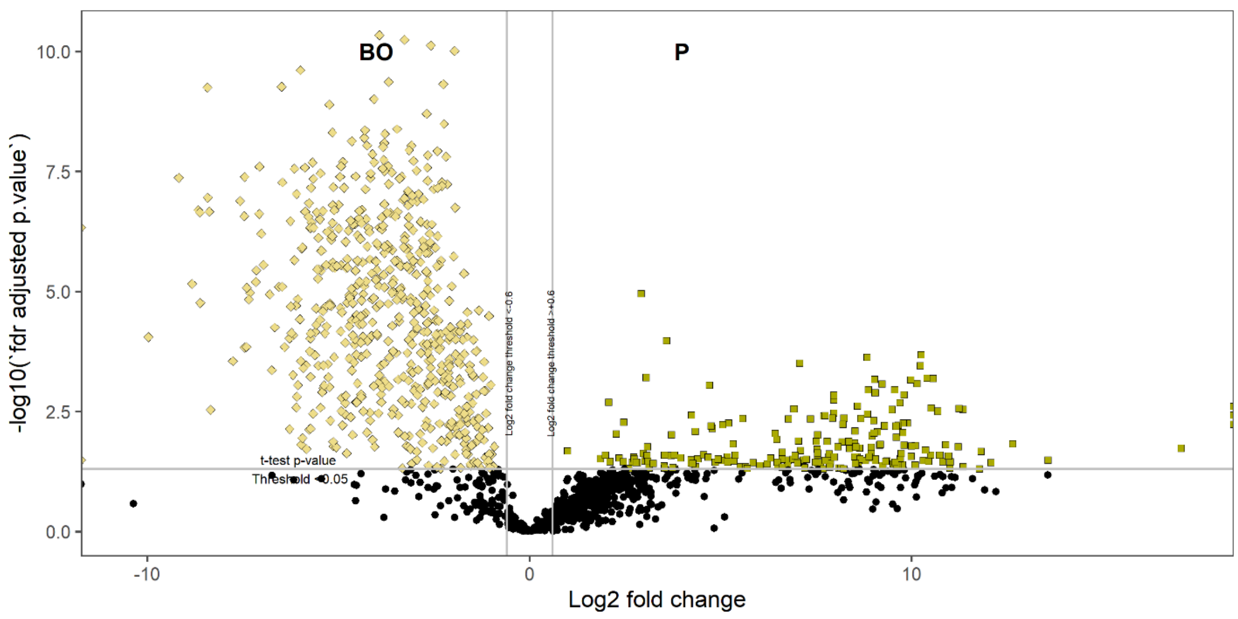

**Figure S2:** Compounds upregulated in P when compared to BO for the redroot P experimental set

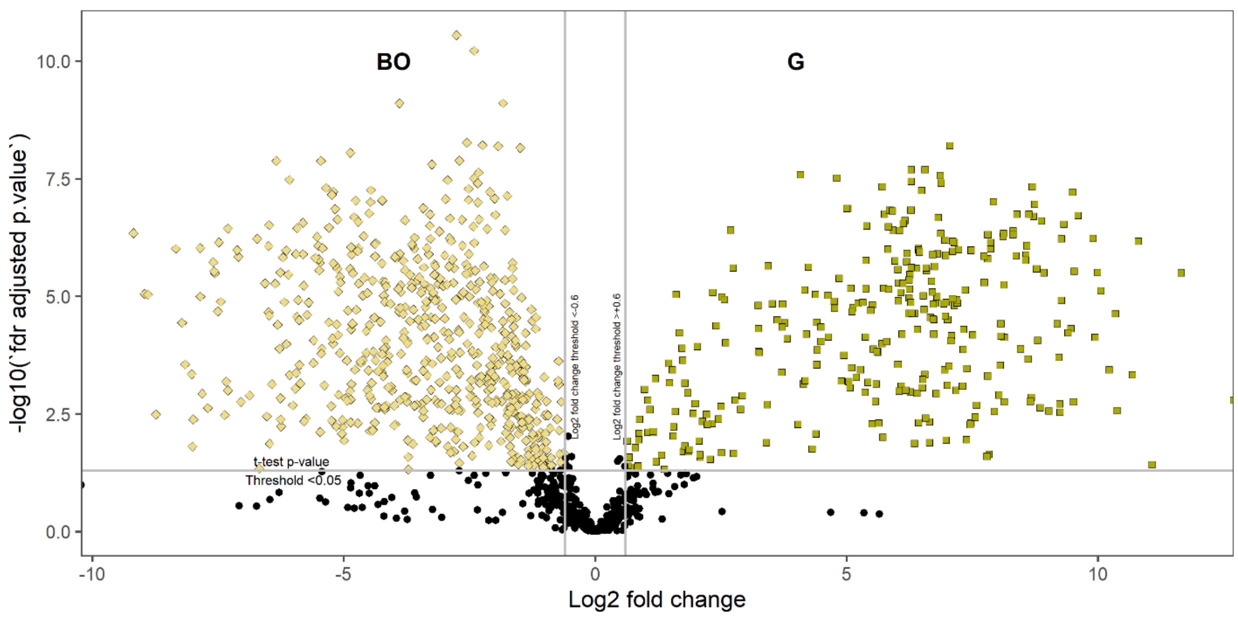

**Figure S3:** Compounds upregulated in G when compared to BO for the G experimental set

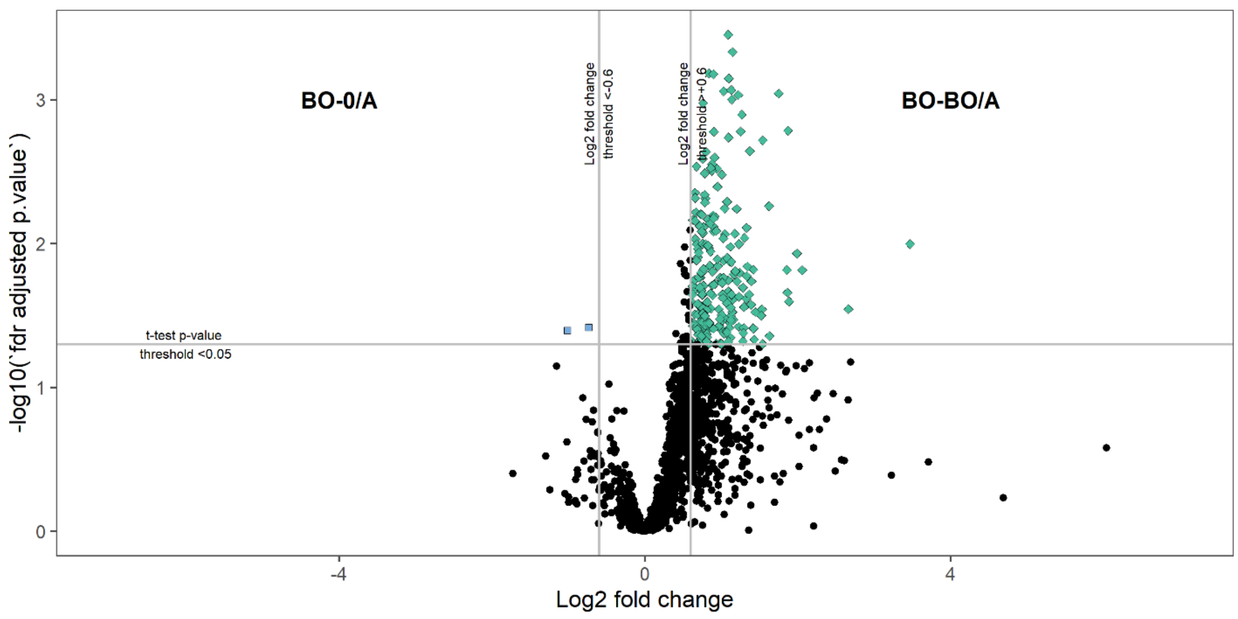

**Figure S4:** Compounds upregulated by BO-BO/A when compared to BO-0/A for the P experimental set

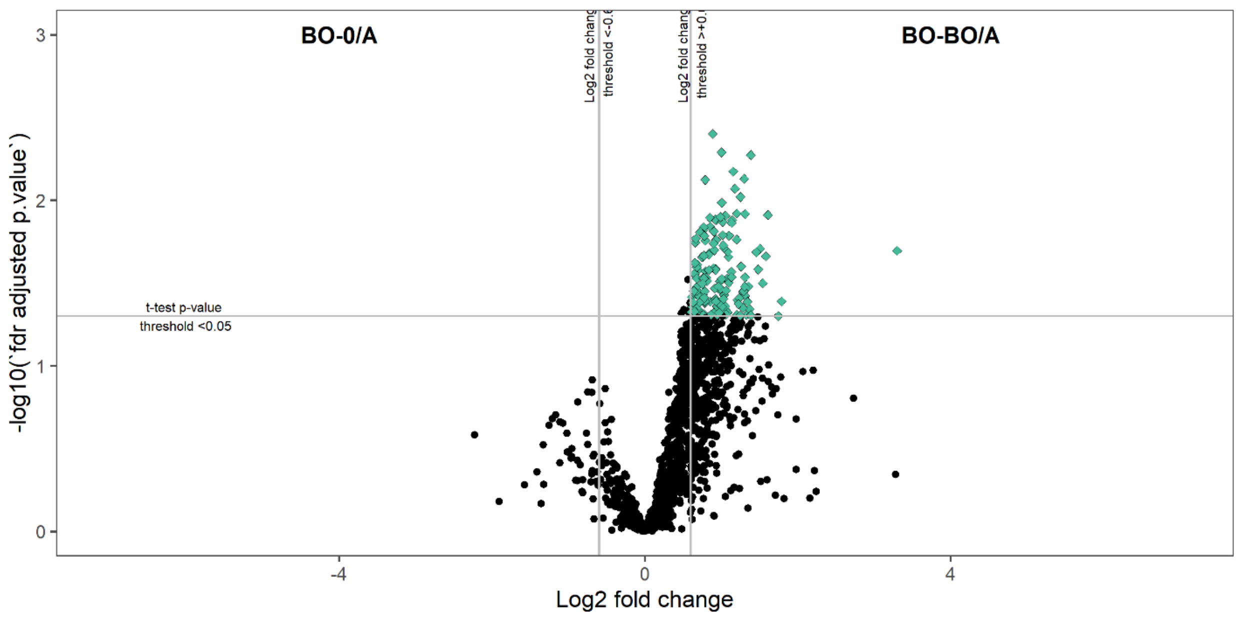

**Figure S5:** Compounds upregulated by BO-BO/A when compared to BO-0/A for the G experimental set

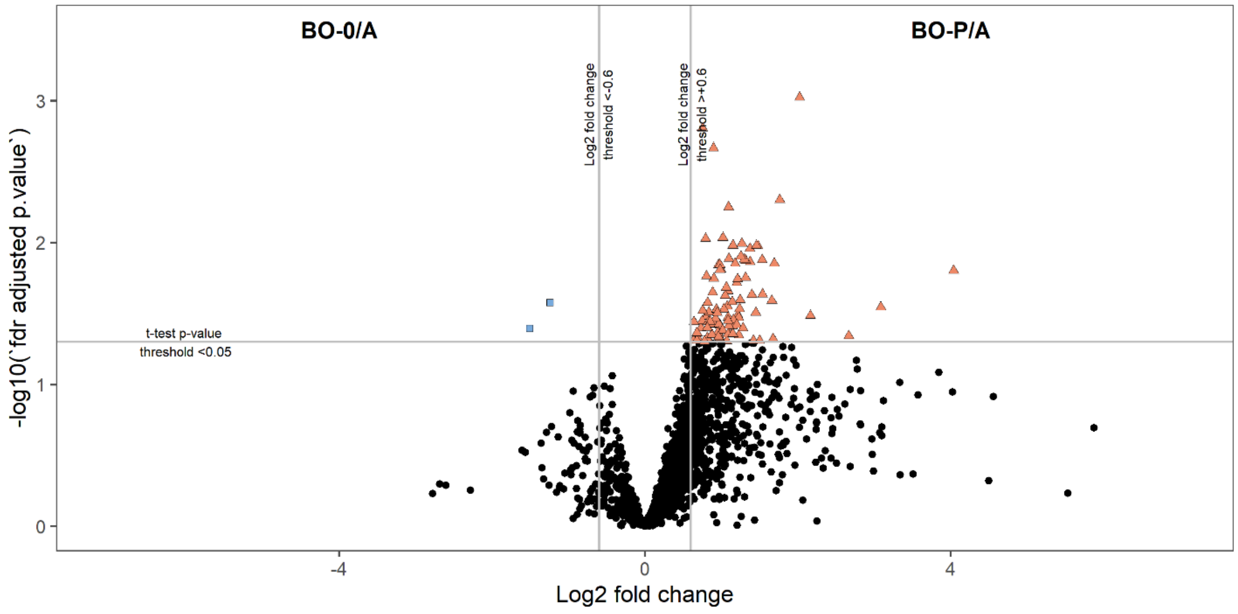

**Figure S6:** Compounds upregulated by BO-P/A when compared to BO-0/A for the P experimental set

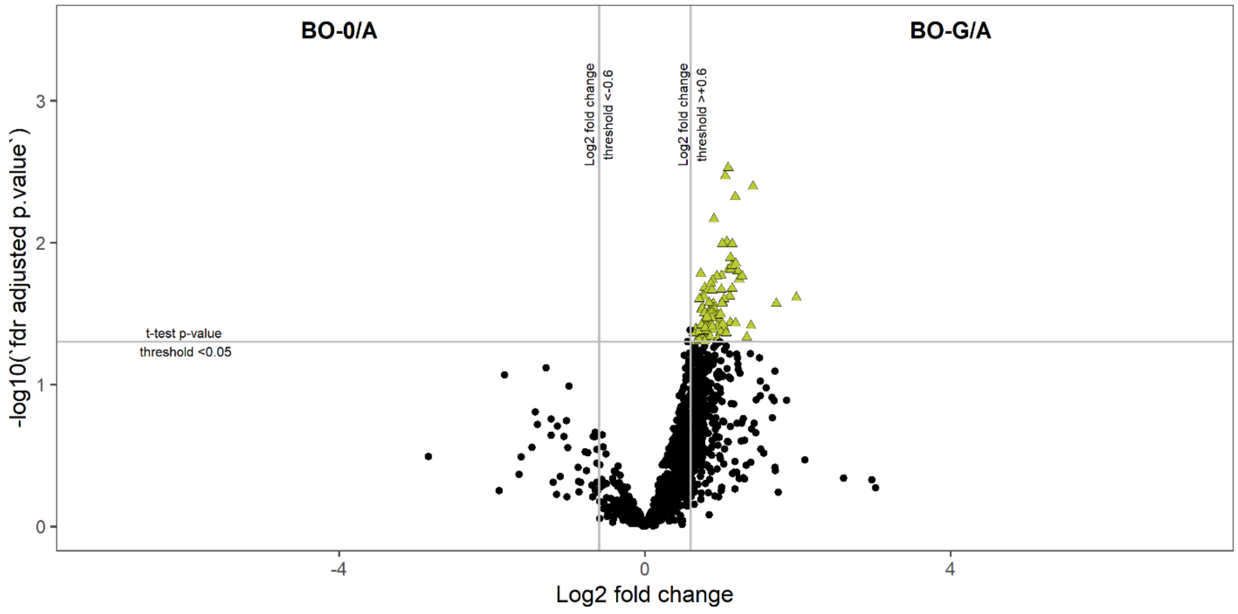

**Figure S7:** Compounds upregulated by BO-G/A when compared to BO-0/A for the G experimental set

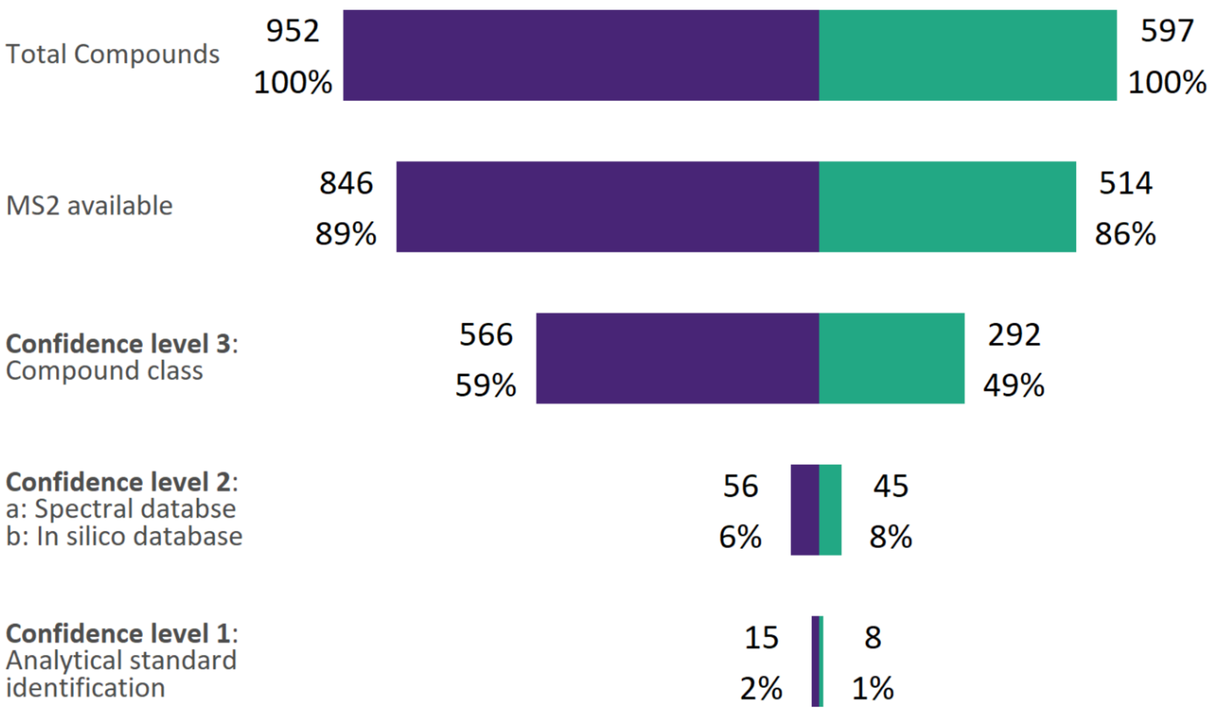

**Figure S9:** Compounds were identified to different levels of confidence based on the Schymanski scale for the P experimental set from the second methanolic re-exudation extract. For confidence levels 1-3, a fragmentation pattern (MS2) is necessary for identification. Identification level 3, compound class identification, was annotated using the CANOPUS application of the SIRIUS open-source software. Identification level 2a was annotated with MS FINDER and GNPS using all available open-source libraries. 2b was annotated with MS FINDER and SIRIUS CSI: FingerID. Confidence level 1 was annotated with an in-house database generated from a set of authentic analytical standards of compounds known to be exudated by the roots of crops. Negative ionization mode data is shown in purple and positive ionization mode is teal.

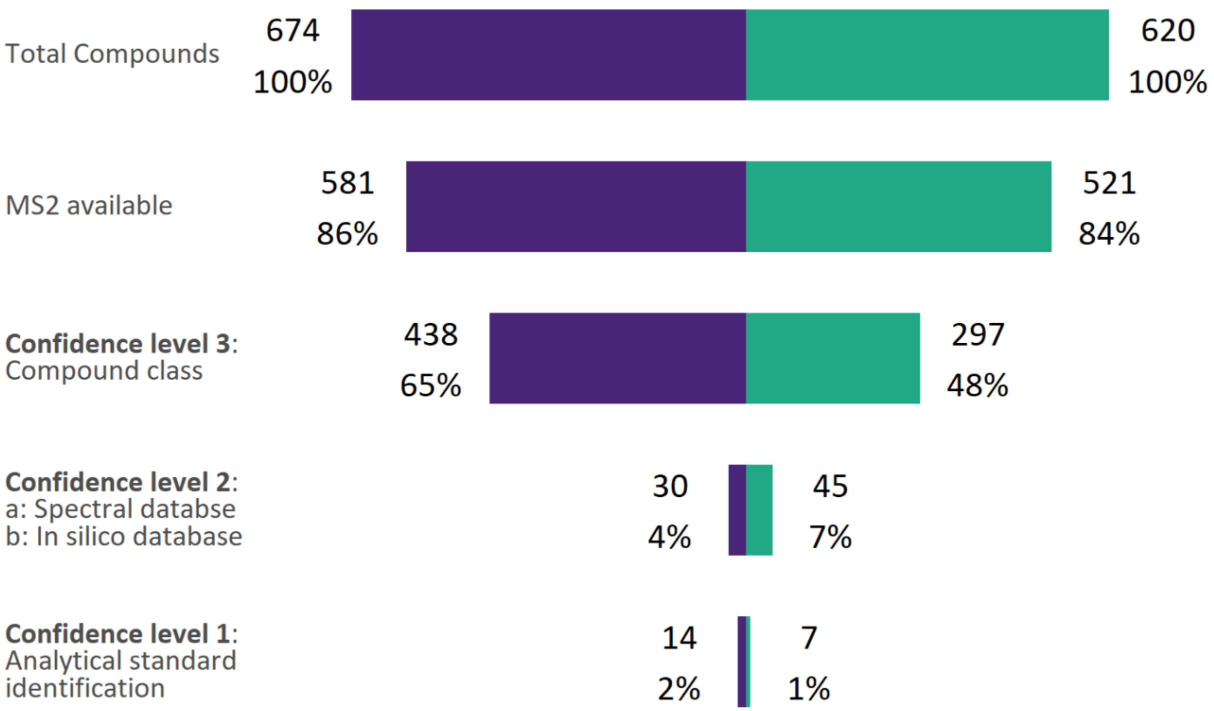

**Figure S8:** Compounds were identified to different levels of confidence based on the Schymanski scale for the G experimental set from the second methanolic re-exudation extract. For confidence levels 1-3, a fragmentation pattern (MS2) is necessary for identification. Identification level 3, compound class identification, was annotated using the CANOPUS application of the SIRIUS open-source software. Identification level 2a was annotated with MS FINDER and GNPS using all available open-source libraries. 2b was annotated with MS FINDER and SIRIUS CSI: FingerID. Confidence level 1 was annotated with an in-house database generated from a set of authentic analytical standards of compounds known to be exudated by the roots of crops. Negative ionization mode data is shown in purple and positive ionization mode is teal.

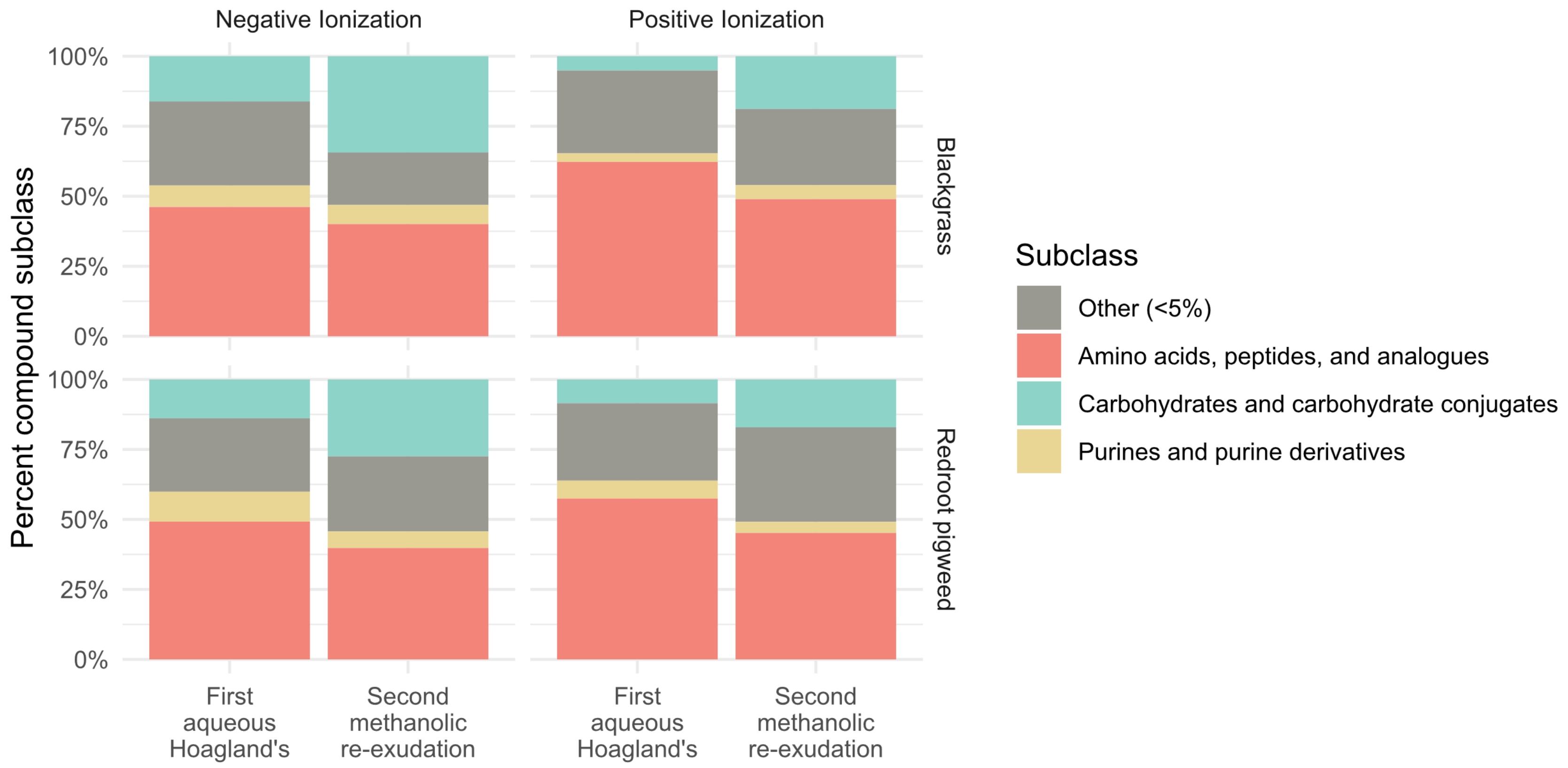

**Figure S10:** Analysis from the CANOPUS function in SIRIUS which identified all possible subclasses of compounds from all sample types exported from MS DIAL compares the first aqueous two-week extraction of Hoagland's solution to the second methanolic 24 hour re-exudation extraction. This is shown for both the P experimental set and the G experimental set in negative and positive ionization mode. Only compounds whose subclass could confidently be annotated were considered by applying a 90% probability score cutoff for the compound subclass identification. Any identification below this threshold were considered unknowns and were thus excluded. Compounds which were of a subclass which comprised less than 5% of the total number of identifiable compounds were binned into an "Other" category.
